# The genomic and cellular basis of biosynthetic innovation in rove beetles

**DOI:** 10.1101/2023.05.29.542378

**Authors:** Sheila A. Kitchen, Thomas H. Naragon, Adrian Brückner, Mark S. Ladinsky, Sofia A. Quinodoz, Jean M. Badroos, Joani W. Viliunas, Julian M. Wagner, David R. Miller, Mina Yousefelahiyeh, Igor A. Antoshechkin, K. Taro Eldredge, Stacy Pirro, Mitchell Guttman, Steven R. Davis, Matthew L. Aardema, Joseph Parker

**Author notes:** Present address: *Department of Marine Biology, Texas A&M University Galveston, TX, USA.

## Abstract

How evolution at the cellular level potentiates change at the macroevolutionary level is a major question in evolutionary biology. With >66,000 described species, rove beetles (Staphylinidae) comprise the largest metazoan family. Their exceptional radiation has been coupled to pervasive biosynthetic innovation whereby numerous lineages bear defensive glands with diverse chemistries. Here, we combine comparative genomic and single-cell transcriptomic data from across the largest rove beetle clade, Aleocharinae. We retrace the functional evolution of two novel secretory cell types that together comprise the tergal gland—a putative catalyst behind Aleocharinae’s megadiversity. We identify key genomic contingencies that were critical to the assembly of each cell type and their organ-level partnership in manufacturing the beetle’s defensive secretion. This process hinged on evolving a mechanism for regulated production of noxious benzoquinones that appears convergent with plant toxin release systems, and synthesis of an effective benzoquinone solvent that weaponized the total secretion. We show that this cooperative biosynthetic system arose at the Jurassic-Cretaceous boundary, and that following its establishment, both cell types underwent ∼150 million years of stasis, their chemistry and core molecular architecture maintained almost clade-wide as Aleocharinae radiated globally into tens of thousands of lineages. Despite this deep conservation, we show that the two cell types have acted as substrates for the emergence of adaptive, biochemical novelties—most dramatically in symbiotic lineages that have infiltrated social insect colonies and produce host behavior-manipulating secretions. Our findings uncover genomic and cell type evolutionary processes underlying the origin, functional conservation and evolvability of a chemical innovation in beetles.

## Introduction

Exceptional radiations are a recurring pattern across the tree of life (*1*). Pinpointing ancient genomic and cellular changes that proved to be innovations for the clades that habored them is a major challenge in evolutionary biology (*2*). The ∼400,000 described species of beetle (Coleoptera) (*3, 4*) are an archetype of evolutionary diversification that has long motivated biologists to consider the causes of species richness (*5–9*). The putative beetle key innovation is the elytron—the hardened forewing that shields the delicate flight wings—a structure that has enabled beetles to diversify in myriad terrestrial niches that are inaccessible to other winged insects (*6, 10–12*). Within the Coleoptera, however, lineage diversity is profoundly unbalanced, with ∼75% of species belonging to just ten of 190 extant beetle families. Efforts to understand the factors behind the exceptional richness of a small handful of beetle lineages have focused primarily on the Phytophaga, a megadiverse clade of ∼125,000 predominantly herbivorous species. The huge species richness of phytophagans has been posited to stem from their co-diversification with angiosperms during the Cretaceous and Cenozoic (*5, 13*), a phenomenon contingent on key metabolic changes that enabled these beetles to unlock recalcitrant trophic resources harbored by plants (*14–18*). The catalytic role played by angiosperm herbivory in beetle cladogenesis is broadly accepted, but leaves open the problem of explaining diversity in the remaining two thirds of the Coleoptera where herbivorous groups comprise only a minority of species (*19*). Amongst the greatest challenges is comprehending the diversity of Staphylinidae— the rove beetles—a clade of 66,459 predominantly predatory species that represents the largest family both in the Coleoptera and within the whole Metazoa (*20–22*).

Explanations for the extraordinary diversity of staphylinids include their short elytra and flexible abdomens that permit efficient movement through soil and litter microhabitats (*20, 23*). Coupled to this flexible body plan is a propensity for chemical innovation, whereby numerous lineages bear abdominal glands with unique, small molecule chemistries (*24, 25*). Species richness across the 34 staphylinid subfamilies is strongly skewed, however, with the largest being the Aleocharinae—a clade of 16,837 known species (*22*), speculated to be an order of magnitude more speciose (*26*) (**Fig. 1A**). Aleocharines are typically small-bodied (2–6 mm) predatory beetles and comprise arguably the most ecologically diverse clade in the Coleoptera. Aleocharines have radiated massively within Earth’s temperate and tropical zones, colonizing myriad terrestrial niches including litter, soil, saproxylic and subcortical microhabitats. The group has evolved to exploit fungi, carrion and vascular plants, and reached environmental extremes in caves, deep soil, intertidal regions and transiently submerged coral reefs (*27–30*). Within these diverse habitats, aleocharines exhibit widespread ecological and trophic specialization, manifested in clades of ectoparasitoids, vertebrate commensals and social insect symbionts, as well as numerous lineages that have shifted to fungus-, dead wood-, plant- and pollen-feeding.

**Figure 1.**
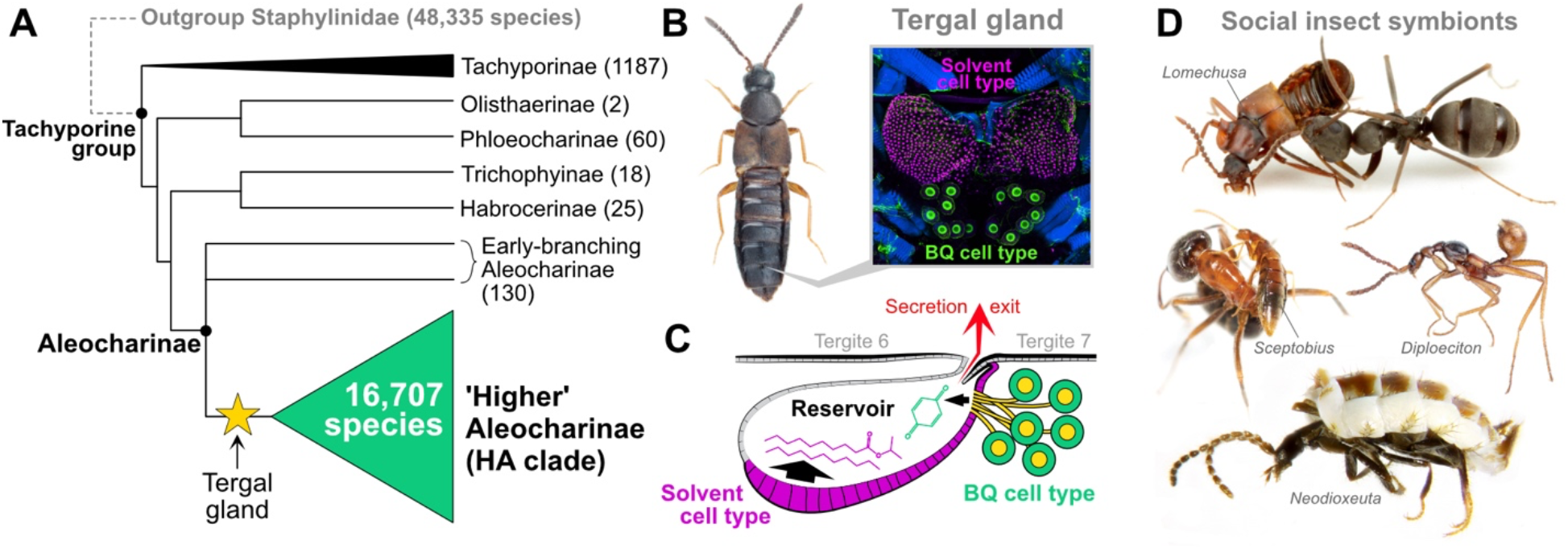
Aleocharine rove beetles. **(A)** Cladogram of tachyporine-group Staphylinidae (*47, 49*) showing major radiation of Higher Aleocharinae. Numbers in parentheses indicate number of extant species. **(B)** Example of a free-living aleocharine (*Atheta* sp.) with confocal image of tergal gland showing position on body. The tergal gland sits dorsally between tergites 6 and 7 and comprises two cell types, the solvent cells (magenta) and BQ cells (green). **(C)** Cartoon of tergal gland showing solvent and BQ cells secreting into a common reservoir that ejects the defensive cocktail between tergites. **(D)** Examples of aleocharine social symbionts of ants and termites displaying behavioral interactions with hosts (chemical manipulation of host ant by *Lomechusa* and grooming host ant by *Sceptobius*) and dramatic morphological divergence (myrmecoid body shape of the myrmecophile *Diploeciton* and physogastric body shape of the termitophile *Neodioxeuta*).

The unparalleled diversification of Aleocharinae has been attributed in part to the beetles’ possession of a defensive "tergal gland"—a structure positioned on the dorsal abdomen that is targetable at other organisms, and which in most species produces a potent, benzoquinone-containing secretion (*24, 31, 32*) (**Fig. 1B, C**). The tergal gland confers effective protection against predators such as ants (*32–37*), and is thought to have enabled aleocharines to radiate explosively in ant-dominated ecosystems worldwide (*38, 39*). The gland has also been proposed to facilitate infiltration of ant and termite colonies, leading to widespread convergent evolution of symbiotic myrmecophiles and termitophiles across the subfamily (*28, 38, 40–43*) (**Fig. 1D**). Tergal gland chemistry has been shown to vary among aleocharine taxa, reflecting potential adaptive streamlining of the secretion to specific niches (*31, 37, 44–46*). The secretion has also been found to possess antimicrobial properties, a function that may aid in colonizing new habitats via pathogen suppression (*32*). Crucially, early branching aleocharine lineages and related outgroup staphylinid subfamilies lack the tergal gland (*31, 47*), and are correspondingly species-poor with limited ecological diversity (*38*) (**Fig. 1A**). In contrast, the gland is conserved across the vast majority of the 10^4^–10^5^ "higher" aleocharine species, secondarily degenerating only in specialized symbiotic taxa where chemical defense is obsolete (*38, 43*). The gland is thus a putative key innovation (*2, 48*)—a trait that is correlated with, and likely contributed to, the exceptional cladogenesis of aleocharines and their diversification into countless ecological niches.

Insights into the function and evolution of the tergal gland have come from studies of the aleocharine *Dalotia coriaria*, revealing how this structure is composed of two secretory cell types that synergize to produce the defensive secretion (*32*). One cell type—the ’BQ cells’—converts dietary aromatic amino acids into toxic benzoquinones. These compounds are solids, however, and depend on the second cell type, the "solvent cells", to synthesize fatty acid-derivatives into which the benzoquinones can dissolve. The resultant cocktail is highly aversive to predators, conferring adaptive value onto this cooperative biosynthetic system (*32*). Here, we retrace the evolution of this chemical innovation in Aleocharinae. We present a chromosome-level reference genome of *Dalotia coriara*, along with draft genome assemblies spanning the Aleocharinae phylogeny. By combining comparative genomic and single-cell transcriptomic insights with analyses of enzyme function, gland chemistry and cellular anatomy, we pinpoint molecular and cellular contingencies that established the tergal gland during basal aleocharine cladogenesis. We show that, since its origin, the cell types comprising this structure have exhibited evolutionary stasis at both the functional and molecular levels as Aleocharinae radiated globally into tens of thousands of lineages. Despite this overt conservation, we present evidence that tergal gland cell types have provided versatile substrates for the emergence of biochemical novelties, providing a catalyst for profound niche specialization in this beetle clade. Our findings connect molecular evolutionary processes underlying the origin and evolution of novel cell types to the ecological and macroevolutionary diversification of a major metazoan radiation.

## Results

### The *Dalotia coriaria* reference genome

To enable broad insights in rove beetle biology, we assembled a high quality, chromosome-level genome of the laboratory model staphylinid, *Dalotia coriaria* (Aleocharinae: Athetini) (**Fig. 2A**). Our approach combined Illumina short paired-end reads (44× coverage) with Oxford Nanopore minION long-reads (54× coverage, N_50_ =7,933) to create an initial 120 Mb draft assembly, *Dcor* v1 (N_50_=3.97 Mb, longest scaffold=12.92 Mb; **Table S1**). The *Dcor* v1 assembly size was close to that predicted by *k*-mer based tools (139 ± 20 Mb), but less than half the flow cytometry estimate (male 294 ± 11 Mb and female 296 ± 13 Mb, **Fig. S1A, B**). Large discrepancies between *k-*mer-and flow-based genome size estimates have recently been observed in beetles (*50, 51*) and can arise from highly repetitive content (*51*). The estimated repeat content of the *Dalotia* genome based on short reads from two separate specimens was 65 to 69% (**Fig. S2A**), composed primarily of a specific 147 bp AT-rich satellite (*Dc-Sat1*) that comprises 55 to 61% of the repeatome (**Fig. S2B, C and Table S2**), found almost exclusively in intergenic regions (**Fig. S2D**). *Dc-Sat1* is not unique to *Dalotia* but has undergone a recent species-specific expansion to dominate the repeat landscape (**Fig. S2B**), consistent with the ’library’ model of satellite evolution (*52*). We found numerous long-reads entirely composed of *Dc-Sat1* arrays and predict these could form kilobase to megabase-scale, higher-ordered DNA structures (**Fig. S2G**) (*53*).

**Figure 2.**
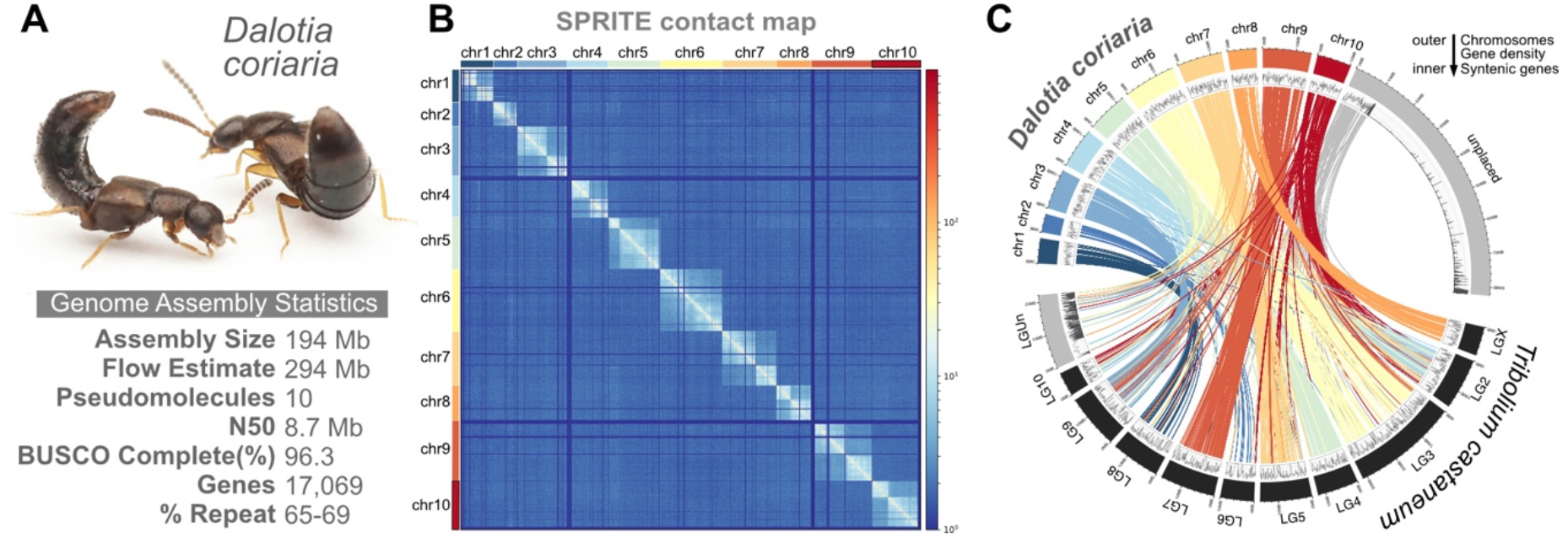
The *Dalotia* reference genome. **(A)** The genome assembly statistics of the free-living, higher aleocharine *Dalotia coriaria*, **(B)** Contact map assembled by Split-Pool Recognition of Interactions by Tag Extension (SPRITE) reveals ten pseudomolecules (chromosomes) of *D. coriaria*. **(C)**Gene density (density plot, middle band) and synteny (links, inner band) between *D. coriaria* and *Tribolium castaneum* chromosomes or linkage groups (outer band). The inner links are colored according to the originating *D. coriaria* chromosome.

To further extend and orient scaffolds, we generated 262 Bionano optical maps and performed a hybrid assembly with *Dcor* v1. *De novo* assembly of optical maps alone produced a 257 Mb assembly, approaching the flow estimate, but the hybrid assembly with *Dcor* v1 incorporated only 96 of those optical maps, giving an assembly size of 122.8 Mb (*Dcor* v2, **Fig. S3A** and **Table S1**). We were able to map 883 10kb or longer minION reads to 124 unincorporated optical maps (74%), suggesting shared repeat structures in the long-read data and optical maps that may not be captured in the hybrid assembly (**Fig. S3B**). We also uniquely mapped 95% of the short- and long-reads to the *Dcor* v2 assembly, indicating that abundant repeats like satellite *Dc-Sat1* are present but collapsed in the *Dcor* V2 assembly. We then produced a chromosome-resolved assembly via Split-Pool Recognition of Interactions by Tag Extension (SPRITE) (*36*)—a method that yields both intra- and interchromosomal contact information (See “*Dalotia* Genome Assembly” in **Methods**). After generating a DNA contact map with 11,674,733 clusters identified by SPRITE, we were able to further improve contiguity into 10 pseudomolecules that contain 98.9% of the *Dcor* v2 assembly with a scaffold N_50_ of 12 Mb (*Dcor* v3, **Fig. 2B** and **Table S1**). The number of pseudomolecules (referred to as chromosomes hereafter) matches the chromosome counts (**Fig. S1C**) and previously published karyotypes of another aleocharine genus, *Aleochara* (*37*). Lastly, we recovered 72 Mb of unincorporated, repeat-rich contigs by mapping the preliminary assemblies back onto the *Dcor* v3 assembly. These repeat-rich contigs were combined with *Dcor* v3 for a final assembly size of 194 Mb (**Table S1**).

Gene content in the *Dcor* v3 assembly is near-complete with 96.3% complete/1.3% partial orthologues recovered from the BUSCO arthropod gene set (n=1,013 genes) (*38*) (**Fig. S4**). We predicted 17,069 protein coding genes using a combination of transcriptome data spanning life stages and tissue types, predicted gene models from the red flour beetle *Tribolium castaneum* (Tenebrionidae) and burying beetle *Nicrophorus vespilloides* (Staphylinidae: Silphinae), as well as *ab initio* tools (see **Methods**) (**Table S3**). Despite a >250 million year phylogenetic distance, gene synteny remains high between *Dalotia* and *Tribolium* (**Fig. 2C**), with 878 syntenic blocks recovered that contain 3–10 shared genes per block. We determined Chr 8 to be the probable X chromosome based on 12.9%, 18.7% and 8.6% protein conservation with *Tribolium* and two rove beetles of the subfamily Staphylininae—*Ocypus olens* and *Philonthus cognatus*, respectively (**Fig. 2C and Fig. S5A**). Chr 2 was identified as the Y chromosome based on significant male-biased expression (χ^2^(54, *N* =13244) = 522.7, *p* < 0.001), but shared little gene content with the putative Y of *P. cognatus* (0.2%) (**Fig. S5**).

### Phylogenomic relationships in Aleocharinae

To explore patterns of genome evolution in Aleocharinae, we generated short-read genomic data for a further 24 ingroup and outgroup species, using *Dcor* v1 assembly to guide genome assembly and inform gene predictions across these taxa. Our taxon sampling was targeted to illuminate traits that arose during the basal cladogenesis of Aleocharinae, principally the tergal gland. We assembled genomes of three representatives of the glandless tribe Gymnusini—the earliest diverging clade within Aleocharinae (*26, 31, 47, 49, 54, 55*). Multiple genomes spanning major gland-bearing higher aleocharine lineages were incorporated, including members of putative early branching tribes within this huge clade: Hypocyphtini, Aleocharini and Oxypodini (*42, 47, 54, 55*).

Taxa from the tribes Mylaenini, Falagriini, Homalotini, Geostibini, Lomechusini and Athetini (to which *Dalotia* belongs) were also included. Among these, genomes of four myrmecophiles were assembled to illuminate evolutionary changes in chemistry associated with symbiosis. Three are members of the "*Ecitochara* group" of Athetini (formerly the tribe Ecitocharini)—a Neotropical clade of ant-mimicking (myrmecoid) symbionts, obligately associated with *Eciton* army ants, in which the tergal gland has undergone secondary degeneration (*56*). The fourth myrmecophile is *Liometoxenus newtonarum* (Oxypodini), a symbiont of *Liometopum* ants from Southern California (*57*). Outgroup genomes were included from members of the subfamily Tachyporinae, allied to Aleocharinae within the Tachyporine-group of Staphylinidae (*34*). On average, the genome completeness for the 24 new assemblies was 92.6% (range 54.7% to 99.5%) (**Fig. S4, Table S3**). Previously published genomes of nine other beetles of high genome completeness were also included, spanning the coleopteran suborder Polyphaga (to which Staphylinidae belongs).

We used a set of 1520 orthologous protein-coding loci to infer a phylogenomic tree of these species, estimating the ages of key nodes using a set of fossil calibrations both within and outside of Aleocharinae (**Fig. 3A, Table S4**). Our topology is strongly supported at all nodes (**Fig. S6A, B**), and broadly congruent with prior molecular phylogenetic studies (*42, 54, 55*). We recovered a monophyletic Aleocharinae, sister to the three tachyporine taxa, and with a crown-group origin in the Early Jurassic (178 Ma; 95% Highest Posterior Density (HPD): 209–150 Ma). Within the Aleocharinae, glandless Gymnusini are resolved as sister to a monophyletic higher Aleocharinae (clade "HA") (*26, 47, 55, 58*). We infer that the tergal gland originated close to the Jurassic-Cretaceous boundary, with the HA crown-group dating to 148 Ma (95% HPD: 176–123 Ma). Consistent with previous studies, Hypocyphtini emerge as the earliest-branching HA lineage (*47, 54, 55, 59*), with Aleocharini the subsequent HA lineage to diverge. Inside the HA, the homalotine *Leptusa* was recovered as sister to the two oxypodine taxa (*Oxypoda* and the myrmecophile *Liometoxenus*), while taxa belonging to the megadiverse "Athetini-Pygostenini-Lomechusini" ("APL") clade (*60, 61*) are recovered as monophyletic, including the tribe Geostibini. We estimate that the APL clade originated in the early Paleocene (64 Ma; 95% HPD: 77–53 Ma)— a younger age than previously estimated (*42*). The APL clade numbers ∼8600 extant described species and includes the greatest number of myrmecophile and termitophile lineages. Its Cenozoic origin implies an exceptional rate of cladogenesis, with recurrent transitions to social insect symbiosis during a window in which modern ants and termites are both thought to have proliferated (*39, 62–64*). Within the APL clade, the myrmecophilous *Ecitochara* group emerges as sister to the remaining athetines, *Dalotia* and *Atheta*, a finding congruent with earlier studies of relationships in Athetini (*60, 61*) (**Fig. 3A**).

**Figure 3.**
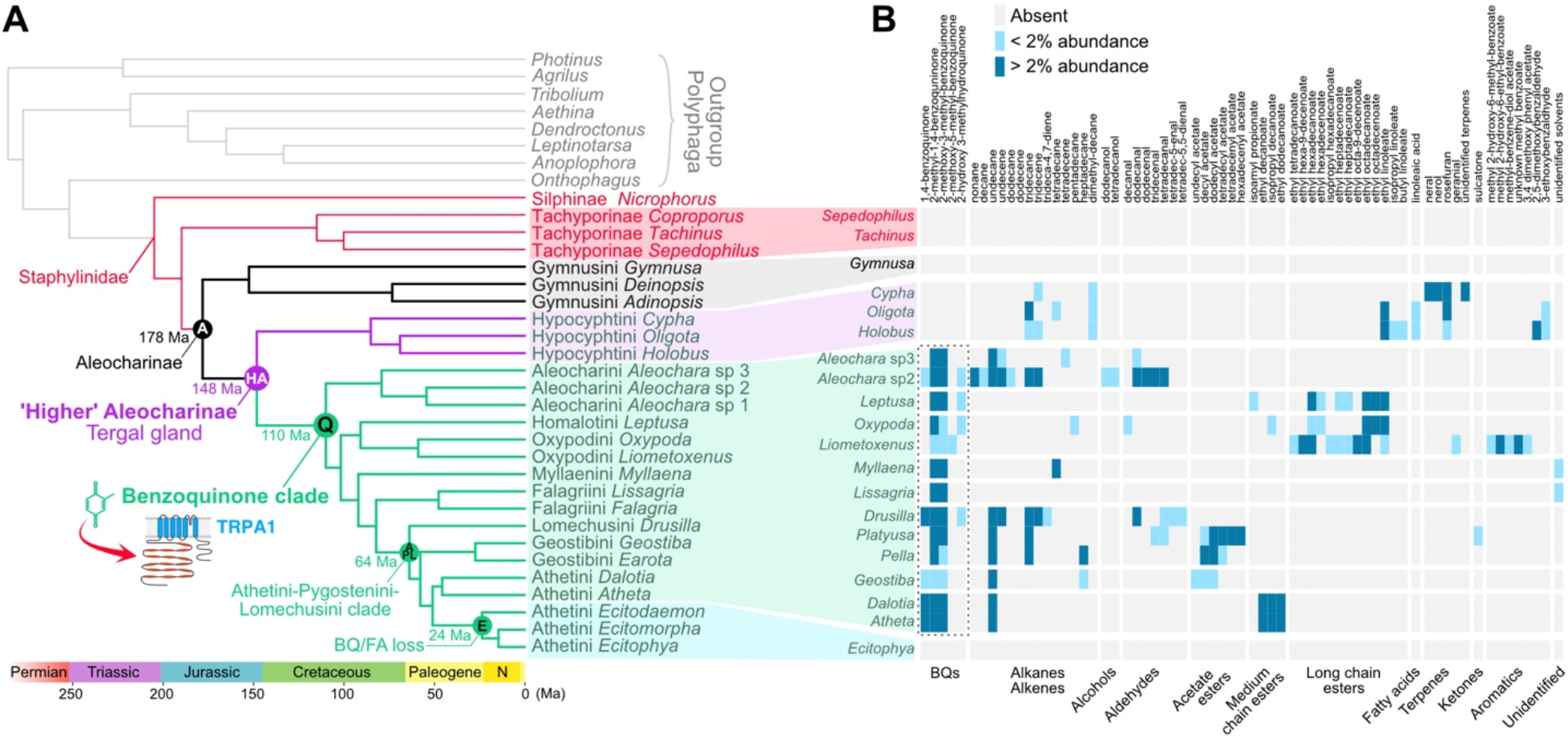
Chemical innovation of rove beetles over 100 million years. **(A)** Dated phylogenomic tree based on 1520 orthologs, with key nodes and their respective ages indicated. **(B)** Heat map of major and minor compounds found in aleocharine tergal glands. The dashed box indicates the broad conservation of a small number of benzoquinones across the majority of aleocharines—the exceptions being the earliest-branching Gymnusini, Hypocyphtini and the *Ecitochara*-group of Athetini.

### Chemical evolution in Aleocharinae

We used the tree topology to explore patterns of chemical evolution across Aleocharinae. We extracted tergal gland secretions from a diversity of taxa spanning the tree and used gas chromatography-mass spectrometry (GC-MS) to characterize the chemical composition (**Fig. 3B**). The "classical" aleocharine tergal gland secretion employs benzoquinones (BQs) as toxic irritants. BQs bind TRPA1 channels (*65*), activating nociceptive neurons to induce a pain response. BQs are solid compounds, however, and are therefore dissolved in a fatty acid (FA)-derived fraction composed of either alkanes, alkenes, aliphatic esters, aldehydes, or a combination thereof. The FA-derived solvent unlocks the BQs’ potency, creating a highly noxious secretion that confers adaptive value on the beetle’s chemical defense (*32*). Consistent with previous studies (*31, 32*), we find that a BQ/FA cocktail is common to the majority of higher aleocharine taxa. The lineages that produce such a secretion comprise a vast clade within the HA that we herein refer to as the "Q clade" (quinone-producing). We infer that the most recent common ancestor (MRCA) of the Q clade arose ∼110 Ma (95% HPD: 132–90 Ma; **Fig. 3A**). *Aleochara* (Aleocharini) represents the earliest-branching lineage within the Q clade (**Fig. 3A, B**). Subsequent to the origin of the BQ/FA cocktail, relative chemical stasis is observed almost throughout subsequent cladogenesis, particularly within the benzoquinone fraction, where a small number of variants of 1,4-benzoquinone are conserved across the majority of Q clade taxa (some species also secrete trace levels of the BQs’ hydroquinone precursors) (**Fig. 3B**).

In contrast, while the use of FA derivatives as BQ solvents is similarly conserved, the precise compounds vary substantially, exhibiting a mosaic of gains and losses. We and others have previously shown how subtle changes in the chain lengths and chemical ratios of the FA-derived fraction strongly influence the secretion’s viscosity, surface coating ability, and efficacy as a solvent for the BQs (*32, 66*). Marked variation in FA derivatives across the Q clade implies that different lineages have modified the physicochemical properties of their secretions. For example, production of medium-chain, acetate- and some long-chain esters appears to have evolved independently within the Q clade (**Fig. 3A, B, Fig. S7**). Esters have been shown to increase the wetting properties of defensive secretions (*67*), and the low-level production of esters in some taxa is consistent with these compounds being surfactants rather than the principal solvent (*32, 67*). In *Dalotia*, inclusion of low-level, medium-chain esters has also been shown to be critical for *Dalotia*’s defensive secretion to suppress microbial growth (*32*). The addition of esters may therefore represent an adaptive enhancement of the secretion in these taxa. Moreover, esters have largely superseded alkanes as the primary solvent in the oxypodines *Oxypoda* and *Liometoxenus* and the homalotine *Leptusa*—potentially via a single secondary loss (or strong diminishment) of alkane production in the common ancestor of this clade (**Fig. 3A, B**). We note that, curiously, both alkanes and esters have been lost in the falagriine *Lissagria*—a finding consistent with earlier chemical data from Falagriini (*31*). Presently unidentified compounds are the likely solvents for the BQs in this tribe.

Evolvability of the tergal gland’s secretion is further underscored by the finding that, despite the general conservation of a BQ/FA cocktail across much of the Aleocharinae, many groups scattered across the tree have incorporated novel compound classes into the secretion, including ketones, terpenes and other aromatics (**Fig. 3B**). Tergal gland chemistry therefore appears to be reprogrammable during evolution, leading to taxon-specific secretions that may facilitate adaption to certain niches. The function of the tergal gland can apparently also become dispensable: species within the *Ecitochara*-group have secondarily lost BQs as well as any solvent compounds (**Fig. 3B**), consistent with degeneration of the gland in these socially integrated myrmecophiles (*56*).

### Stasis in gland cell type evolution

We asked how changes at the genomic, pathway, and cell type levels underlie the evolution of tergal gland chemistry. The gland is composed of two secretory cell types that are unique to aleocharines, and which work together to produce the secretion. The "BQ cells" manufacture the benzoquinones, while the "solvent cells" produce the fatty acid derivatives into which the BQs dissolve (**Fig. 1B, C**). Previously, we generated BQ and solvent cell type-specific transcriptomes from *Dalotia coriaria*, enabling us to elucidate biosynthetic pathways that produce *Dalotia*’s BQ/FA cocktail (*32*). To gain insight into the origins and functional evolution of the BQ and solvent cells, we sought to retrace their evolution prior to and during the HA clade’s diversification. *Dalotia*’s tergal gland secretion contains three benzoquinones; these are dissolved in a large volume of a medium chain alkane solvent, undecane, along with three aliphatic esters: ethyl decanoate, isopropyl decanoate and ethyl dodecanoate (**Fig. 4A, upper trace**). The earliest-branching lineage that produces a BQ/FA cocktail comparable to that of *Dalotia* is *Aleochara* (the tribe Aleocharinae), the two lineages demarcating the Q clade that encompasses the entire higher Aleocharinae minus the tribe Hypocyphtini (**Fig. 3A**). Although *Aleochara* diverged from *Dalotia* in the Early Cretaceous, 110 Ma (**Fig. 3A, Fig S6-7**), species of *Aleochara* nevertheless produce two or all three of the same BQs as *Dalotia* (**Fig. 3B**). Similarly, these BQs are dissolved in alkanes, predominantly undecane and tridecane; in some *Aleochara* species the solvent additionally contains corresponding aldehyde precursors, as well as alkenes. Unlike *Dalotia*, however, *Aleochara* secretions do not apparently contain esters (**Fig. 3B**) (*31, 68*).

**Figure 4.**
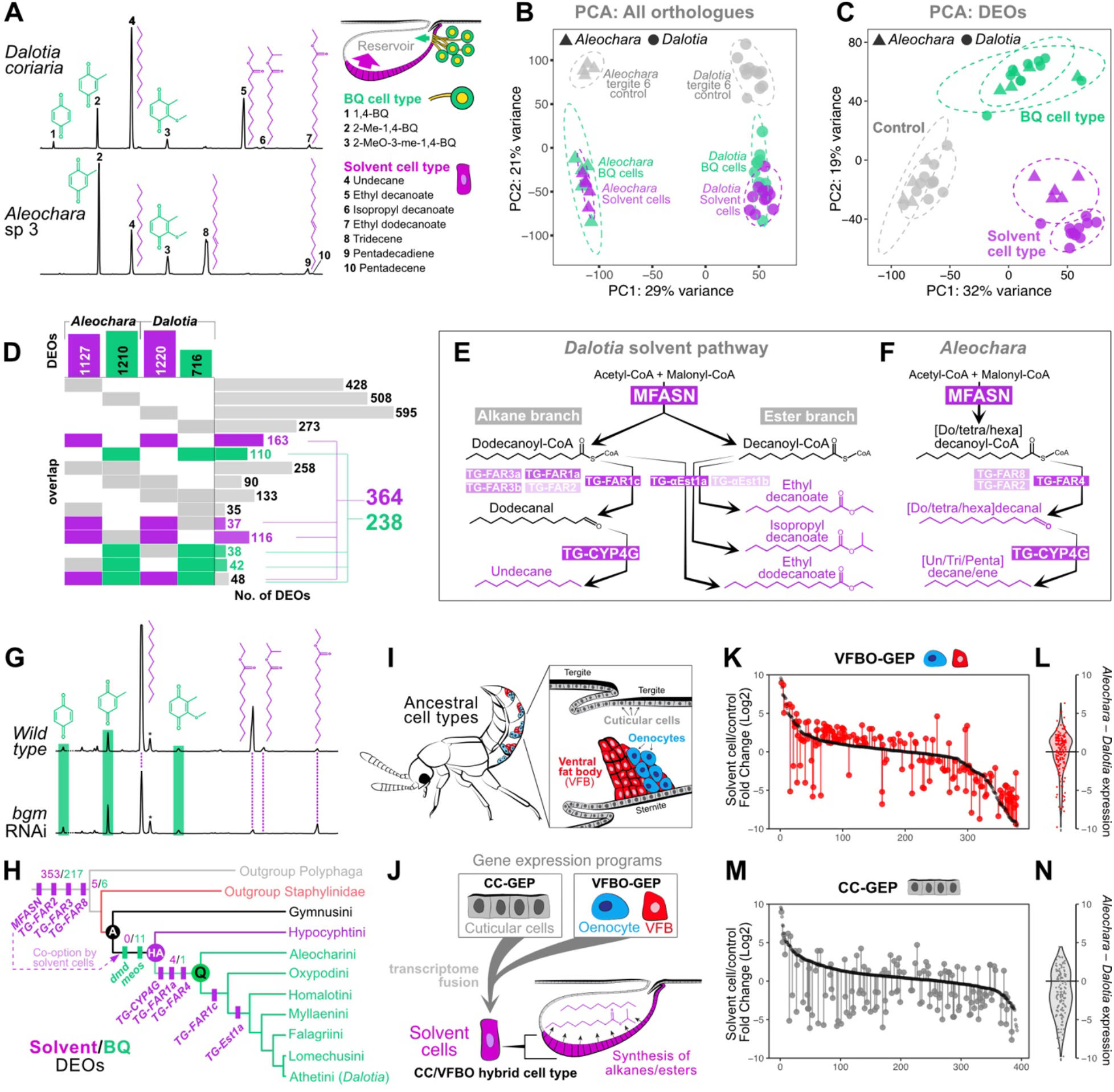
Deep conservation of tergal gland gene repertoire in the Q clade. **(A)** GC traces of *Dalotia* and *Aleochara* glandular compounds and their cell type of origin. **(B)** PCA of all expressed orthologues in *Dalotia* and *Aleochara* solvent cells, BQ cells and control tissue (tergite 6). **(C)** PCA of 2837 DEOs in *Dalotia* and *Aleochara* cell types and control tissue. **(D)** UpSet plot showing shared DEOs for each cell type by species and cell type. **(E-F)** Solvent pathways in *Dalotia* and *Aleochara*, with cases of paralogue co-expression in solvent cells. Transparency of the purple boxes is equal to the maximum log_2_ fold-change above the control tissue for paralogues. **(G)** Example GC traces of glandular compounds in wildtype animals (top trace, n=14) and *bgm*-silenced animals (n=42). **(H)** Cladogram of Aleocharinae and outgroups showing origins of enzyme paralogues involved in solvent and benzoquinone biosynthesis. **(I-J)** Schematic of abdominal cell types with distinct gene expression programs (GEPs) for the ventral fat body/oenocytes and cuticle cells (**I**), hybridization of which created the solvent cell type (**J**). **(K, M)** *Aleochara* solvent cell expression (red or grey) relative to *Dalotia* solvent cell expression (black) for orthologues of the highest z-score ranked genes that belong to VFBO-GEP and CC-GEP. **(L, N)** Violin plots showing the difference in *Aleochara* from *Dalotia* solvent cell expression for all genes within each GEP.

We assembled a draft genome of a Southern Californian species of *Aleochara* (*Aleochara* sp. 3 in **Fig. 3**), secretions from which share with *Dalotia* two BQs (2-methyl-1,4-BQ and 2-methoxy-3-methyl-1,4,-BQ) and undecane (**Fig. 4A, lower trace**). We dissected replicates of BQ and solvent cells from tergal glands of this *Aleochara* species and assembled single-cell transcriptomes via SMARTseq, creating a data set directly comparable to that obtained from the homologous cell types in *Dalotia* (**Fig S8, File S1**). To explore gene expression evolution between *Dalotia* and *Aleochara* BQ, solvent and other abdominal cell types, we performed principal component analyses (PCA) on replicate SMARTseq transcriptomes, restricting our analysis to the 9314 orthologous loci that are shared between these two beetle species. We performed two PCAs: 1) all orthologous transcripts, including constitutive "housekeeping" loci that are not differentially expressed in any cell type; 2) "Differentially Expressed Orthologues" (DEOs)—the subset of orthologues that are significantly differentially expressed in at least one pairwise comparison between cell types, and which account for the transcriptomic (and hence likely functional) differences between cell types.

Notably, PCA based on all orthologous transcripts strongly separated the cell types by beetle species along PC1, indicating a pervasive, species-specific evolutionary divergence in gene expression that is shared by the BQ and solvent cell transcriptomes within a species (**Fig. 4B**). Conversely, however, when the analysis was restricted to the 2837 DEOs, each cell type now clustered with the homologous cell type from the other beetle species, with strong separation of BQ and solvent cells both from each other and from control tissue (**Fig. 4C**). Hence, despite the ∼110 MY separation between *Aleochara* and *Dalotia*, their BQ and solvent cells each differentially express common gene sets that likely underlie the conserved biosynthetic functions of these cell types across the Q clade. Relative evolutionary stasis of the DEOs in each cell type has occurred despite marked, species-specific divergence in the remaining constitutive fraction of the transcriptome.

### A core, conserved solvent cell transcriptome

We examined the basis for the transcriptomic similarity between *Dalotia* and *Aleochara* tergal gland cell types and identified 364 DEOs in the solvent cells of both species and 238 DEOs shared by their BQ cells (**Fig. 4D**). These sets of DEOs define deeply conserved "core" transcriptomes within each cell type. We asked if these core transcriptomes might encompass ancient biosynthetic toolkits within the Q clade, and found that tergal gland cells of *Dalotia* and *Aleochara* express homologous pathways for defensive compound biosynthesis. In *Dalotia*, synthesis of the alkane and esters by solvent cells derives from a bifurcating fatty acid pathway in which a fatty acid synthase, Master FASN (MFASN), produces C10 and C12 fatty acid precursors (**Fig. 4E**). In one downstream pathway branch, the C12 fatty acid is reduced to an aldehyde by a fatty acyl-CoA reductase (Tergal Gland FAR1c; TG-FAR1c, formerly "TG-FAR" (*32*)); the aldehyde is then decarbonylated by a 4G-class cytochrome P450 (TG-CYP4G), yielding undecane. In a parallel branch, the C10 fatty acid is esterified by a carboxylesterase of the ɑ-esterase family (TG-ɑEst1a; formerly *"*TG-ɑEst*"* (*32*)), resulting in the two C10 esters (**Fig. 4E**). Traces of the C12 fatty acid are also esterified by TG-ɑEst1a to make the ethyl dodecanoate (**Fig. 4E**).

Core components of this pathway appear deeply conserved within the Q clade. As in *Dalotia*, MFASN is the sole fatty acid synthase expressed in *Aleochara*’s solvent cells (**Fig. 4F, File S1, Fig. S9A**); likewise, the terminal alkane decarbonylase, *TG-CYP4G*, exists as a single copy gene in both *Dalotia* and *Aleochara* and comprises part of the solvent cell core transcriptome (**Fig. 4C, File S1, Fig. S9B**). Multiple other core components of the solvent cell transcriptome have predicted roles in solvent biosynthesis, and the core transcriptome is significantly enriched in biological processes related to fatty acid synthesis and modification (**Table S5**). One previously uncharacterized step in solvent production is the essential activation of fatty acids produced by MFASN by the addition of CoA (*69*). Among core solvent cell transcripts, we identified a very long-chain-fatty-acid-CoA synthase (LC-FACS), orthologous to the *Drosophila* gene *bubblegum* (*bgm*) (**Fig. S9C**). Silencing *bgm* expression in *Dalotia* with RNAi caused a significant reduction in undecane levels (avg. 41% of GFP control, Wilcoxon signed-rank with Bonferroni p-adjusted = 0.005), and a near-complete loss of ethyl decanoate (avg. 12% of GFP control, p-adjusted < 0.001; **Fig. 4G, Fig. S9D**). We suggest Bgm is at least partially responsible for activation of fatty acid precursors of defensive alkanes and esters in Q clade aleocharines.

Beyond a core transcriptome of orthologous loci, the use of functionally equivalent paralogues can also be seen in *Aleochara* and *Dalotia*. *Dalotia* solvent cells express five *FAR* paralogues (**Fig. 4E**), one of which, *TG-FAR1c*, accounts for virtually all undecane synthesis (*32*). In every *Aleochara* genome we surveyed, however, an orthologue of *TG-FAR1c* was not detected (**Fig. S10**). Instead, *Aleochara* solvent cells express three *FAR* paralogues—*TG-FAR2*, *4* and *8*, one or more of which likely perform the equivalent step in alkane synthesis (**Fig. 4F, Fig. S10**). The *FAR* family undergoes extensive gene birth-and-death in insects (*70*). Indeed, weak expression of *TG-FAR2* in *Dalotia* solvent cells may be a vestige of its earlier involvement in alkane production prior to the more recent birth of *TG-FAR1c* (**Fig. 4M; Fig. S10**). In total, 27 *FAR* copies are encoded in the *Dalotia* genome and 21 in *Aleochara* (**Fig. S10**). Evolutionary turnover in the expression of duplicates from such large enzyme families has likely contributed to the conserved function of the solvent cell type in Aleocharinae, but has also been an apparent source of novelty. One key difference between the two beetles’ solvent pathways is the ester branch, which is present in *Dalotia* but not in *Aleochara* (**Fig. 4E, F**). *Dalotia*’s production of esters is mediated by *TG-ɑEst1a*—a gene with no apparent orthologue in *Aleochara* (**Fig. S11**). Indeed, no functionally equivalent ɑ-esterases or members of other carboxylesterase families are expressed whatsoever in *Aleochara*’s solvent cells (**Fig. S11, File S1**). Appending an ester branch to the pathway was therefore a more recent innovation in solvent cell evolution, contingent on the birth and recruitment of *TG-ɑEst1a*.

### Solvent cell evolution through ancient transcriptome hybridization

Notably, the solvent cell core transcriptome is composed of a majority of co-opted ancient genes, with a minority of recent, taxon-restricted duplicates that arose in the HA or Q clades. We infer that 353 of 364 core loci have orthologues present across the outgroup Polyphaga (**Fig. 4H**). Such strong predominance of co-option may stem from how the solvent cell type is thought to have originated. Solvent cells are a secretory cell type but nevertheless form part of the beetle’s exoskeleton: they are continuous with the chitinous epidermis, secrete chitin themselves, and comprise a region of intersegmental membrane between tergites 6 and 7 (*31, 33, 71, 72*). Solvent cells have therefore been postulated as a ’hybrid’ cell type, with properties of both exoskeleton-forming and glandular cells. Using a single-cell RNA-seq atlas of *Dalotia*’s abdominal cell types, we previously showed that solvent cells are indeed a transcriptomic hybrid composed of two major gene expression programs—one that defines abdominal cuticular cells (the ’cuticular cell’ gene expression program; CC-GEP), and another that defines two ancient metabolic cell types: adipocyte-like ventral fat body cells, and oenocytes that produce cuticular hydrocarbon pheromones (Ventral Fat Body/Oenocyte-GEP; VFBO-GEP) (*32*) (**Fig. 4I**). The VFBO-GEP is strongly enriched for gene products involved in fatty acid synthesis and modification (*32*), implying that co-option of VFBO-GEP into cuticular cells provided the biosynthetic machinery for solvent production, transforming ancestral cuticle into solvent cells (**Fig. 4J**).

Given the deep conservation of solvent cells, this hybridization event was likely ancient within Aleocharinae. We examined the extent to which VFBO-GEP and CC-GEP are conserved in *Aleochara* and *Dalotia* solvent cells. We first ranked *Dalotia* loci according to their z-score within both VFBO-GEP and CC-GEP, and then compared expression of each *Dalotia* locus in the solvent cells to that of its orthologue in solvent cells of *Aleochara*. Strikingly, orthologues of *Dalotia*’s VFBO-GEP loci are also differentially expressed in solvent cells of *Aleochara*, the relative expression of these orthologues being strongly correlated between the two species (Spearman r=0.72, p <0.001; **Fig. 4K, L**). Conversely, conservation of CC-GEP expression in solvent cells is weaker, with fewer *Aleochara* orthologues showing comparable expression to *Dalotia* solvent cells (Spearman r=0.21, p=0.013; **Fig. 4M, N**). These findings imply that formation of the solvent cells, via recruitment of VFBO-GEP into cuticle cells, pre-dates the Q clade MRCA. Subsequent conservation of VFBO-GEP in solvent cells has occurred despite greater divergence in the transcriptional program for this hybrid cell type’s cuticular identity.

### Evolution of benzoquinone production and the BQ cell type

Akin to the solvent cells, we find evidence of deep evolutionary stasis at the molecular level within the BQ cell type. In *Dalotia*, benzoquinones have been shown to derive from dietary aromatic amino acids such as tyrosine (*32*) (**Fig. 5A, B**). These are converted to 4-hydroxybenzoic acid (4-HB), which is modified in the mitochondrion of BQ cells via sequential actions of ubiquinone/coenzyme Q pathway enzymes. The resultant hydroquinones are then thought to be secreted into the lumen of the BQ cell, where they undergo oxidation by a secreted multicopper oxidase of the laccase family, named Decommissioned (Dmd), which converts the hydroquinones into the final, toxic BQs (**Fig. 5A, B**). Critical components of this pathway are conserved in *Aleochara*. As in *Dalotia*, the *Aleochara* genome contains a single orthologue of *dmd*, which is amongst the most strongly upregulated transcripts in *Aleochara* BQ cells (**Fig. 5B, Fig. S8**).

**Figure 5.**
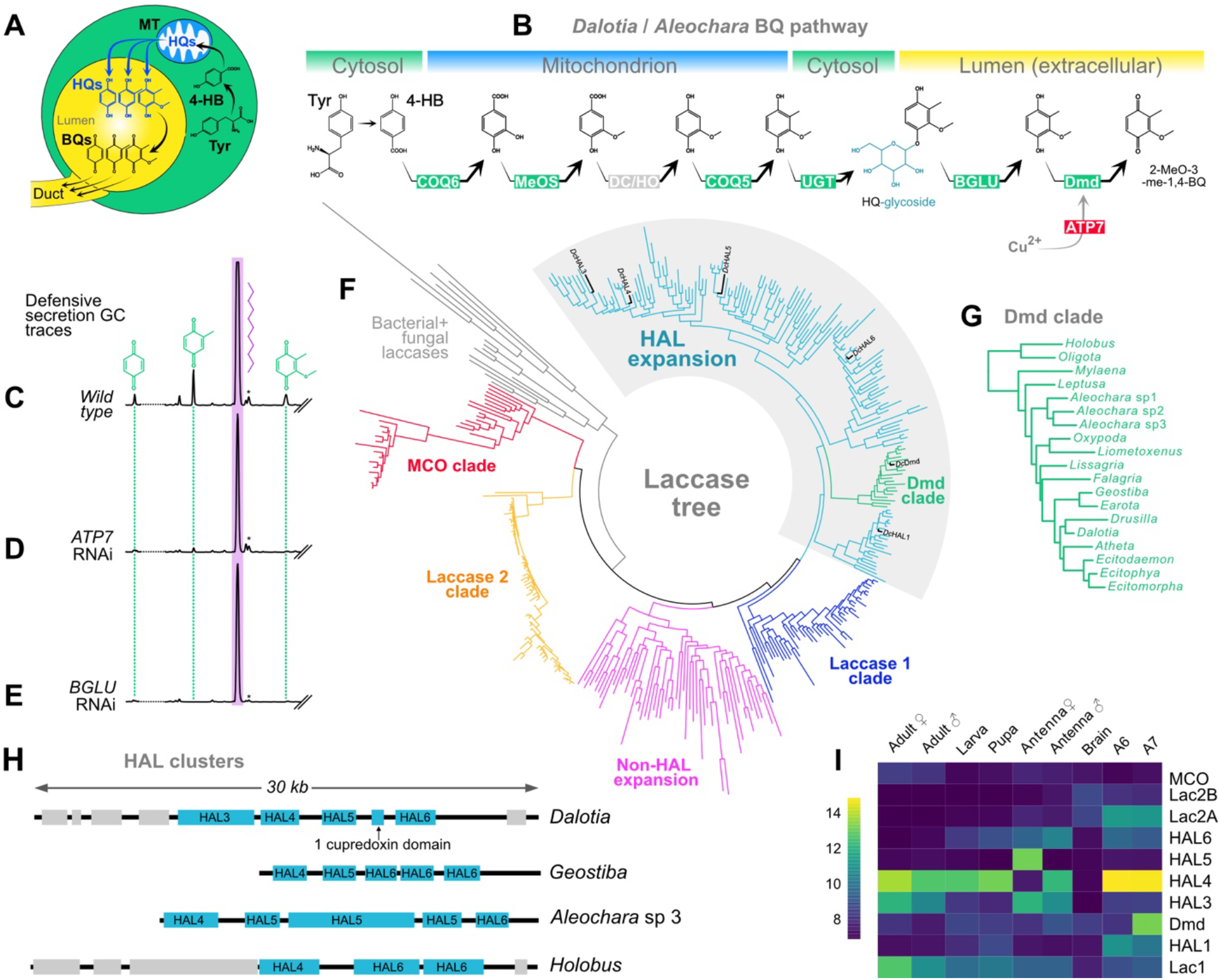
Evolution of BQ chemistry. **(A)** Cartoon of BQ cell showing route of benzoquinone synthesis from dietary tyrosine (Tyr) through to secreted benzoquinones. (**B**) The benzoquinone pathway in Q clade Aleocharinae, showing cellular locations of each enzymatic step. **(C–E)** GC traces of glandular compounds in wildtype *Dalotia* (**C**) and *ATP7*-silenced animals (**D**) and *β-glucosidase -*silenced animals. Levels of benzoquinones are strongly and selectively diminished by *ATP7* or *β-glucosidase* silencing, while levels of undecane are unchanged. Dotted line indicates position of hexane contamination peaks that have been removed for clarity. Asterisks denote peaks of dimethyl-BQ spiked in as a positive control. **(F)** ML tree of laccase gene family showing different clades and expansion of laccases in higher aleocharines (Higher Aleocharine Laccase (HAL) expansion in light blue, *Dalotia* HAL paralogues are indicated). Substitution model LG+R10 with 1000 bootstrap replicates. **(G)** Expanded tree of Dmd orthologues from panel **F** reveals conservation of this laccase across higher aleocharine taxa. **(H)** The genomic HAL cluster of selected aleocharine taxa. **(I)** Expression heat map of *Dalotia* laccases, including HAL paralogues, from RNAseq data obtained from tissues, life stages and sexes.

Hydroquinone oxidation by Dmd is thus likely an ancient, terminal step in BQ biosynthesis (**Fig. 5B**). We also find evidence that upstream mitochondrial steps are conserved. For example, like most BQ-clade aleocharines, both *Dalotia* and *Aleochara* produce 2-methoxy-3-methyl-1,4-BQ (**Fig. 4A**). In *Dalotia*, the methoxy group is added to the aromatic core by a mitochondrial enzyme, Methoxyless (MeOS)—an aleocharine-specific duplicate of the enzyme COQ3 that adds a methoxy group to ubiquinone (*32*). We recovered a single *meos* orthologue in *Aleochara*, which is again a component of the BQ cell type’s core transcriptome and likely functions identically (**Fig. 5B, Fig. S8, Fig. S12A**).

As a whole, the BQ core transcriptome is enriched in biological processes related to mitochondrial metabolism and metal ion transport (**Table S5**). Other core transcripts represent newly discovered components with putative functions in benzoquinone synthesis, secretion or trafficking. The activity of laccases related to Dmd has been shown to depend on elevated cellular import of Cu^2+^—a process that in mammalian systems is mediated by ATPase transporters ATP7A and B and the cytosolic copper chaperone ATX1 (*73, 74*). Conspicuously, both the single copy aleocharine homolog of mammalian *ATP7A and B* and the *ATX1* orthologue comprise part of the BQ cell type’s core transcriptome (**Fig. S8, Fig. S12B)**. We silenced the ATP7 transporter in *Dalotia* and observed strongly diminished levels of the highest abundance BQ in the secretion, 2-methyl-1,4-BQ (Wilcoxon signed-rank with Bonferroni p-adjusted = 0.0267, **Fig. 5C, D, Fig. S12C**). Elevated Cu^2+^ in BQ cells is thus likely essential for Dmd activity, providing the cofactor for this metalloenzyme (**Fig. 5B**).

Upstream of Dmd, the mechanisms of intracellular trafficking of hydroquinone precursors were previously unknown. Despite the widespread use of benzoquinones in arthropod chemical defenses (*75*), it has been unclear how these highly cytotoxic compounds are safely produced and transported inside cells prior to secretion (*32, 76*). In plants, small molecule toxins are often conjugated to sugars, creating glycosides that render many such compounds relatively harmless (*77, 78*). Binding to a hydrophilic sugar group also facilitates the cytosolic transport of non-polar small molecules, and their secretion from the cell (*77, 78*). Upon herbivory, the glycoside is released from damaged cells, where it commonly undergoes cleavage by a β-glucosidase enzyme that removes the sugar moiety, thereby activating the toxin (*79*). An analogous mechanism of sugar conjugation and cleavage has previously been hypothesized as a possible means for benzoquinone regulation in insect defense glands (*80*). Remarkably, we noticed that one of the core BQ cell loci is a predicted β-glucosidase (BGLU). Moreover, silencing this BQ cell-expressed *BGLU* in *Dalotia* led to near-complete elimination of all benzoquinones from the tergal gland secretion (pBQ, meBQ and dimethylBQ Wilcoxon signed-rank with Bonferroni p-adjusted < 0.001 for each compound; **Fig. 5C, E, Fig. S12D**).

This result strongly implies that hydroquinone glycosides may indeed be the form in which the BQ cells transport hydroquinones through the cytosol; further, that a specific β-glucosidase enzyme is responsible for cleavage of the sugar moiety, releasing the toxin. The BQ cell BGLU encodes a secreted protein, implying that hydroquinone glycosides likely represent the form that is secreted into the BQ cell lumen, prior to their cleavage by BGLU and subsequent oxidation to benzoquinones by Dmd (**Fig. 5B**). Additional support for this model is the observation that among the most strongly upregulated core transcripts in both *Dalotia* and *Aleochara* BQ cells is a UDP-glycosyltransferase (UGT), an enzyme with a classical role in glycoside synthesis, in this case via conjugation of toxins to glucose or related sugars (*81, 82*). UGT is thus a candidate enzyme for producing the hydroquinone glycosides (**Fig. 5B**). Together, these results point to aleocharines employing a mechanism of benzoquinone regulation that has convergently evolved with small molecule chemical defense mechanisms found in plants.

As in solvent cells, the core transcriptome of BQ cells is composed predominantly of ancient, co-opted genes, with orthologues of 217 out of the 238 core loci occurring across polyphagan Coleoptera (**Fig. 4H**). Twelve loci, however, appear to represent aleocharine-specific novelties that arose in stem lineages of the HA or Q clades. We posit that the origins of some of these may have potentiated the evolution of benzoquinone synthesis in Aleocharinae. One of these loci is the *COQ3* paralogue *methoxyless*, which originated along the HA stem (**Fig. 4H**). *COQ3* is a single copy gene in most eukaryotes (*83*). However, the locus has repeatedly duplicated in both aleocharine and tachyporine rove beetles (**Fig. S12A**), yielding four copies in the *Dalotia* genome of which *meos* is specialized for benzoquinone methoxylation (**Fig. 5B**) (*32*). Most notably, *dmd* itself is specific to the HA, orthologues of this laccase occurring in genomes of all HA taxa assembled in this study (**Fig. 5G**). We retraced the origin of this novel laccase and found that *dmd* is but one of a major, monophyletic expansion of laccase enzymes that has emerged in the genomes of higher aleocharines. This "Higher Aleocharine Laccase" (HAL) clade encompasses six paralogues in *Dalotia* but up to fifteen in other species (**Fig. 5F, Fig. S13**).

We found evidence of significant episodic selection on almost all internal branches leading to the major splits in the HAL expansion, suggesting neofunctionalization of these duplicates (aBSREL select branch test, p<0.05, **Fig. S13**). The HAL copies can be dispersed within the genome, but many sit in tandem within a single genomic cluster (**Fig. 5H**). Irrespective of their genomic location, in *Dalotia* each HAL copy is expressed in a different pattern across tissues, developmental stages and sexes, implying that these novel duplicates have evolved distinct functions (**Fig. 5I**). We note that an independent "non-HAL" expansion exists within genomes of the earliest-diverging, glandless aleocharine tribe Gymnusini, as well in the genomes of closely related outgroup tachyporine rove beetles (**Fig. 5F**). The non-HAL clade must thus predate Aleocharinae and have been lost in higher aleocharines, where it was replaced with the HAL expansion. Aside from Dmd, the functions of these laccases are unknown. The genomes of most insects, including beetles, encode only three conserved laccases (**Fig. 5F**), including Laccase 2 that has a canonical function in pigmenting and sclerotizing the insect cuticle (*84, 85*). However, laccases in general are well known for efficiently oxidizing phenolic compounds (*86*), and we speculate that diversification of these enzymes in higher aleocharines may have enabled these beetles to better metabolize or detoxify soil-, plant-or fungal-derived phenolics (to which these beetles must be routinely exposed). We suggest that a byproduct of expanding the laccase repertoire in these beetles was the birth of a duplicate that was ultimately neofunctionalized for benzoquinone synthesis.

### Gland conservation and divergence in the earliest-branching HA lineage

Our findings uncover an ancient gland toolkit in the Q clade that has been preserved as these beetles radiated over the past ∼110 million years. Yet, the tergal gland is thought to predate the Q clade: this structure is a synapomorphy for the Higher Aleocharinae—a larger group that includes both the Q clade and a further, early-branching lineage: the small tribe Hypocyphtini (**Fig. 3A**) (*47, 54, 55, 59*). Hypocyphtini may provide critical insights into the evolution of the tergal gland, but to date, their chemistry and glandular anatomy are unexplored beyond confirming that they possess a solvent cell reservoir—the basis for their placement in Higher Aleocharinae (*47, 59*). Hypocyphtini is an enigma, however, in that its known members are specialist mite predators, some performing important roles in the biological control of pest mite species (*87, 88*). This specialized biology contrasts with the generalist predatory lifestyle that is thought to be ancestral in Aleocharinae. Morphologically, the beetles are also divergent, with a minute, compact body with a short abdomen (**Fig. 6A–C**). Due to the key phylogenetic position of Hypocyphtini, we assembled draft genomes and profiled the secretions of members of three genera that cover the tribe’s diversity: *Cypha*, *Oligota* and *Holobus* (**Fig. 3A**).

**Figure 6.**
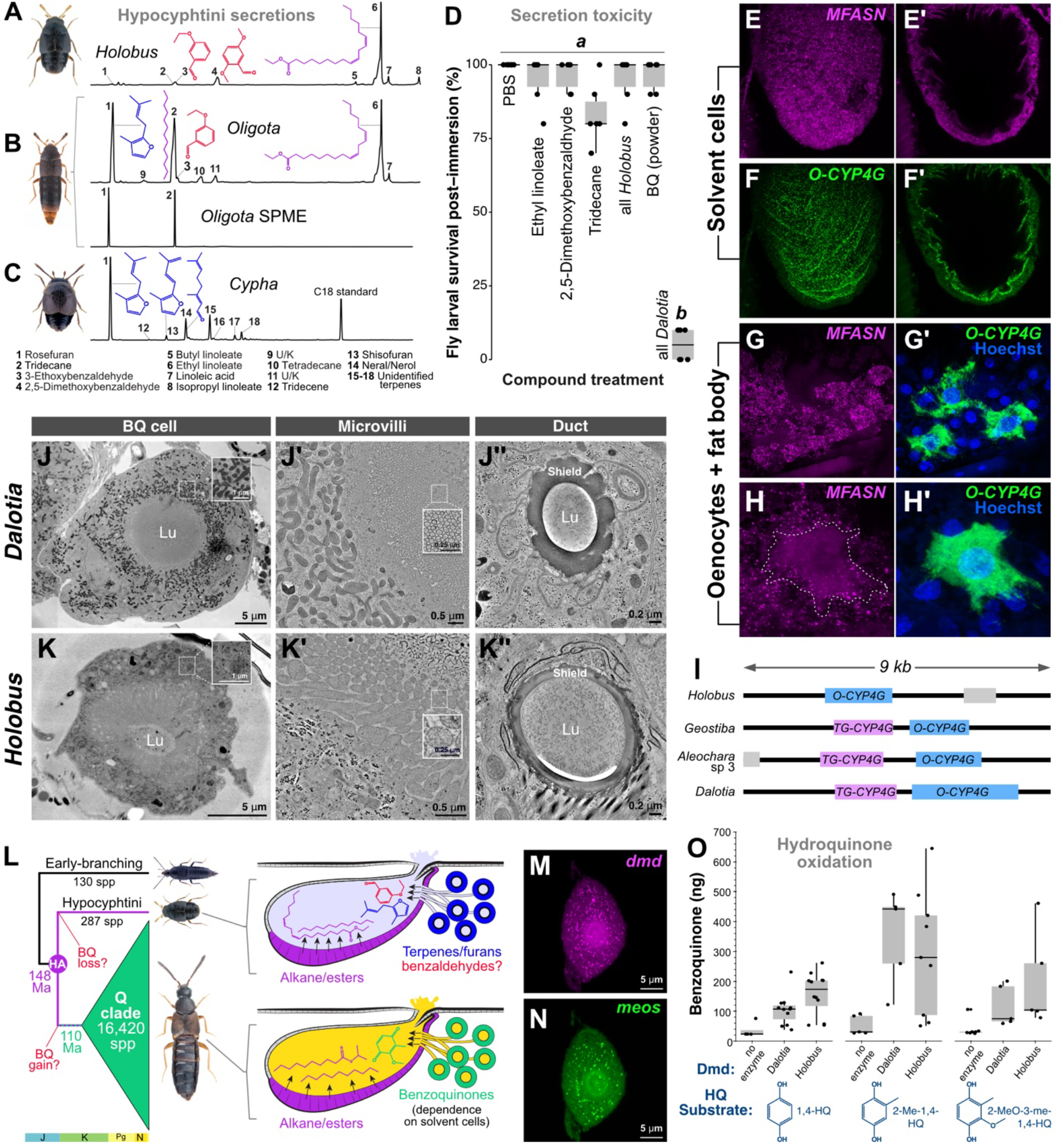
Glandular biology of the earliest-branching HA lineage. **(A-C)** GC traces of glandular compounds from hypocyphtines: *Holobus* **(A)**, *Oligota* **(B, B’)** and *Cypha* **(C). B’** shows headspace volatiles from 20 *Oligota* beetles detected via single-phase microextraction (SPME). **(D)** Survival of *Drosophila* larvae following immersion in synthetic hypocyphtine or *Dalotia* secretions. **(E, F)** HCR labeling of *MFASN* **(E, E’**, magenta) and *O-CYP4G* **(F, F’**, green) transcripts in region of *Oligota* solvent reservoir **(E, F** show labelling within plane of solvent cell epithelium; **E’**, **F’** show cross section through reservoir epithelium). **(G, H)** HCR labelling of *MFASN* (magenta) and *O-CYP4G* (green) in *Oligota* fat body and oenocytes (blue: Hoechst-stained nuclei). **(J, K)** TEM micrographs of *Dalotia* (top) and *Holobus* (bottom) BQ cells. Lu: lumen. Insets in **J** and **K** show differing mitochondrial densities between the two species (electron-dense structures). **J’** and **K’** show differing microvillar organization and density within BQ cell lumens. **J’’** and **K’’** show differing shield thickness within the internal lumen (Lu) of duct cells. **L:** Phylogenetic topology of deepest divergences in Aleocharinae showing relative species richness of major clades, and the respective chemistries of the Hypocyphtini and Q clades that comprise the HA. Alternative scenarios posit that benzoquinones were either gained in the Q clade or lost in Hypocyphtini. **(M, N)** Localization of *dmd* (**M**, magenta) and *meos* (**N**, green) expression in *Oligota* BQ cells. **(O)** Dmd of *Holobus* can convert hydroquinone substrates (HQs) to the corresponding benzoquinones at comparable efficiency to *Dalotia* Dmd.

All three beetles produce an alkane (tridecane) or its corresponding alkene (tridecene) (**Fig. 6A–C**). Further, two genera produce a long-chain fatty acid, linoleic acid, and ester derivatives thereof, revealing conservation of fatty acid-derived solvents across the HA clade (**Fig. 3B**). Using Hybridization Chain Reaction (HCR) in a species of *Oligota*, we labeled abdominal cells expressing the solvent pathway fatty acid synthase, *MFASN*, revealing expression in both solvent cells and fat body (**Fig. 6E, G, H**), mirroring the pattern in *Dalotia* (**Fig. S9E**) (*30*). We infer that *MFASN* was co-opted into solvent cells in the HA stem (rather than the Q clade stem) and has been functionally maintained in this role as the HA radiated throughout the Cretaceous and Cenozoic (**Fig. 4H**). In contrast to the conserved function of *MFASN*, however, the terminal decarbonylase for alkane synthesis, *TG-CYP4G*, is absent in hypocyphtine genomes (**Fig. 6I**). *TG-CYP4G* is thus an apparent Q clade novelty (**Fig. 4H**). *TG-CYP4G* is a duplicate of an ancient cytochrome P450 that is broadly conserved in insects and functions in oenocytes, where it decarbonylates very long chain aldehydes to produce alkane and alkene cuticular hydrocarbon pheromones (CHCs) (*89, 90*). This more ancient gene copy, *Oenocyte-CYP4G* (*O-CYP4G*), is conserved across the Coleoptera (**Fig. S9B**). Remarkably, in hypocyphtines, it is *O-CYP4G* that is expressed in solvent cells, suggesting that the enzyme was first co-opted from oenocytes prior to the duplication event that yielded *TG-CYP4G* (**Fig. 6F**). We examined sequence evolution within *TG-CYP4G* and found that following its origin via duplication, *TG-CYP4G* experienced episodic selection, with 12 codons showing signatures of positive selection and three others showing relaxed selection within the Q clade (aBSREL ω_2_=8.32, LRT=18.74, p<0.001; codeml ω_2_=999, p<0.001) (**Fig. S9B**). We infer that *O-CYP4G* was co-opted into solvent cells in the BQ clade stem lineage; the enzyme functioned pleiotropically in both CHC and defensive alkane synthesis—a situation preserved in hypocyphtines. Subsequently, the gene duplicated in the Q clade stem, yielding *TG-CYP4G*, which underwent neofunctionalization to generate a solvent cell-specific enzyme that replaced *O-CYP4G*. The tandem syntenic arrangement of *O-CYP4G and TG-CYP4G* has been conserved across the Q clade (**Fig. 6I**).

The most remarkable feature of the hypocyphtine secretions is, however, the complete absence of benzoquinones. Instead, all three beetles secrete a novel compound class in the form of a furan, rosefuran (**Fig. 6A–C**). Further, *Cypha* also produces monoterpenes (the compound class from which the furan is likely derived). Additionally, both *Holobus* and *Oligota* produce benzaldehydes—compounds unseen in other aleocharines. We relate these chemical novelties to the specialized acariphagous biology of hypocyphtines. Rosefuran is a mite sex pheromone (*91*); the monoterpene neral is a mite attractant or alarm pheromone (*92, 93*); while the use of benzaldehydes as pheromones is also widespread in mites (*92, 94–96*). Consequently, we propose that hypocyphtines possess modified tergal gland chemistries that adapt them for mite predation. Conversely, chemical defense seems unlikely. The furan and terpenes lack pronounced toxicity or irritant properties, and nor do the benzaldehydes, which we tested by immersing *Drosophila* larvae in 2,5-dimethoxybenzaldehyde—a compound produced by *Holobus*. Larval immersion in 2,5-dimethoxybenzaldehyde caused no reduction in survival when applied either alone or when mixed with the specific alkane and ester that *Holobus* beetles produce (**Fig. 6D**). This contrasts with potent lethality caused by immersion in a synthetic *Dalotia* tergal gland secretion (**Fig. 6D**). We note also that the reduced abdominal mobility of hypocyphtines likely precludes them from directly smearing secretions on other organisms—the mode of deployment in many aleocharines (*33, 35*). Sampling volatiles in the headspace above *Oligota* beetles, we detected strong secretion of rosefuran and tridecane but no linoleic acid derivatives, which appear not to be volatilized (**Fig. 6B**, lower trace). We hypothesize that volatile release of the rosefuran may provide chemical camouflage during predation, or act as a chemical lure for prey.

We examined the cellular ultrastructure of the tergal gland using electron tomography and confirm that hypocyphtines possess BQ cells, which appear homologous to those of *Dalotia*. *Dalotia* BQ cells are extremely large (∼30 µm diameter) spherical acini, with a hollow lumen formed by involution of the apical cell membrane (**Fig. 6J).** A dense network of narrow, apical microvilli extends into the lumen, presumably secreting hydroquinones together with BGLU and Dmd for cleavage and oxidation to benzoquinones (**Fig 6J’**). Connected to each BQ cell is a long, convoluted duct cell, enveloping a lumen with a thick, protective shield for channeling benzoquinones into the gland reservoir (**Fig. 6J’’**). Although the BQ cells of the hypocyphtine *Holobus* are smaller (∼15 µm diameter), they share this overall anatomy (**Fig. 6K**). Both the solvent cells and BQ cells are thus synapomorphies of the HA, dating to the MRCA of this vast clade at the Jurassic-Cretaceous boundary (**Fig. 3A**). The BQ cells of the two beetles nevertheless differ in certain key aspects. Most striking is their differing mitochondrial content: *Dalotia* BQ cells are extremely rich in mitochondria, consistent with a high demand for hydroquinone synthesis by these organelles (**Fig. 6J, inset**). Conversely, *Holobus* BQ cells lack such abundant mitochondria (**Fig. 6K**). We propose that the BQ cells of hypocyphtines do not synthesize benzoquinones but are probably responsible for producing some or all of the novel, non-fatty acid derived compounds these beetles secrete, such as furans, terpenes and benzaldehydes. We speculate that other ultrastructural differences of the BQ cells may correspond to the reduced need for protection from cytotoxicity: their lumenal microvilli are much thicker and less rigidly organized (**Fig. 6K’**), and the duct cell lumen is also wider and less heavily protected (**Fig. 6K’’**).

Due to the minute size of hypocyphtines (**Fig. S12E**), we have been unable to dissect their tergal gland cells for single cell transcriptomics, so the pathways these cells express remain unknown. However, the apparent lack of benzoquinone production by hypocyphtine BQ cells raises a more fundamental question about the ancestral biosynthetic function of this cell type (**Fig. 6L**). Their specialized secretions imply that hypocyphtines may have sacrificed benzoquinones in favor of furan/terpene/benzaldehydes (**Fig. 6L**). Curiously, however, we find two marker genes of benzoquinone synthesis—dmd and meos—are present in the genomes of hypocyphtines (**Fig. 5G**). Using HCR, we also observe clear expression of both loci within the BQ cells of *Oligota* (**Fig. 6M, N**). Further, comparison of abdominal segments using bulk RNAseq revealed 360 differentially expressed genes upregulated in the gland-bearing segment of *Holobus*, of which 16 orthologs were also enriched in *Dalotia*’s gland-bearing segment (**File S1**). Of these, *dmd, ATP7,* and *BGLU* are upregulated in the gland-segment of *Holobus* (**File S1**).

We asked whether hypocyphtine dmd orthologues encode functional proteins by synthesizing and purifying a hypocyphtine Dmd and testing its ability to oxidize hydroquinones *in vitro* (**Fig. 6O**). Curiously, the hypocyphtine enzyme functions equivalently to *Dalotia* Dmd, confirming that the protein is functionally intact in hypocyphtines, despite these beetles not producing benzoquinones. The lack of pseudogenization of Dmd implies that the enzyme performs a different role, either within the BQ cells themselves or possibly elsewhere in the beetle. While it remains plausible that benzoquinones were gained along the Q clade stem, we favor the view that it arose in the HA clade stem, and that hypocyphtines have secondarily lost benzoquinone synthesis (**Fig. 6L**). Hypocyphtine chemistry appears closely linked to the derived acariphagous diet of these beetles, and is thus unlikely to represent the primitive condition in the HA. We suggest that the continued expression of *dmd* and *meos* in BQ cells may constitute a ’molecular spandrel’ that is evidence of the cell type’s prior function in benzoquinone production (*97*, *98*; see discussion). Consequently, the functions of both the solvent and BQ cell types—and their cooperative interaction that yields the BQ/FA cocktail (*32*)—may have been present in the MRCA of the HA clade, 148 Ma.

### Evolvability of tergal gland cell types under symbiosis

Hypocyphtine secretions reveal how the tergal gland has been a substrate for the evolution of specialized chemical ecological interactions in Aleocharinae. Chemical innovation in aleocharines is well known for being taken to the extreme in many symbiotic lineages that have evolved to live inside colonies of ants and termites. A large body of literature has documented the chemical interactions between symbiotic aleocharines and their hosts (*36–38, 40, 41, 45, 46, 99–104*). In numerous cases, interactions are mediated by abdominal glandular secretions that confuse, pacify or appease workers, or elicit the beetle’s adoption into the nest. Different symbiotic lineages have been hypothesized to repurpose the tergal gland to produce such host-manipulating secretions, implying potential biosynthetic reprogramming of the BQ and/or solvent cell types (*41, 44–46*). We sought to illuminate how the transition to a symbiotic lifestyle is manifested molecularly in the BQ and solvent cell types. In pursuing this question, we discovered an instance of dramatic modification of tergal gland chemistry in the myrmecophile *Liometoxenus*—a genus described relatively recently, for which no prior chemical, behavioral or genomic data existed (*57*) (**Fig. 7A**). We routinely find *Liometoxenus* inhabiting colonies of *Liometopum occidentale* ants in the Angeles National Forest (CA: LA County) and have observed that the beetles execute a remarkable behavioral interaction with host workers. Rather than deploying a noxious chemical defense, the beetle secretes a volatile cocktail that acts at a distance to intoxicate ants, impairing their locomotion and attenuating their aggression towards the beetle (**Video S1**). In addition to overriding host aggression, the secretion subdues ants, enabling *Liometoxenus* to feed upon workers.

**Figure 7.**
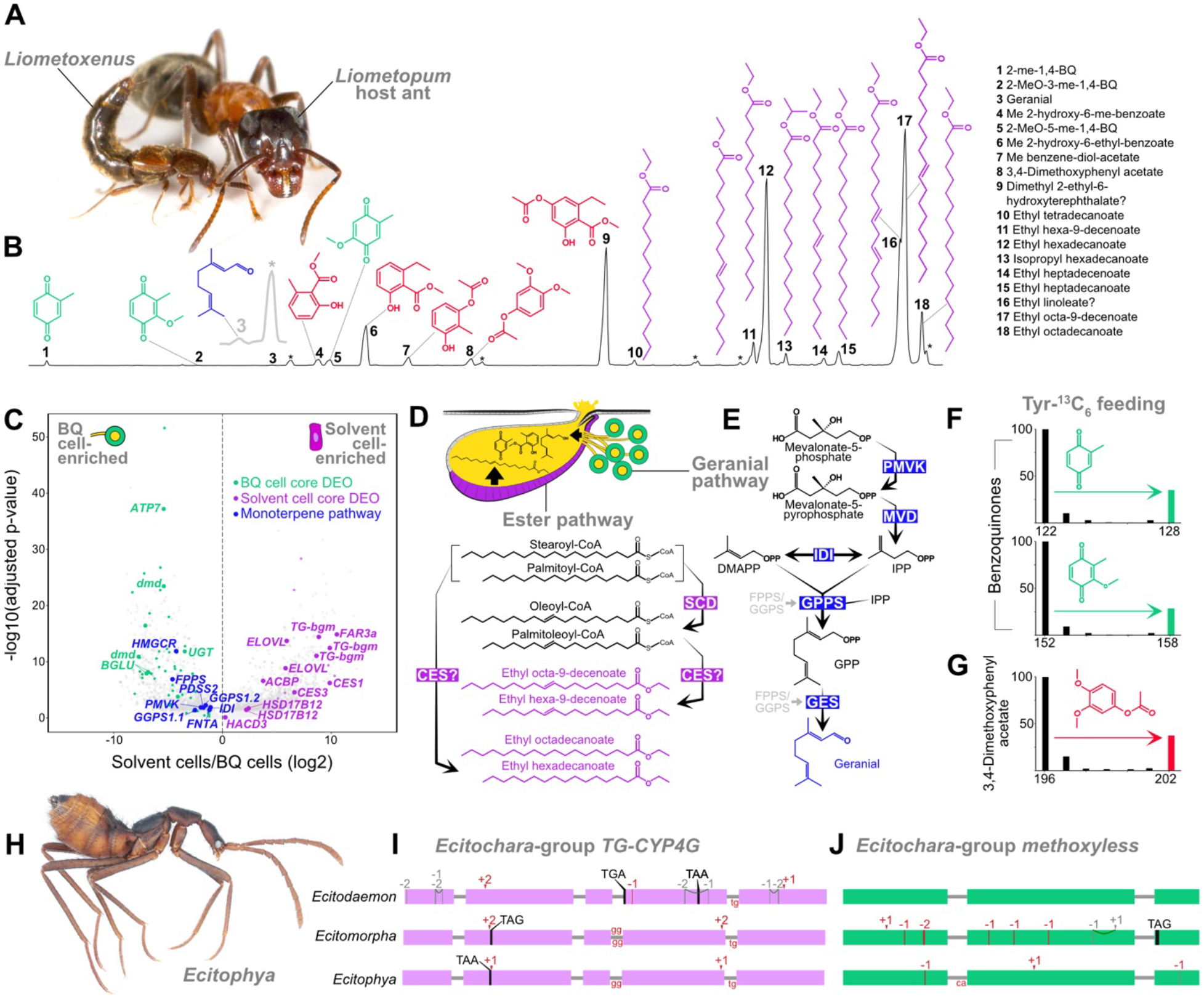
Tergal gland evolution in myrmecophiles. **(A)** *Liometoxenus newtonarum* beetle and *Liometopum occidentale* host ant. Photo credit: David Miller. **(B)** GC trace of *Liometoxenus* gland compounds. Magnified region of geranial peak (compound 3) is shown in grey. Asterisks mark contaminants. **(C)** Volcano plot of DEGs between the solvent cells (positive log_2_ fold-change) and BQ cells (negative log_2_ fold-change) for *Liometoxenus.* DEOs of biosynthetic genes from Figure 3 are colored (solvent cell= purple, BQ cell= green) and key biosynthetic genes are labeled, including the inferred monoterpene pathway expressed in BQ cells (blue). *IDI*: *isopentenyl-diphosphate delta-isomerase 1*; *FPPS*: *farnesyl pyrophosphate synthase*; *HMG-CoA*: *3-hydroxy-3-methylglutaryl-coenzyme A reductase*; *PMVK*: *phosphomevalonate kinase*; *FNTA*: *farnesyltransferase/geranylgeranyltransferase type-1 subunit alpha*; *GGPS1_1*/*GGPS1_2*: *geranylgeranyl pyrophosphate synthase*; *PDSS2*: *decaprenyl-diphosphate synthase subunit 2*; *SCD5.2*: *stearoyl-CoA desaturase 5*; *CES1*: *Carboxylesterase 4A*; *ACBP*: *Acyl-CoA-binding protein homolog*; *CES3*: *type-B carboxylesterase*. **(D)** Cartoon of tergal gland showing hypothesized pathway for *Liometoxenus* aliphatic esters. SCD: Acyl-CoA Delta(11) desaturase. "CES" denotes the hypothesized function of either or both the Carboxylesterase 4A (CES1) or the type-B carboxylesterase (CES3) that are upregulated in *Liometoxenus* solvent cells. (**E)** Inferred terpene pathway leading to the synthesis of geranial. See part **C** for protein acronyms. **(F, G)** Representative mass spectra of molecular ion regions of specific compounds from *Liometoxenus* fed with dead ants infused with ^13^C_6_-Tyr. Spectra were recorded in single-ion mode. **(F)** Spectra for 2-methyl-1,4-BQ (MW 122) and 2-methoxy-3-methyl-1,4,-BQ (MW=152) reveal a strong [M+6]^+^ enrichment for both compounds due to ^13^C_6_ incorporation (green bars). **(G)** Spectrum for 2-hydroxy-6-methyl-benzoate showing strong [M+6]^+^ enrichment due to ^13^C_6_ incorporation (red bar). **H:** *Ecitophya simulans* beetle. **I, J:** Gene models of *TG-CYP4G* and *methoxyless* from three *Ecitochara*-group species, showing inactivating mutations. Negative/positive numbers are frameshift base pair deletions/insertions against the reference genome (*Dalotia*). Premature stop codons are shown, and splice junction mutations are shown at intron-exon boundaries.

We profiled the tergal gland chemistry of *Liometoxenus* and found a complex cocktail containing 18 compounds spanning multiple classes: long- and medium-chain aliphatic esters including both saturated and unsaturated species; low levels of benzoquinones identical to those found in free-living species; a long series of aromatic esters, including some in high abundance; and low amounts of a single terpene, geranial (**Fig. 7B**). To our knowledge, this secretion is the most diverse chemical mixture produced by a single rove beetle species. Its behavioral effects on ants imply an adaptive role in the beetle’s symbiotic lifestyle. To illuminate how *Liometoxenus* evolved such a complex secretion, we sequenced and assembled the genome of *L. newtonarum*, and created single-cell transcriptomes for both the BQ and solvent cells via SMART-Seq. We find evidence for dramatic biosynthetic pathway evolution in *Liometoxenus* tergal gland cell types (**Fig. 7C**). First, we find that an entire monoterpene synthesis pathway has been recruited into BQ cells, presumably leading to the low concentration of geranial in the secretion. Enzymes for every step from the mevalonate-5-phosphate precursor to geraniol pyrophosphate are expressed in BQ cells (**Fig. 7C**). While we cannot identity with certainty the terminal geranial synthase (GES), the BQ cells express *Liometoxenus*-specific duplicates of both FPPS and GPPS—enzymes that in other terpene-producing insects such as butterflies, hemipterans and outgroup beetles have convergently duplicated and been neofunctionalized to create terpene synthases (*105–107*) (**Fig. S14A, B**). We hypothesize that a parallel scenario has occurred in *Liometoxenus*.

Synthesis of benzoquinones appears to be largely identical to other aleocharines, with expression of many core components conserved in *Liometoxenus* BQ cells (which also express an additional, *Liometoxenus*-specific *dmd* duplicate) (**Fig. 7C**). However, the molecular precursor of the benzoquinones—tyrosine—appears to have become strongly biased towards the synthesis of new compounds. Feeding *Liometoxenus* adults Tyr-^13^C_6_, we observed strong ^13^C incorporation into the benzoquinones, their molecular weights increasing by +6, confirming the benzene rings are derived from aromatic amino acids (**Fig. 7F**), as in *Dalotia* (*32*). However, in addition to the benzoquinones, we saw significant Tyr-^13^C_6_ incorporation into some of the aromatic esters, which now dominate the secretion (**Fig. 7G**). Like the benzoquinones, these compounds are thus not directly sequestered from the beetle’s diet, nor are their benzene rings synthesized *de novo* by the beetle. Instead, their synthesis occurs via a novel pathway, parallel to the benzoquinone pathway, but fueled by the same dietary metabolite. Presently, we cannot speculate over how *Liometoxenus* creates such a variety of esterified compounds; no obvious candidate enzymes for these cyclic compounds emerge from the transcriptomes of either the BQ or solvent cells.

Unlike the solvents in *Dalotia* and *Aleochara*, headspace sampling reveals that the longer chain length esters of *Liometoxenus* are non-volatile (**Fig. S15A**). The esters appear instead to act as a solvent contained within the reservoir from which the remaining compounds volatilize, influencing host ant behavior from a distance. This unusual solvent fraction arises from a pathway that appears non-homologous to that of other aleocharines (**Fig. 7D**). The precursors of the solvent compounds are likely palmitic and stearic acid (C16 and C18)—amongst the most common free fatty acids in insects, and which are made via lipogenesis in the fat body (*108, 109*). However, we hypothesize that additional synthesis within the solvent cells is likely given the strong expression of key enzymes that drive the fatty acid elongation cycle (**Fig. 7C**), which we speculate may underlie the range of different chain length compounds in the secretion. Unlike the *Dalotia* ester pathway, *Liometoxenus* does not employ an ɑ-esterase to convert these fatty acids to esters; instead, carboxylesterases of alternative families are upregulated in solvent cells, and likely carry out esterification (**Fig. 7C, D**; **Fig. S11**). Most esters are present in both unsaturated and desaturated forms (**Fig. 7B**), and we find that *Liometoxenus* solvent cells strongly express a homologue of the canonical metazoan stearic/palmitic acid desaturase, SCD (stearoyl-CoA desaturase) (**Fig. 7D**).

*Liometoxenus* uses its tergal gland secretion to manipulate worker behavior, but appears not to engage in complex social interactions with its host ant. In the most specialized, highly integrated symbionts, however, noxious defenses appear less critical as the beetles evolve social behaviors and chemical mimicry strategies that assimilate them into the colony’s social structure. In several such taxa, the tergal gland has evolutionarily degenerated (e.g. *56*, *110*, *111*). In our phylogenomic sampling, we included members of one such clade—the *Ecitochara* group, represented by the genera *Ecitophya*, *Ecitomorpha* and *Ecitodaemon* (*56*). These beetles are morphological ant mimics (**Fig. 7H**), which are accepted into nomadic colonies of *Eciton* army ants (*56, 112*). As a first glimpse into how such an integrated lifestyle impacts genomic and phenotypic evolution, we analyzed the evolution of the BQ and solvent cell core transcriptome in these beetles. Consistent with most core loci having been co-opted from roles outside of the tergal gland, there is strong evidence that 35 to 38% of the 554 BQ and solvent cell type core loci remain present (intact or partially intact) in the genomes of these myrmecophiles. While a large number of core loci were missing or partially missing (21-33%) due to the fragmentation of the ecitocharine assemblies, we found clear evidence of pseudogenization and gene loss in 13, 10 and 12 of the core biosynthetic genes from *Ecitodaemon*, *Ecitomorpha* and *Ecitophya*, respectively (**Table S6**). Multiple inactivating mutations, including frameshifts and premature stop codons, have accumulated in both the solvent cell terminal carbonylase, *TG-CYP4G*, and the benzoquinone-modifying enzyme, *methoxyless* (**Fig. 7I, J**).

Such a pattern of gene inactivation is consistent with relaxed selection on these loci— presumably a consequence of degeneration of the now-obsolete tergal gland. We note that pseudogenization may have occurred independently in crown-group lineages rather than along the *Ecitochara*-group stem. Specific inactivating mutations are often not shared by all three taxa, with only *TG-CYP4G* of *Ecitophya* and *Ecitomorpha* sharing a subset of changes (**Fig. 7I**). Moreover, *Ecitodaemon* appears to still possess an intact *methoxyless* gene (**Fig. 7J**). Given that the three genera share a MRCA ∼24 Ma (95% HPD: 33–15 Ma; **Fig. 3A**), which presumably already lacked the function of the tergal gland, these idiosyncratic patterns of gene inactivating mutations implies a surprisingly slow rate of coding sequence decay in these myrmecophiles. Alternatively, gland loss may be a recent, convergent phenomenon across the three genera. All three species also possess an apparently intact orthologue of the benzoquinone pathway laccase, *dmd* (**Fig. 5G**). We synthesized and purified Dmd from *Ecitophya* and tested its ability to oxidize hydroquinones in *vitro*. *Ecitophya* Dmd is functional (**Fig. S15B**); hence, we posit that this enzyme plays an alternative role in these myrmecophiles. This result parallels our findings in the Hypocyphtini, which similarly possess an intact Dmd despite lacking benzoquinones (**Fig. 6O**).

## Discussion

The radiation of Metazoa’s largest family, Staphylinidae, has been coupled to pervasive biochemical innovation, precipitated by widespread convergent evolution of abdominal exocrine glands. Underlying this phenomenon has been a diversification of taxon-restricted secretory cell types that express unique, small molecule biosynthesis pathways. Here, we examined the evolution of one such structure—the tergal gland of Aleocharinae. By combining genomic, single-cell transcriptomic and chemical data from across the aleocharine phylogeny, we uncovered evolutionary changes at the genome, pathway and cell type levels that underlie the gland’s assembly in early aleocharines, its deep functional conservation as the beetles radiated globally, and its potential for evolvability via biosynthetic repurposing—a process that has chemically adapted aleocharines to specialized niches. The origin of the tergal gland and its seeming versatility have facilitated novel ecologies and behaviors (*31, 37, 44–46*). Our findings underscore how new organismal properties can derive from the *de novo* evolution of cell types, with ramifications at the macroevolutionary scale.

### Genomic contingencies and modular evolution of gland cell types

We inferred that two taxon-restricted cell types comprising the tergal gland—the solvent and BQ cells—arose early in aleocharine evolution, along the HA stem lineage. The functional assembly of these cell types created a cooperative biosynthetic system where benzoquinone toxicity is unlocked by a fatty acid-derived solvent (the BQ/FA cocktail). Following the origin of this novel organ, its two constituent cell types and their secretions have been broadly conserved across most of the HA—numbering tens of thousands of extant lineages that began diversifying in the Early Cretaceous. We have presented evidence that both the BQ and solvent cells express deeply conserved core transcriptomes that contribute to each cell type’s biosynthetic function. These core transcriptomes comprise a majority of phylogenetically ancient, co-opted loci that pre-date Aleocharinae, as well as more recent paralogues that originated along stem lineages of either the HA or Q clades, including key benzoquinone and solvent pathway enzymes. A major expansion of laccase enzymes along the HA stem was a decisive genomic contingency, yielding the terminal hydroquinone oxidase, *Decommissioned*. Similarly in the HA stem, the birth of the *methoxyless* gene within an expansion of *COQ3* loci enabled a hydroxylation of these toxic compounds that has been widely conserved across benzoquinone-producing taxa (**Fig. 3A**) (*31*). Gene co-option in novel cell types has been hypothesized as a driver of gene duplication, which permits escape from pleiotropic conflict (*76*). Such a scenario may explain the origin of taxon-restricted loci expressed in the BQ and solvent cells. The evolution of *TG-CYP4G* provides a clear case where co-option of an ancestral enzyme into solvent cells occurred prior to its duplication, which in turn permitted the neofunctionalization of the duplicate copy in this novel cell type.

Notably, we found that conservation of the core BQ and solvent cell transcriptomes across the Q clade has occurred despite lineage-specific divergence in the remaining constitutive fraction of the transcriptome. Further, in the case of solvent cells, we found that a major expression program for fatty acid synthesis and modification, OVFB-GEP, has been broadly conserved in solvent cells across the Q clade, while a co-expressed program that confers cuticular identity (CC-GEP) exhibits much weaker conservation. These patterns imply that long-term stabilizing selection on defensive chemistry has occurred almost clade-wide across the higher Aleocharinae. At the cellular level, this is reflected in the conservation of the BQ and solvent cells and their cooperative, biosynthetic interaction, manifesting in the relative evolutionary stasis of the differentially expressed portion of each cell type’s transcriptome that confers cellular identity and function. Simultaneously, the remainder of the transcriptome has experienced substantial change, presumably reflecting varying selection regimes in different taxa. These results indicate the modular nature of transcriptome evolution, where expression programs that confer different subfunctions within cell types appear relatively decoupled from each other evolutionarily. The origin of the solvent cell itself—inferred to have arisen via co-option of OVFB-GEP into intersegmental cells of the beetle’s abdomen (*32*)—further underscores the role of transcriptome modularity in tergal gland cell type evolution.

### Ecological specialization through cell type evolvability

The broad conservation of aleocharine defensive chemistry has not precluded dramatic evolutionary modification of the biosynthetic output of both the solvent and BQ cells. In the case of the solvent cells, the specific fatty acid derivatives can vary extensively, with predicted effects on the secretion’s physicochemical properties. Across the phylogeny, we find substantial variation in the functional groups, numbers, ratios, carbon chain lengths and degrees of saturation of these compounds. These changes arise from several mechanisms, including the differential recruitment of esterases and desaturases into solvent cells, and the presumed loss of FAR and CYP4G activities in taxa that have become reliant on esters. Variation in precursor fatty acid chain length appears likely, as well as extension of fatty acids via elongase activity. While the functions of these modifications remain to be examined, we suggest that some of them may reflect adaptive streamlining of the secretion—in particular, refinements of its viscosity and surface coating ability (wetness). Differences in chain length, functional group, and likely other types of covalent modification can profoundly alter these properties, which are themselves temperature-dependent (*32, 66, 67*). Such streamlining may enable production of a functional secretion despite microclimatic differences in the niches different taxa occupy. Physicochemical streamlining may also permit different modes of gland deployment (e.g. directly smearing the secretion onto targets versus volatilizing the secretion to act at a distance).

In contrast to the variation in fatty acid-derived solvents, aleocharines produce only a small number of benzoquinone species, potentially as few as four across the entire subfamily (**Fig. 3B**) (*31*). Nevertheless, the BQ cell has proven highly modifiable, permitting aleocharines to secrete new compound classes. In the earliest-branching HA tribe, Hypocyphtini, benzoquinones have been lost, but BQ cells appear to have been repurposed for synthesis of other compound—furans, terpenes and benzaldehydes. Production of these compounds, which include known mite pheromones, is likely an adaptation to the specialized acariphagous lifestyles of this tribe. Most dramatically, we found a high complexity secretion in the myrmecophile *Liometoxenus* that attenuates worker aggression and facilitates infiltration of host colonies. The anatomies of the BQ cells and their associated ducts appear tailored for controlled production of cytotoxic benzoquinones. It is unclear how the synthesis, intracellular trafficking and secretion of other types of compounds can evolve within such a pre-existing cellular environment. Although the subcellular regulation of small molecules inside animal cells is extremely poorly understood (*76*), the BQ cell type’s evolutionary versatility implies that certain cellular anatomies are efficient for synthesis and secretion of multiple compound types. Our finding that aleocharines employ a plant-like system of toxin regulation involving secretion and cleavage of hydroquinone glycosides suggests a broadly versatile mechanism that may be co-opted for production of other compounds.

### Regulatory basis of evolvability

How cell types gain new multi-enzyme pathways presents a conundrum, since a battery of loci must become co-expressed within the same cell simultaneously to render them all—and the compounds they produce—visible to natural selection. For the synthesis of geranial, for example, we uncovered an entire monoterpene pathway in the BQ cells of *Liometoxenus*. How this occurred mechanistically is unknown, but we suggest it could arise from co-regulation of multiple loci by an upstream "terminal selector" transcription factor (*113*). We previously proposed that two abdominal Hox proteins, Abdominal A and Abdominal B, may have recruited OVFB-GEP into cuticle cells, creating the solvent cell type (*32*). Both Hox proteins are needed for BQ and solvent cell development (*114*), and also remain active post-differentiation. In this context, we speculate that these Hox proteins may play governing roles in recruiting novel enzymes, pathways, or larger expression modules into the BQ and solvent cells, either directly or via co-option of intermediate transcription factors that regulate biosynthetic programs in other cell types.

A perplexing finding is the continued transcription of enzymes that functioned ancestrally within tergal gland cell types but no longer apparently influence biosynthesis. Most strikingly, we detected *dmd* and *meos* transcripts in hypocyphtine BQ cells, despite the inferred loss of benzoquinone synthesis in these beetles. Although these enzymes may perform other roles within this cell type, we suggest their persistent expression may also derive from enhancer pleiotropy: regulatory elements that drive expression both in the tergal gland and in other organismal contexts where their gene products are visible to natural selection. The costs of continued transcription of loci encoding past functions may be relatively low and outweighed by selection to maintain expression elsewhere. That Dmd orthologues from the hypocyphtine *Holobus* and the myrmecophile *Ecitophya* are functionally intact indicate involvement of this laccase elsewhere in aleocharine metabolism, or possibly in the production of benzoquinones in larvae—a life stage that in some other aleocharines has been shown to produce a BQ/FA cocktail from a developmentally distinct abdominal gland (*115*).

### Cell type evolution of a key innovation

The inordinate diversity of beetles is thought to have been contingent on the evolution of protective elytra (*6, 10–12*). Paradoxically, the largest and most ecologically diverse beetle family has partially forsaken this trait, reducing elytron size to expose a soft, unprotected abdomen. Staphylinid cladogenesis may, ironically, have hinged on this loss of physical protection—the flexible abdomen proving a versatile substrate for an alternative mode of protection in the form of targetable defensive glands. The evolution of novel cell types comprising peripheral structures such as exocrine glands can profoundly modulate the interaction between an organism and its environment (*76*). Analogous to the origin of photoreceptors (*116*) or defensive nematocysts (*117*), the tergal gland may be a more recent example where the *de novo* evolution of cell types has enabled a clade to enter many new adaptive zones. Through chemical and antimicrobial defense, the gland has bought aleocharines enemy-free-space to colonize and diversify throughout Earth’s terrestrial ecosystems. As a reprogrammable device, the gland has enabled aleocharines to evolve specialized ecological relationships with other species. Such direct connections between the tergal gland and Aleocharinae’s numerical and ecological diversity implicate this structure and its two cell types as a key innovation behind one of Coleoptera’s most successful radiations.

## Supporting information

File_S1_expression_results

## Acknowledgements

We thank Charlie Barnes, Mike Caterino, Munetoshi Maruyama, Thomas Schmitt and Christoph von Beeren for beetle specimens used for genome sequencing. We are grateful to Taku Shimada for *Dalotia* photography and Udo Schmidt for the use of specimen images. This work was funded by grants to J.P. from the National Institutes of Health (1R34NS118470-01), National Science Foundation (2047472 CAREER), as well as support from a Shurl and Kay Curci Foundation grant, a Rita Allen Foundation Scholarship, a Pew Biomedical Scholarship, and an Alfred P. Sloan Fellowship. Additional funding was provided by Iridian Genomes (IRGEN_RG_2021-1345 Genomic Studies of Eukaryotic Taxa), and a Gerstner Fellowship in Bioinformatics and Computational Biology to M.L.A. This work was also supported by the Millard and Muriel Jacobs Genetics and Genomics Laboratory at California Institute of Technology. A.B. was a Simons Fellow of the Life Sciences Research Foundation (LSRF). J.M.W is a National Science Foundation Graduate Research Fellowship recipient.

## Author contributions

Conceptualization: J.P., M.L.A. and S.A.K. Methodology: J.P., M.L.A. and S.A.K. Investigation: S.A.K., T.H.N., A.B., M.S.L., S.Q., J.M.B., J.M.W., D.R.M., M.Y., I.A.A., S.P., M.G., S.R.D., M.L.A. and J.P. Formal analysis: S.A.K., T.H.N., and A.B. Data Curation: S.A.K. Writing – Original Draft: J.P. and S.A.K. Writing – Review & Editing: J.P. and S.A.K. Supervision: J.P. Project Administration: J.P. Funding Acquisition: J.P.

## Materials and Methods

### DNA extraction and short and long-read sequencing

*Dalotia* were collected from an inbred population (original source: Applied Bionomics, Canada) reared in the lab as described previously (*114*). Other taxa were collected from various locations or donated to this study (see **Table S3**). For Illumina sequencing, DNA was isolated from a single specimen, with the exception of *Cypha longicornis*, *Holobus* sp. and *Oligota* sp. with two, five and seven specimens, respectively. We used either a non-destructive extraction method described by Maruyama and Parker (*42*) or a complete tissue homogenization with the Qiagen DNeasy Blood and Tissue extraction kit (Qiagen, Germany) following the manufacturer’s protocol with slight modifications as follows. Tissue was homogenized in the ATL lysis-proteinase K solution and incubated for 4 h or overnight at 56 °C. The tissue solution was incubated in RNaseA (Qiagen, Germany) for 2 min followed by the manufacturer’s protocol. Two rounds of DNA elution were performed with 100 µl warmed elution buffer (50 °C) each round. DNA was concentrated using the Monarch PCR and DNA Cleanup kit (New England Biolabs, MA) with warmed elution buffer. DNA quantity was assessed using the Qubit High Sensitivity dsDNA kit (Thermo Scientific, MA) and DNA integrity was assessed visually with gel electrophoresis. To complement field identifications, we also amplified fragments of *cytochrome c oxidase subunit 1* and *18S* rRNA for each specimen according to Maruyama and Parker (2017). PCR products were purified using ExoSAP-IT (ThermoFisher, MA) and sequenced by Laragen (Culver City, CA). Illumina paired-end sequencing libraries were prepared using the Illumina TruSeq DNA (Illumina, CA) or NEBNext Ultra FS DNA library kits (paired-end 150bp reads, average insert size 155 ± 105 bp, New England Biolabs, MA)) following the manufacturer’s protocol, quantified with Agilent Bioanalyzer High Sensitivity DNA kit (Agilent Technologies, CA) and sequenced on various Illumina platforms by Iridian Genomics, Macrogen (now Psomagen), Fulgent Genetics, Genewiz, and the Millard and Muriel Jacobs Genetics and Genomics Laboratory at Caltech (**Table S7**). For *Dalotia*, two rounds of MinION Nanopore vR9 sequencing libraries (Oxford Nanopore Technologies, UK) were prepared using genomic DNA extracted from approximately 25 male beetles using the Qiagen MagAttract HMW DNA Kit (Qiagen, Germany) and run on MinION flow cells at the Millard and Muriel Jacobs Genetics and Genomics Laboratory, Caltech.

### Bionano Optical Mapping

Optical maps were generated on the Bionano Genomics Saphyr system from ∼3 µg of ultra-high molecular weight genomic DNA extracted from 100 2^nd^ and 3^rd^ instar *Dalotia* larvae by HistoGenetics (Ossining, NY). The genomic DNA was fluorescently labeled with restriction enzyme DLE-1 (motif CTTAAG) with an average labeling density of 13 per 100 kbp. Total amount of labeled DNA was 755.67 Gbp. The raw Bionano data is available at CaltechDATA: https://doi.org/10.22002/1914a-m9460.

### SPRITE

For the Split-Pool Recognition of Interactions by Tag Extension (SPRITE) protocol, 92 male *Dalotia* were prepared as described in Quinodoz, et al. (*118*) with some modifications. Beetles were macerated with a glass dounce in 8 ml of 2 mM disuccinimidyl glutarate cross-linking solution at room temperature and rocked gently for 45 min. The cell suspension was pelleted by centrifugation for 8 min at 2500 xg at room temperature, rinsed in PBS and re-pelleted. A 3% paraformaldehyde solution in PBS was added and rocked gently at room temperature for exactly 10 min followed by the addition of 2.5 M glycine solution at room temperature for 5 min to quench the crosslinking reaction. Cells were pelleted by centrifugation at 4 °C for 4 min at 2500 x g. The pellet was washed in cold 1x PBS and 0.5% BSA two times, aliquoted, flash frozen in liquid nitrogen and stored at -80 °C. Cells pellets were thawed on ice and then lysed using buffers A, B and C in the SPRITE protocol (*119*) with buffer exchanges following centrifugation at 2500 xg for 8 min. The lysed cells were sonicated at 4 °C for 1 min (0.7s on, 3.3s off) with a chilled Branson needle tip sonicator. DNA fragmentation of lysate was performed with the addition of 3 µl of Turbo DNase (Thermo Fisher, CA) to 5 µl of lysate, 2 µl of 10X SPRITE DNase Buffer, and 5 µl of water at 37C for 17 min to obtain a fragment size distribution between 50 to 1000 bp. The cross-links were then reversed and the remainder of the protocol was followed as previously described. The distribution of cluster sizes and ligation efficiency was checked with a Illumina MiSeq run in house prior to shipping the twenty paired-end libraries for sequencing on the Illumina HiSeqX by Fulgent Genetics.

### Illumina genome assemblies

Read quality for each taxon was assessed using FastQC v0.11.8 (*120*). Illumina adapters, low-quality nucleotide bases (phred score below 15) from the 3’ and 5’ ends and reads shorter than 50 bp were removed using cutadapt v1.18 (*121*). From the filtered reads, *in silico* genome size estimates were calculated using kmergenie v.1.7048 (*122*) GenomeScope v1.0 (*123*), covEST v0.5.6 (*124*), and findGSE v0.1.0 (*125*). The latter three required a *k-mer* histogram computed by jellyfish v2.2.10 (*126*) with *k-mer* size of 21. The *in silico* estimates were compared to flow cytometry estimates for *Dalotia* (*n*= 13 female and n=14 male adult heads, and 3^rd^ and 1^st^ stage instars) performed by Dr. J. Spencer Johnston at Texas A&M University. Samples were run on a Beckman Coulter Cytoflex flow cytometer against both *Drosophila melanogaster* (1C = 175 Mbp) and *Drosophila virilis* (1C = 328 Mbp) standards as described in Johnston et al. 2019 (*127*). The ploidy level for each taxon was inferred using Smudgeplot (*128*) that calculates the coverage of heterozygous *k-mer* pairs from the short read sequences. A preliminary assembly was constructed from the filtered, adapter-trimmed reads using MEGAHIT v1.1.3 (*129*) with multiple *k-mer* sizes (--k-list= 21, 29, 39, 59, 79, 99, 119). Assembled contigs identified as bacterial contaminants with low GC content, high coverage and blast matches to the nr database (downloaded February 2019, e-value 1e-25) were removed using Blobtools v1.0 (*130*). For all the genome assemblies, except *Dalotia* described below, the filtered contigs were assembled into scaffolds with three iterations of the Redundans v0.14a (*131*) reference-based pipeline using the *Dalotia* hybrid assembly (v1) as a reference (--iters 3, --limit 0.5, --nogapclosing). Scaffolds smaller than 1 kb were removed and gaps were filled using GapCloser v1.12 (*132*).

### *Dalotia coriaria* genome assembly

The *Dalotia* genome was first assembled using a hybrid approach with short and long reads (**Fig S21**). Illumina reads were processed and assembled as described above until scaffolding. We removed 1,503 assembled bacterial contigs and 701 scaffolds smaller than 1000 bp prior to short-read scaffolding. Scaffolding was performed using SOAPdenovo2-fusion v2.04 (*132*) with a *k-mer* size of 75 optimized around the “best” k predicted by kmergenie. This was followed by long-read scaffolding with SSPACE-LongRead v1.1 (*133*) using uncorrected Nanopore reads (n= 4,150,648) and optimized parameters reported by Karlsson et al. (*134*). Separately, a long-read assembly was constructed with WTDBG2 v2.3 (*135*) using corrected Nanopore reads (n=848,141) from the *correct* step in canu v1.8 (*136*). We abandoned using canu beyond this step due to the runtime exceeding one month. The two genome assemblies (hybrid and long-read only) were merged using quickmerge v0.3 (*137*) (-hco 5.0 -c 1.5 -l 800000 -ml 10000) where the WTDBG2 assembly acted as the reference for whole genome alignment with nucmer (*138*). The merged hybrid assembly (*Dcor* v1, **Table S1**) was polished twice using racon v1.3.3 (*139*), gap-filled using LR_Gapcloser v1.1. (*140*) and finished with two additional rounds of short-read polishing using Pilon v1.23 (*141*). Allelic scaffold copies identified by Purge Haplotigs (*142*) based on both long-read (-l 15 -m 70 -h 100) and short-read (-l 8 -m 51 -h 140) coverage were removed resulting in the *Dcor* v1 assembly.

Consensus optical maps were generated *de novo* using Bionano Solve v3.7.1 and used to reorient and correct mis-assemblies of the *Dcor* v1 assembly using HybridScaffold v11657 (**Table S1**). Because only a third of the optical maps aligned to the *Dcor* v1 assembly, we aligned the optical maps to preliminary assemblies and raw reads with read length of 10kb or longer using RefAligner v12432 with default settings to calculate the proportion of contigs or reads not contained within the assembly. Assembly gaps were filled in this new assembly, *Dcor* v2, using LR_Gapcloser v1.1 with uncorrected Nanopore reads.

To get the assembly to the chromosome scale, the SPRITE fastq reads were processed by trimming the adapters using cutadapt v1.18 and identifying reads with five barcode tags using *BarcodeIdentification.jar* and *get_full_barcodes.py* scripts of SPRITE protocol. Complete reads were mapped to the *Dcor* v2 assembly with bowtie2 v2.3.4.1, filtered for mapping quality (-bq 20) and primary mapping (-F 256) using samtools v1.8 (*143*) and grouped into clusters using the *get_clusters.py* script from the SPRITE protocol. Clusters belonging to size classes 2 to 100 were first converted into the cool matrix format using *make_sprite_cooler.sh* script and then converted to the h5 format using hicexplorer v2.1.1. Matrix bin sizes were merged using *hicMergeMatrixBins* (-nb 30) and corrected using *hicCorrectMatrix* (--filterThreshold -2 2) to remove low and high coverage bins. The matrix was then used to orient and scaffold the *Dcor* v2 assembly using HiCAssembler v1.1.1 with coordinates of misassemblies identified using the *plotScaffoldInteractive* tool provided (--min_scaffold_length 200000 --bin_size 10000 --misassembly_zscore_threshold -1.0 --num_iterations 4). Pseudochromosomes 1 and 5 were manually split at low contact density regions and renamed using the bedtools “getfasta*”* tool (*144*).

To identify sequences that were not incorporated in the chromosome-resolved assembly, the preliminary assemblies from SSPACE-LongRead and WTDBG2 (both corrected and uncorrected versions, **Figure S1A**) were mapped back to the SPRITE assembly with minimap2 v2.15 full genome alignment setting (-ax asm5) (*145*). Unmapped scaffolds/contigs were extracted using samtools v1.8 utilities *view* and *fasta,* filtered using Purge Haplotigs with short-read coverage (-l 20 -m 51 -h 140) and then sequences shorter than 1000 bp were removed. The remaining contigs were combined with the SPRITE assembly for the final assembly version, *Dcor* v3. Genome completeness of *Dcor* v3 and the other genome assemblies used in this study was assessed using BUSCO v4.1.1 with the Arthropoda odb10 orthologous gene set (n=1013) curated from 90 species (*146*).

### Repeat identification and masking

To predict repeat content of the genome assemblies, we used a reference-based and a read-based approach. For the assembly-based predictions, we used methods described by Bruckner et al. (*147*). Species-specific libraries were constructed with RepeatModeler v 1.0.11 (*148*) and MITE tracker (*149*). Each library was filtered for genuine proteins based on significant blast homology (e-value 1e-5) to a local database of beetle proteins (*Agrilus planipennis*, *Anoplophora glabripennis*, *Aethina tumida*, *Dendroctonus ponderosae*, *Leptinotarsa decemlineata*, *Nicrophorus vespilloides*, *Onthophagus taurus*, and *Tribolium castaneum*; see **Table S3** for accessions). Blast reports were manually screened to remove non-repeat hits. Repeats without classification but blast homology to known TEs in the beetle protein database were retained whereas those with no blast homology were removed (*150*). The remaining repeat families were combined with the Arthropoda sequences in RepBase and clustered using vsearch v 2.7.1 (--iddef 1 --id 0.8 --strand both) (*151*). For each genome assembly, RepeatMasker v 4.07 (*152*) was used to soft mask repeats using the filtered repeat library. A summary of the masked repeat content was generated using the “buildSummary.pl” script, a utility of RepeatMasker. We also predicted the repeat content of each species using the adapter-trimmed reads with dnaPipeTE v1.3.1 (*153*), setting a genome coverage of 0.25 based on the predicted k-mer genome size estimates with two rounds of TRINITY assembly. The predicted repeats were filtered as described above by blast searches against the local database of beetle proteins, and reads counts adjusted to calculate the final repeat content.

We explored additional tools to annotate the repeat content of *Dalotia* given that the most abundant repeats lacked annotation from the dnaPipeTE results for the two *Dalotia* samples (WGS1 and WGS2). We used RepeatExplorer2 (*154*) that incorporates additional repeat databases and a satellite identification pipeline. We randomly subsampled two million paired-end reads from *Dalotia* WGS1 and *Dalotia* WGS2 using the “sample” tool in the program seqtk v1.3 (https://github.com/lh3/seqtk). The reads were uploaded to the RepeatExplorer2 Galaxy portal, and we followed the procedure one as described by Novák et al. 2020. Only 2% of the reads were used in the analysis due to RAM limitations of the Galaxy portal. Nevertheless, 60% of the reads for both samples were assigned to a 147 bp satellite (Dc-Sat1) that matched the abundant repeats of the dnaPipeTE results and was also present in the assembly-based method (“rnd-5_family-549”). To estimate the abundance of the Dc-Sat1 in the *Dcor* v3 assembly, we used bedtools v2.26.0 “intersect” given the genomic location of repeats predicted by RepeatMasker and bed files of the genomic coordinates of exons, introns and intergenic region boundaries. To see if the Dc-Sat1 was shared among the beetles in this study, five million reads were subsampled from each species and screened for the consensus sequence of Dc-Sat1 using RepeatProfiler v1.1 (*155*) with default settings. Lastly, we estimated the Kimura’s distance, or nucleotide sequence divergence, of the Dc-Sat1 with RepeatMasker on a subset of five million reads followed by RepeatMasker utility scripts “buildSummary.pl” and “calcDivergenceFromAlign.pl”. Long minION reads with abundant copies of Dc-Sat1 as determined by TideHunter v1.2.2 using default settings (*156*) were visualized using FlexiDot v1.06 with a word size of 147 (*157*). The secondary structure of the Dc-Sat1 satellite was predicted using VectorBuilder (https://en.vectorbuilder.com/tool/dna-secondary-structure.html).

### Dalotia gene predictions and annotation

A combination of *ab inito* (GeneMark-ES v4.33(*158*) and reference-based (BRAKER v2.1.2 (*159*), PASA v 2.3.3 (*160*), exonerate (*161*) and GeMoMA v1.6.1 (*162*)) tools were used for gene prediction in the *Dalotia* assembly versions as previously described (*147*). For BRAKER and PASA, diverse transcriptomic data sets (larvae, pupae, male and female antenna, male and female whole body, female brain, and abdominal segments 6 and 7) were mapped to the *Dalotia* genome *Dcor* v3 using STAR v2.6.1 (*163*). With the resulting alignment file, a genome-guided transcriptome assembly was constructed with TRINITY v2.5.1 (*164*) as described below. The transcriptome assembly constructed from all tissue types and life-stages was then used for gene prediction with PASA run with the Transdecoder option (https://github.com/TransDecoder/TransDecoder), GMAP (*165*) and blat (*166*) aligners, and a maximum intron length of 300 kb. To identify homologs of insect genes, we aligned 3,483,422 insect genes from the UniProt database (downloaded March 2019) to the *Dalotia* genome using exonerate v2.4.0, keeping alignment predictions with at least 80% percent coverage.

For the *Dcor* v1 assembly, gene predictions were combined with EVidenceModeler (*160*) with the following weights: PASA=10, BRAKER_HiQ=4, BRAKER=1, GeneMark=1, and exonerate=1. BRAKER_HiQ predictions were given higher weight because they had >90% coverage of the exon boundaries. Gene predictions from *Dcor* v1 were lifted over to subsequent versions using Liftoff v1.6.1 (*167*) with default settings and the polish option. In place of exonerate in later assembly versions, we used the homology-based prediction tool GeMoMa v1.6.1 with gene models from the beetle phylogenetically closest to *Dalotia* with a previously annotated genome, *Nicrophorus* (Staphylinidae: Silphinae; NCBI: GCF_001412225.1), as we as from the beetle with the highest quality, annotated coleopteran genome, *Tribolium* (Tenebrionidae; NCBI: GCF_000002335.3). We combined all predictions with EVidenceModeler with the following weights: GeMoMa=4, PASA=4, Liftoff= 4, BRAKER_HiQ=4, BRAKER=1 and GeneMark=1. The predicted genes were searched against the NCBI nr (February 2019), UniProt (February 2019), PFAM (v 32, August 2018), merops (v 12, October 2017) and CAZy (v 7, August 2018) databases. The hmm-based results of PFAM and CAZy were filtered using cath-tools v 0.16.2 (https://cath-tools.readthedocs.io/en/; e-value 1e-5) and the blast-based searches were filtered by the top hit (e-value 1e-5 threshold). Predicted genes were also assigned to orthologous groups using eggNOG emapper v2.1.5 (*168*) against the eggNOG 5.0 database. Gene annotation was assigned by the UniProt hit if the e-value < 1e-10 followed by NCBI annotation if the e-value < 1e-10, and then eggNOG annotation if the e-value < 1e-10. If no homology was recovered, then the gene was annotated as “hypothetical protein”. The final assembly and associated annotation files can be downloaded at CaltechDATA: https://doi.org/10.22002/62xxb-mak64.

### Gene predictions of other genome assemblies

A similar strategy to gene prediction was used for the remaining genome assemblies presented in this study. When transcriptomic data was available (*Holobus* sp., *Drusilla canaliculata*, *Lissagria laeviuscula*, *Aleochara* sp. 3, and *Liometoxenus newtonarum*), both *ab inito* and reference-based tools were used as described above with slight modifications. In addition to *Nicrophorus* and *Tribolium* gene models, gene models from *Dcor* v2 assembly were used for the homology-based predictions with GeMoMa. The respective genome-guided transcriptome assemblies for each species based on available whole body RNAseq read sets were used as the input of PASA and BRAKER and run as described above for *Dalotia*. EvidenceModeler weights were assigned as follows: PASA= 10, BRAKER_HiQ=4, BRAKER=1, GeMoMa=1, and GeneMark=1. For species where no transcriptomic data was collected, we only used *ab inito* and homology-based predictions. We used an additional *ab inito* tool augustus v3.23 (*169*) that was run with three different configuration files: honeybee, tribolium2012, and species-specific file based on a random set of 200 genes from the BUSCO training set using the etraining tool. To combine the *ab inito* predictions with GeMoMa predictions, EVidenceModeler weights were GeMoMa=5, species-specific=1, honeybee=1, tribolium2012=1, and GeneMark=1. All Illumina-only genome assemblies are available at CaltechDATA: https://doi.org/10.22002/k8sfv-dw648. Predicted genes of *Aleochara* sp., *Holobus* sp. and *L. newtonarum* were assigned annotation through either orthology to *Dalotia* genes from the OrthoFinder2 results or from eggNOG orthology searches when no *Dalotia* orthologue was found.

### Phylogenomic tree construction and dating

For the phylogenomic analysis, we included the genome assemblies of 26 Staphylinidae species from this study and nine published genome assemblies of beetle species spanning the suborder Polyphaga (**Table S3**). In the case of the published genome assemblies, we removed predicted isoforms with cdHIT v4.8.1 (*170*) if the pairwise protein sequence identities were at least 98% identical (-c 0.98) for at least 30% of the alignment (-aL 0.3 -aS 0.3). Protein-coding sequences for all species were clustered into orthogroups, a group of orthologous genes, with OrthoFinder v2.5.2 (-M msa -S diamond_ultra_sens -A mafft -T fasttree) (*171*). The 9,971 mafft sequence alignments of orthogroups that had at least 18 taxa present were then trimmed using the gappyout method of trimAl v1.4.1 (*172*). An approximate maximum likelihood gene tree was constructed for each trimmed alignment with FastTree2 v2.1.11 (-slow –gamma) (*173*). To reduce the alignments to a strict set of orthologs, we used PhyloPyPruner v1.2.4. (https://gitlab.com/fethalen/phylopypruner) with the following parameters: --min-len 100 --trim-lb 3 --min-support 0.75 --prune MI --min-taxa 28 --mask pdist --trim-divergent 0.75 --min-pdist 0.01 --min-gene-occupancy 0.1 --subclades subclade.txt --root midpoint --outgroup Apla PPYR. The resulting concatenated supermatrix consisted of 1,300,484 amino acid sites with 3,060 gene partitions. To improve the phylogenetic signal, the information content of each partition was calculated using MARE_v0.1.2-rc with default settings, except to ensure all taxa were retained (*174*). The optimized supermatrix from MARE contained 1,520 gene partitions (577,200 aligned amino acid sites).

With the reduced and optimized gene partitions, we constructed the species tree using both maximum likelihood and quartet-based coalescent methods. To find the best substitution model, we ran ModelFinder (*175*) with a subset of protein models (LG, WAG, JTT, Dayhoff, Q.insect) on the gene partitions and examined the top 10% of the partition merging schemes (-rcluster 10) (*176*). Using the best-scoring partitioning scheme, a maximum likelihood species tree was estimated from the concatenated supermatrix using IQ-TREE v2.2.0-beta (*177*) with a 1,000 ultrafast bootstrap replicates (*178*). For the same set of genes, a coalescent species tree analysis was carried out in ASTRAL v5.6.3 (*179*) using gene trees estimated from the pruned alignments in IQ-TREE following model selection by ModelFinder. Topological support is presented as the quartet support, or gene tree conflict around a given node.

To date the species tree, ten conservative fossil calibration points were selected from a literature survey (**Table S4**). This set of fossils contained eight calibration points previously reported for the family Staphylindae (Maruyama and Parker 2017). The other two calibration points were selected from recent phylogenomic studies on Coleoptera (Zhang et al. 2018, McKenna et al. 2019 and Cai et al. 2022). These included bounded constraint on the root of the tree, the Crown Polyphaga (237 to 293 Ma), and lower bound estimate on Crown Chrysomeloidea (122.5 Ma). Divergence time analysis was performed with MCMCtree and CODEML implemented in the PAML v4.9 package (*180*) on the concatenated supermatrix and maximum likelihood species tree. As part of the approximate likelihood calibration method, we generated a Hessian matrix in CODEML using empirically estimated base frequencies on the protein supermatrix from the LG substitution matrix (lg.dat) with four rate categories. We obtained 200,000 trees with a sampling frequency every 100 iterations after discarding 20,000 trees as burn-in. Default parameters were set as described in McKenna et al. 2019. For all calibration points except the root age, we applied a soft minimum age using a truncated Cauchy distribution with an offset of 0.1, scale parameter of 1 and left tail probability of 2.5%. At the root, we provided a soft joint bound with an error probability of 0.1 on the minimum and maximum age. Convergence of two independent MCMC runs was checked in Tracer v1.7.2 (*181*). The final species tree was plotted using the R package MCMCtreeR (*182*).

### Phylogenetic analyses of select gene families

For select orthogroups of interest, we manually refined gene predictions where necessary and constructed gene trees. We manually screened sequences for the presence of start and stop codons and compared the length of each sequence against the length distribution of all sequences within a given orthogroup. If sequences were flagged as partial, we extracted the corresponding scaffold from the genome and attempted to extend the scaffold manually using the unfiltered megahits assembly of that species. The extended scaffolds were then re-processed through the Augustus webserver using either the *Apis mellifera* or *Tribolium castaneum* configuration files to re-predict coding sequence. We added *Drosophila melanogaster* orthologs to each orthogroup using phylogenetic-informed orthology searches with shoot.bio (*183*) as well as literature searches. We aligned the protein sequences with mafft v7.505 (*184*). The protein alignment was then trimmed with trimAl v1.4.1 using the *gappyout* method. A maximum likelihood tree was constructed with both the trimmed and untrimmed protein alignments using IQ-TREE v2.2.0-beta with a 1,000 ultrafast bootstrap replicates. The best protein model was selected by ModelFinder with a subset of substitution models (LG, WAG, JTT, Dayhoff, and Q.insect). Classification of FARs and esterases followed the assigned groups designated by Tupec et al. (*185*) and Oakeshott et al. (*186*) respectively. Curated protein and nucleotide sequences used in the phylogenetic analyses and IQ-TREE tree files can be found at CaltechData: https://doi.org/10.22002/cgsw0-9kk67.

### Selection tests and inactivating mutations

We performed positive selection tests using the adaptive branch-site random effects likelihood method (aBSREL) in the HyPhy package v2.5.38 (*187, 188*) and the branch-site models implemented by CODEML in the ete3 v3.1.2 toolkit (*180, 189*). Both tools used the codon alignment and gene tree as input. Protein alignments were converted into codon alignments with tranalign v6.6.0.0, a tool within the EMBOSS suite (*180*). For aBSREL, we tested branches using both an exploratory approach across the whole tree and hypothesis approach on select branches of interest (foreground) against the background. A likelihood ratio test was performed on the fit of the full model on each branch against the null model, where no positive selection rate class is allowed on that branch. For CODEML, we tested the branch-site model on select branches and the model fit was compared against the null model with a likelihood ratio test. For branches under selection, the Bayes-Empirical Bayes method identified codons with signatures of positive selection that had a posterior probability threshold ≥ 0.95.

Inactivating mutations were detected using an orthology-based, reference genome alignment method Tool to infer Orthologs from Genome Alignments (TOGA (*190*) for the three ecitocharine taxa against the *Dcor* v3 assembly. To make alignment chain files, each taxon was aligned to the *Dalotia* assembly twice using the utility script “make_chains.py” (https://github.com/hillerlab/make_lastz_chains) with default settings (K = 2400, L = 3000, H = 2000, Y = 9400, default lastz scoring matrix) and University of California Santa Cruz genome browser settings for insect alignments (K = 2200, L = 4000, H = 2000, Y = 3400, HoxD55.q lastz scoring matrix). The chain files were then used as input for TOGA with the “–fragmented-genome” parameter to infer orthologous genes from multiple aligned contigs. To account for sequencing errors and/or sequence divergence, the predicted gene-inactivating mutations (frameshift insertion/deletions, premature stop codons, splice site mutations and deletions of exons or entire genes) from the core biosynthetic differentially expressed orthologs of the solvent and BQ cells (n=554) were manually inspected with independent gene predictions for each respective taxon and predicted mutations from snpEff v5.0e (*191*) using a variant call file (VCF) produced by aligning the short reads of each ecitocharine taxon to the *Dalotia* genome assembly with bwa v0.1.17 (*192*), following the GATK best practices pipeline (*193*), and filtering SNPs (’MQ > 40 & INFO/DP < 1200 & QD > 2.0 & FS < 60.0 & MQRankSum > -12.5 & ReadPosRankSum > -8.0 & SOR < 3.0) and INDELS (MQ > 40 & INFO/DP < 1200 & QD > 2.0 & FS < 200.0 & ReadPosRankSum > -20.0 & SOR < 10.0) with bcftools v1.8. (*143*). Given the fragmentation of our assemblies from the ecitocharine taxa (**Table S3**), we excluded predicted “loss” genes if the evidence was based solely on missing and/or deleted exons. Mutations were visualized using the “plot_mutations.py” utility script. The results of TOGA and annotated VCF files from snpEff are available on CaltechData: https://data.caltech.edu/records/6xjn1-e3085.

### Gene synteny

We compared the gene content and identified the sex chromosomes of the *Dalotia* genome assembly against the chromosome-scale genome assemblies of the outgroup beetles *T. castaneum* (NCBI: GCF_000002335.3) and *P. pyralis* (http://www.fireflybase.org/), and two rove beetles *Ocypus olens* (NCBI: GCA_910593695.1) and *Philonthus cognatus* (NCBI: GCA_932526585.1). Gene synteny was assessed using the “promer” and “show-coords” programs within the MUMmer package v 3.23 with an alignment length of at least 100 amino acids (-L 100) and percent identity of 50% (-I 50) between the reference and target genomes. To identify regions of gene synteny between all pairwise genome comparisons, the all-vs-all blast results from OrthoFinder were used as input for DAGchainer (-M 50 -D 5 -g 1 -A 3 -E 10) (*194*).

### Gland volatile quantification

Beetles were individually submersed in 70 μl hexane (NN), after 10 min the solvent was separated from the insect, transferred into a new vial and frozen at -80°C for further analysis. A GCMS-QP2020 gas chromatography/mass-spectrometry system (Shimadzu, Kyōto, Japan) equipped with a ZB-5MS fused silica capillary column (30 m x 0.25 mm ID, df= 0.25 μm) from Phenomenex (Torrance, CA) was used to profile the gland contents: crude sample aliquots (2 μl) were injected into split/splitless-injector which operated in splitless-mode at a temperature of 310°C. Helium was used as the carrier-gas with a constant flow rate of 2.13 ml/min. The chromatographic conditions were as follows: column temperature was set to 40°C with a 1-minute hold after which the temperature was initially increased 30°C/min to 250°C and further increased 50°C/min to a final temperature of 320°C and held for 5 minutes. Electron impact ionization spectra were recorded at 70 eV ion source voltage, with a scan rate of 0.2 scans/sec from *m/z* 40 to 450. The ion source of the mass spectrometer and the transfer line were kept at 230°C and 320°C, respectively. Compounds were previously identified and in addition authentic standards were used to construct four-point calibration curves for external standardization and quantification of benzoquinones, esters and alkanes.

### Ancestral state reconstruction

We used ancestral state reconstruction to estimate chemical class evolution along the species tree. Each chemical class was treated as a binary, discrete character of either present (1) or absent (0) in a given extant lineage. Extant taxa for which no chemical data has been collected were assigned a value of “-9”. We first applied a maximum likelihood method using an equal rates model with the *ace* command in R package ape v5.6-2 (*195*). Second, we used the re-rooting method of Yang et al. (*196*) to estimate marginal states for species with no chemical data implemented in phytools v1.0-3 (*197*). Probabilities of the state being absent were assigned a value of 0.5 in *Aleochara* sp1, *Falagria* and *Earota* and 0.9 in the ecitocharine clade.

### Biochemical tracer experiment in Liometoxenus

Wild caught *Liometoxenus* individuals were housed in 10 cm plastic containers with moistened tissue paper for several days with various food sources (sugar water, dead ants and frozen fruit flies) prior to experimentation. Ten beetles were chemically disarmed on CO_2_ gas as previously described for *Dalotia* (*32*) and split into two containers, one with the same food sources and the other where the stable isotope precursor ^13^C_6_-tyrosine (>99% enrichment, Sigma-Aldrich, MO) was added to each food source. The isotope-labeled and control food was refreshed every three days. Beetles were sacrificed over the course of two weeks for hexane extractions either because their health declined, or the end of the experiment was reached. Hexane extracts were analyzed with a GC-MS as described above. Electron ionization mass spectra of characteristic fragment ions were monitored in single ion mode (SIM) and at 70 eV.

### Double-stranded RNA synthesis and knockdown

Double-stranded RNA constructs were prepared as previously described (*32, 114*). Our target sequences were cloned into a pCR2.1-TOPO vector (Thermo Fisher, CA) using primers with T7 linkers as follows: *very long-chain-fatty-acid-CoA ligase bubblegum* (*bgm*) F: (5’-TAATACGACTCACTATAGGGCGATGCTGAAGGTTGGCTAC-3’) and R: (5’-TAATACGACTCACTATAGGGCAATTTCAATGTGGGCCCCA-3’), *copper-transporting ATPase 1* (*ATP7*) F: (5’-TAATACGACTCACTATAGGTGACAACGCAGGATATCCCTCCGG-3’) and R: (5’-TAATACGACTCACTATAGGGCTTCTGGTTTCACAGGATCCGCC-3’), and *β-glucosidase* (*BGLU*) F: (5’-TAATACGACTCACTATAGGGCGTGCGCGTGTTGATTACGTC-3’) and R: (5’-TAATACGACTCACTATAGGTGCAGTAACGCGAACGCCATCA-3’). After synthesis, the dsRNA was cleaned using the MEGAclear Transcription Clean-Up kit (Thermo Fisher, CA) and quantified on the NanoDrop (Thermo Fisher, CA). Target and control, a green fluorescent protein, constructs were diluted with DEPC-treated 1x PBS and 1 µl of blue food dye to a final concentration of 2 mg/ml. The constructs were microinjected into third instar larvae from the laboratory *Dalotia* population according to Parker et al. (*114*). Following injection, larvae were reared in individual 5 cm Petri dishes on filter paper until eclosion. After eclosion, adult beetles were moved into new Petri dishes and fed frozen fruit flies for ten days, at which point they were used for chemical analysis. Statistical difference of glandular secretions of specific compounds between RNAi-treated and GFP-treated was tested with Wilcoxon signed rank test with a Bonferroni multiple test correction for the various compounds per beetle.

### Drosophila toxicity bioassay

We tested the toxicity of the major compounds of the *Holobus* gland secretion on survival of *Drosophila melanogaster* larvae as previously described (*32, 67*). The major compounds were prepared to mimic natural ratios of the gland secretion: 5% of tridecane, 15% of 2,5-dimethoxybenzaldehyde, and 80% of ethyl linoleate (all Sigma Aldrich, MO) (**Fig. 6A**). Each compound was tested independently along with the mixture of all three compounds in the *Holobus* glandular secretion. We also tested the addition of 2-methyl-1,4-benzoquinone without a solvent (powder form) and with the *Holobus* secretion mixture (28 mg). A mixture of the *Dalotia* gland secretion compounds (*32*) and 1x PBS were used as the positive and negative controls, respectively. Over two experimental trials, wandering third instar *Drosophila* larvae were submerged in the 1 ml of the various mixtures for ∼1 s or dipped in solid BQ powder (n=25 larvae per mixture) and moved to three replicate culture tubes. Survival was scored after 1 hr and after eclosion. At 1 hr, dead larvae were distinguishable by a change in coloration to black or dark down, or loss of tissue integrity. Differences in survival were tested using an ANOVA with a Tukey *post hoc* test correction in the statistical package R v4.2.1.

### Chromosome squashes

The chromosome preparation protocol was modified from Rożek et al. (*198*). Testes of immobilized *Dalotia* (*n*=10) were dissected in 1x PBS under a stereomicroscope. Testes were transferred to a hypotonic KaryoMAX Colcemid Solution (Gibco, NY) at a final concentration of 0.5 µg/ml in 1x PBS for 1 hr at room temperature with gentle rocking. The solution was discarded after 2 min centrifugation at 500 xg and replaced with 2 ml of 0.075M KCl for 20 min. Following another round of centrifugation, the testes were transferred to freshly prepared Fix I solution (3:1 absolute 96% ethanol:glacial acetic acid) and left to sit for 30 min at room temperature. The Fix I solution was replaced after 30 mins with fresh Fix I and stored at 4 °C for up to two years. The remaining fixative solutions (Fix II – 1:1 absolute 96% ethanol:glacial acetic acid and Fix IV – 7:2:1 glacial acetic acid: absolute 96% ethanol:distilled water) were prepared fresh and brought to 32 °C when preparing for the squashes. The testes were transferred from Fix I to Fix II and then Fix II to Fix IV, with 30 min incubation intervals in each solution at room temperature. The testes were stored in Fix IV at 4 °C overnight for 10-12 hr. Fixed testes tissue was then transferred to a clean microscope slide resting on blotting paper. The tissue was macerated quickly using dissecting needles in a few drops of 70% acetic acid. The tissue was squashed between two microscope slides as described in (*198*) and frozen on dry ice. The final preps were stained with nuclear stain Hoechst 33342 (1:2000), mounted in 25 µl of VectaShield Mounting Media (Vector Laboratories, CA) and imaged using the 100x objective on the Zeiss LSM 880 Confocal Laser Scanning Microscope (Zeiss, Germany).

### Electron Microscopy and Dual-Axis Tomography

For sample preparation, beetle abdomens were dissected in a fixative comprising 3% glutaraldehyde, 1% paraformaldehyde, 5% sucrose in 0.1 M sodium cacodylate trihydrate. Dissected tissue was then placed in fresh fixative at 4°C. Pre-fixed segments were rinsed with fresh cacodylate buffer and placed individually into brass planchettes (Type A; Ted Pella, Inc., CA) prefilled with 10% Ficoll in cacodylate buffer. Samples were covered with the flat side of a Type-B brass planchette and then ultrarapidly frozen with a HPM-010 high-pressure freezing machine (Bal-Tec/ABRA, Switzerland). The vitrified samples were transferred under liquid nitrogen to cryotubes (Nunc) containing a frozen solution of 2.5% osmium tetroxide, 0.05% uranyl acetate in acetone. Tubes were loaded into an AFS-2 freeze-substitution machine (Leica Microsystems, Vienna) and processed at -90°C for 72 h, warmed over 12 h to -20°C, held at that temperature for 6 h, then warmed to 4°C for 2 h. The fixative was removed, and the samples rinsed 4x with cold acetone, following which they were infiltrated with Epon-Araldite resin (Electron Microscopy Sciences, PA) over 48 h. Samples were flat-embedded between two Teflon-coated glass microscope slides and the resin was polymerized at 60°C for 48 h.

Flat-embedded beetle segments were observed by phase-contrast LM to determine sample quality and specifically locate suitable tergal gland components. These regions were extracted with a microsurgical scalpel, oriented for en face (dorsal-to-ventral) sectioning and glued to the tips of plastic microtomy stubs. Semi-thick (170 nm) serial sections were cut with a UC6 ultramicrotome (Leica Microsystems, Vienna) using a diamond knife (Diatome, Ltd. Switzerland). Sections were placed on Formvar-coated copper-rhodium slot grids (Electron Microscopy Sciences, PA) and stained with 3% uranyl acetate and lead citrate. Gold beads (10 nm) were placed on both surfaces of the grid to serve as fiducial markers for subsequent tomographic image alignment. Grids were placed in a dual-axis tomography holder (Model 2040, E.A. Fischione Instruments, PA) and imaged with a Tecnai T12 transmission electron microscope (120 KeV) equipped with a 2k x 2k CCD camera (XP1000; Gatan, Inc. Pleasanton CA). Tomographic tilt-series and large-area montaged overviews were acquired automatically using the SerialEM software package (*199*). For tomography, samples were tilted +/-62° and images collected at 1° intervals. The grid was then rotated 90° and a similar series taken about the orthogonal axis. Tomographic data was calculated, analyzed and modeled using the IMOD software package (*200–202*) on iMac Pro and Mac Studio M1 computers (Apple, Inc., Cupertino, CA).

### RNA extraction and transcriptome assemblies

Specimens used for transcriptome sequencing (*Aleochara sp.* 3 male body (n=1), *Dalotia* male antenna (n= approx. 100), female antenna (n= approx. 100), female brain (n=1), larvae (n= approx. 100), pupae (n= approx. 20), male body (n=1), female body (n=1), *Holobus* male body (n=5), and *Liometoxenus* male head (n=1) and body(n=1)) were either extracted live or from flash-frozen material stored at -80 °C. Total RNA was extracted from the different species, life stages and tissue types using either the ZYMO *Quick*-RNA Tissue/Insect extraction kit (ZYMO Research, CA) or a combination of Trizol (Life Technologies, CA) and Qiagen RNeasy Mini kit (Qiagen, Germany) extraction protocol as previously described (*203*) (see **Table S7**). RNA integrity and quantity was assessed with the Nanodrop (Thermo Fisher, CA) and Bioanalyzer High Sensitivity RNA Analysis kit (Agilent Technologies, CA). Paired-end, 150bp sequencing libraries were prepared using the Illumina TruSeq RNA library kit by various companies listed in **Table S7** and sequenced on Illumina HiSeq X platform (Illumina, CA).

Transcriptomes used in gene predictions described above were either assembled *de novo* (*Liometoxenus*) or from genome-guided RNAseq read alignments (*Dalotia*, *Holobus* and *Aleochara* sp. 3) with Trinity v2.5.1 (*204*) using the diverse data sets available for each species (**Table S7**). For the genome-guided assemblies, adapter-trimmed RNAseq reads were aligned to each respective reference genome using STAR v2.6.1 (*163*) and assembled with the maximum intron length of 10000bp and jaccard clip option in Trinity. Previously published *de novo* assembled transcriptomes of *Drusilla* and *Lissagria,* both construced from male and female whole body RNAseq reads, were also used in gene predictions (*32*).

### SMART-seq transcriptome sequencing

Microdissection of the specific gland cell types from *Aleochara* and *Liometoxenous* was performed as previously described (*32*). Due to the size of *Holobus* (**Fig. S12E**), the entire tergite 6 (control) and tergite 7 (gland segment) were dissected in ice-cold DEPC-treated PBS, flash frozen and stored at -80°C until processed. Library preparation was done from either frozen cells or Trizol extracted total RNA (3 out of 4 *Aleochara* control samples) using the NEBNext Single Cell/Low Input RNA Library Prep Kit for Illumina together with NEBNext Multiplex Oligos (New England Biolab) according to the manufacturer’s protocol. PCR cycles during the cDNA amplification step varied depending on the sample type and species. For example, in *Aleochara*, cycles ranged from 9 PCR cycles for total RNA input, 14 PCR cycles for solvent cells up to 20 PCR cycles for BQ cells. All *Holobus* preps were held for 14 PCR cycles and all *Liometoxenous* preps were held for 20 PCR cycles. Final library amplification ranged from 8-12 PCR cycles depending on the intermediate concentrations of the library during the procedure. The quality and concentration of the resulting libraries were assessed using the Qubit High Sensitivity dsDNA kit (Thermo Scientific) and Agilent Bioanalyzer High Sensitivity DNA assay. The 50bp libraries were sequenced on Illumina HiSeq2500 or NextSeq 2000 with about 20-30 million reads per library by Millard and Muriel Jacobs Genetics and Genomics Laboratory at Caltech.

### Differential expression analysis

SMART-Seq reads were aligned to each respective species genome assembly with STAR v2.6.1 (*163*) and read counts extracted with featureCounts v2.0.0 (*205*) only considering primary alignments (--primary) that mapped to the same chromosome and strand (-C) with a minimum mapping quality of 10 (-Q 10). Genes with fewer than 10 read counts for the minimum group size of a given species and cell type were removed (*Dalotia* n=10, *Aleochara* n=4, *Holobus* n=4, and *Liometoxenus* n=5). Differential gene expression was tested for each species using DESeq2 v1.30.1 (*206*) with the design *tissue type* (BQ cell, Solvent cell, or control) + *batch* for cell-specific data sets of *Dalotia*, *Aleochara* and *Liometoxenus* or *segment type* (gland or non-gland) *+ batch* for bulk abdominal segment comparisons of *Holobus* and *Dalotia*. Sequencing batch was added for all species except *Aleochara*, which was processed in one sequencing run. Bulk RNAseq reads from *Dalotia* gland and non-gland segments (*32*) were processed as above with technical replicates collapsed using “collapseReplicates” function in DESeq2. DEGs were identified in each species for each pairwise comparison of cell type or segment type using a Wald test with adjusted p-value ≤ 0.05). DEGs that displayed cell type enriched expression were those with 2-fold higher log_2_ expression in one cell type relative to the other gland cell type or control.

To compare expression among species, variance stabilized count matrices of all genes for each species were joined by the OrthoFinder assigned orthologous groups. In cases where multiple orthologs were assigned to the same orthogroup, genes were sorted by their adjusted p-values from the gland cell type against control tests, with the lowest value selected to represent the orthogroup. From this reduced matrix, the median expression value of a 1000 conserved orthologs (lowest Coefficient of Variation (CV; standard deviation/mean expression)) among the species was used as the scaling factor to further normalize the read counts to account for species differences (*207*). A principal component analysis was performed on the transformed data using prcomp function in the R package Stats v3.6.0, first with all orthologues and second with only differentially expressed orthologs in one pairwise test, or DEOs. An UpSet plot of the ortholog expression by cell type and species was inspired by customized_upset_plots (https://github.com/cxli233/customized_upset_plots). To summarize gene functions, GO and KEGG enrichment test on core BQ cell and solvent cell DEOs were then performed with clusterProfiler v3.18.1 (*208*) using a q-value cutoff of ≤0.05 and the simplify function to reduce similarity in GO terms. A custom gene ontology (GO) database was made for *Dalotia* using GO terms assigned from the eggNOG database and Uniprot blast matches with AnnotationForge v1.38.0 (*209*).

To explore the conservation of abdomninal gene expression programs (GEPs) identified in *Dalotia* from a prior study (*32*) with other species, *Dalotia* transcripts with high z-score rank to the cuticle cells and ventral fat body and oenocytes GEPs were mapped to the *Dalotia* gene models using GMAP v 2017-11-15 (*165*). Spearman correlation of GEP expression between *Dalotia* genes and their corresponding *Aleochara* orthologs was performed using cor.test in R. To get qualitative differences between tissue types and life-stages of *Dalotia*, all transcriptome data sets were mapped to the Dcor v3 assembly using STAR v2.6.1 (*163*) and gene counts extracted using featureCounts v2.0.0. Heat maps were generated from normalized variance stabilized counts from DESeq2 and the R package pheatmap. Sex-biased expression was calculated as the difference in library normalized log_2_ counts using the *normTransform* function in DESeq2 for the male and female whole-body transcriptomes. Differences were categorized as 2-, 5- and 10-fold higher in one sex over per gene and then tabulated by chromosome. Statistical differences in the proportion of biased genes were found using a Pearson’s Chi-square test with multiple test correction with R package chisq.posthoc.test v0.1.2

### In Situ Hybridization Chain Reaction

DNA probe sets were either purchased from Molecular Technologies (Pasadena, CA; https://www.moleculartechnologies.org/) or generated using the “insitu_probe_generator” tool (https://github.com/rwnull/insitu_probe_generator) and the pool of oligos was purchased from Integrated DNA Technologies (Coralville, IA) (**Table S8**). DNA HCR amplifier, HCR hairpins as well as hybridization, wash and amplification buffers were purchased from Molecular Technologies. The abdominal sections of adult *Oligota* sp. and *Aleochara* sp. 3 beetles were fixed as previously described for *Dalotia* (*32*). The amplification and detection stages followed published protocols (*210*). Probes were initiated with B1-Alexa546, B3-Alexa647 or B4-Alexa488 amplifiers. After amplification and before the final wash steps, Hoechst 33342 (1:2000) to mark nuclei, and Alexa 488-or Alexa-647-Wheat Germ Agglutinin Conjugate (WGA; 1:200) to label cell membranes were added. Tissue samples were imaged as whole mounts of dorsal abdomens in ProLong Gold Antifade Mountant (ThermoFisher), using a Zeiss LSM 880 with Airyscan fast.

### Dmd enzymatic activity

Purified protein of Dmd from *Dalotia, Holobus* and *Ecitophya* was prepared as described by Bruckner et al. (*32*). Enzymatic activity of each protein was tested against a standard substrate, ABTS, and hydroquinone (HQ) substrates (1,4-hydroquinone, 2-methyl-1,4-hydroquinone and 2-methoxy-3-methy-1,4-hydroquinone). The reaction mixture was prepared as 5 mM MES, 0.3 M CuSO_4_, and either 2 mM of ABTS or 2mM HQ, with 0.5 mM of the test protein. Reactions were UV recorded for 1 min and directly quenched with 0.05M EDTA before being flash frozen and stored at -80 °C until further analysis. A total of three replicates were tested for each substrate-enzyme combination and the no-enzyme control. We used a liquid-liquid extraction protocol described previously for chemical profiling the end reactions with the GC/MS (Brucker et al. 2021) and sample aliquots (1 µl) were quantified on GCMSQP2020 gas chromatography/mass spectrometry system equipped with a ZB-5MS fused silica capillary column (30 m x 0.25 mm ID, df = 0.25 mm) from Phenomenex (Torrance, CA, USA). Helium was used as the carrier-gas with a constant flow rate of 2.13 ml/min. The chromatographic conditions were as follows: column temperature at the start was 40 C with a 1-min hold after which the temperature was initially increased 50 C/min to 250 C and further increased 60 C/min to a final temperature of 300 C and held for 2 min. Electron impact ionization spectra were recorded at 70 eV ion source voltage, with a scan rate of 0.2 scans/sec from m/z 40 to 450. The ion source of the mass spectrometer and the transfer line were kept at 230 C and 320 C, respectively. We used synthetic 1,4-BQ, 2-methyl-1,4-BQ and 2-methoxy-3-methyl-1,4-BQ to quantify the amounts of benzoquinones in nanograms.

---

**Supplemental Figure 1.**
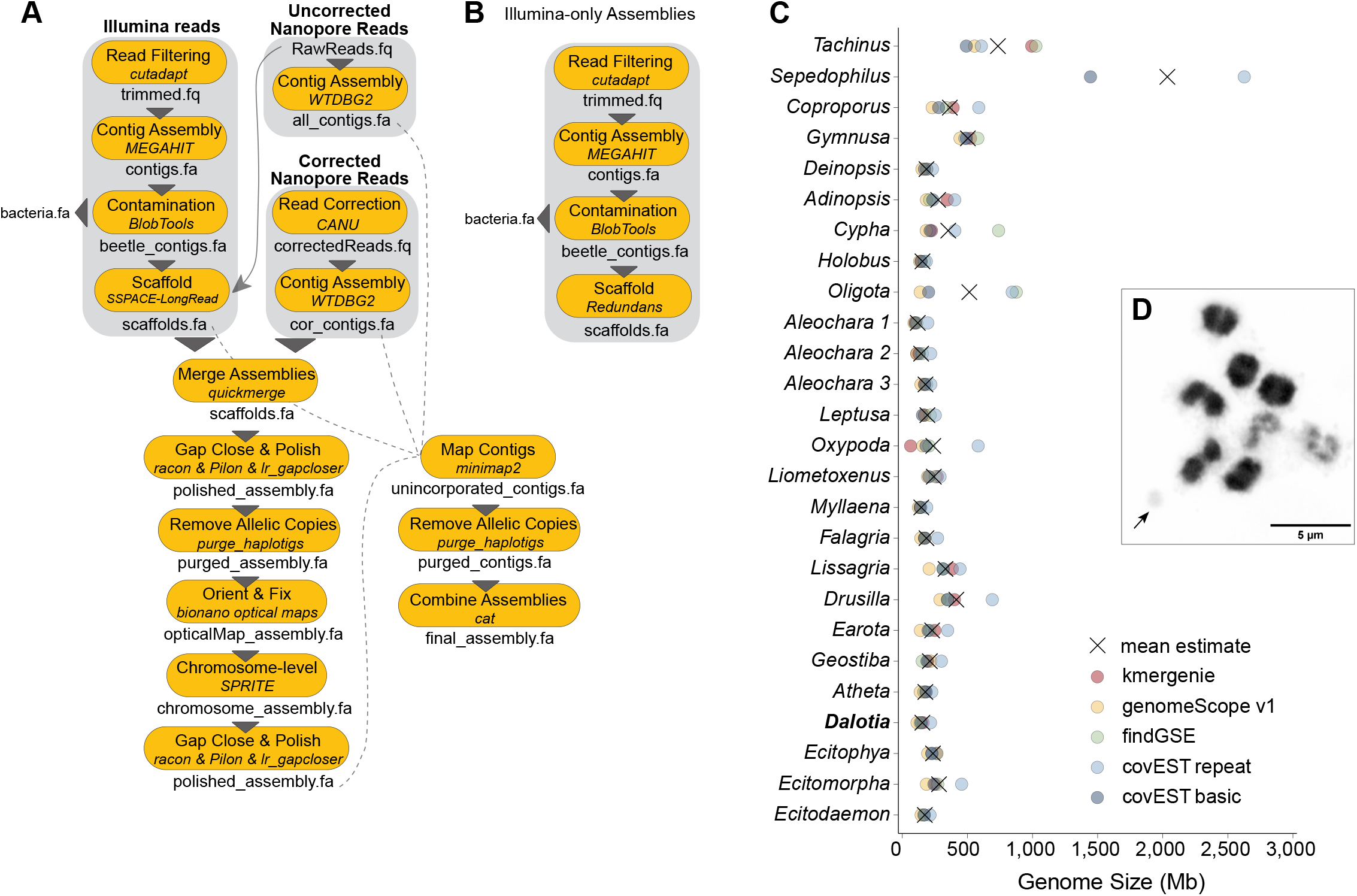
Genome assembly, sizes and karyotype. **(A)** A schematic of the different bioinformatic steps and data sets (Illumina short reads, Nanopore long reads, BioNano optical maps, and chromatin interaction reads via SPRITE) used to assemble the hybrid genome assembly of *Dalotia coriaria*. Contigs from the preliminary assemblies that did not map to the polished assembly were further filtered to remove putative haplotigs and then combined with the polished assembly for the final genome version (*Dcor* v3). **(B)** A schematic of the bioinformatic steps used to assemble the remaining genomes of the samples with only Illumina short-read data. **(C)** Estimates of genome size from five *k*-mer based tools (circles: red= kmergenie, yellow= genomeScope, green= findGSE, light blue= covEST repeat, and dark blue= covEST basic; X is the mean estimate). **(D)** Karyotype of *Dalotia* during mitosis (2n=9+Xyp). The arrow indicates the small Y chromosome.

**Supplemental Figure 2.**
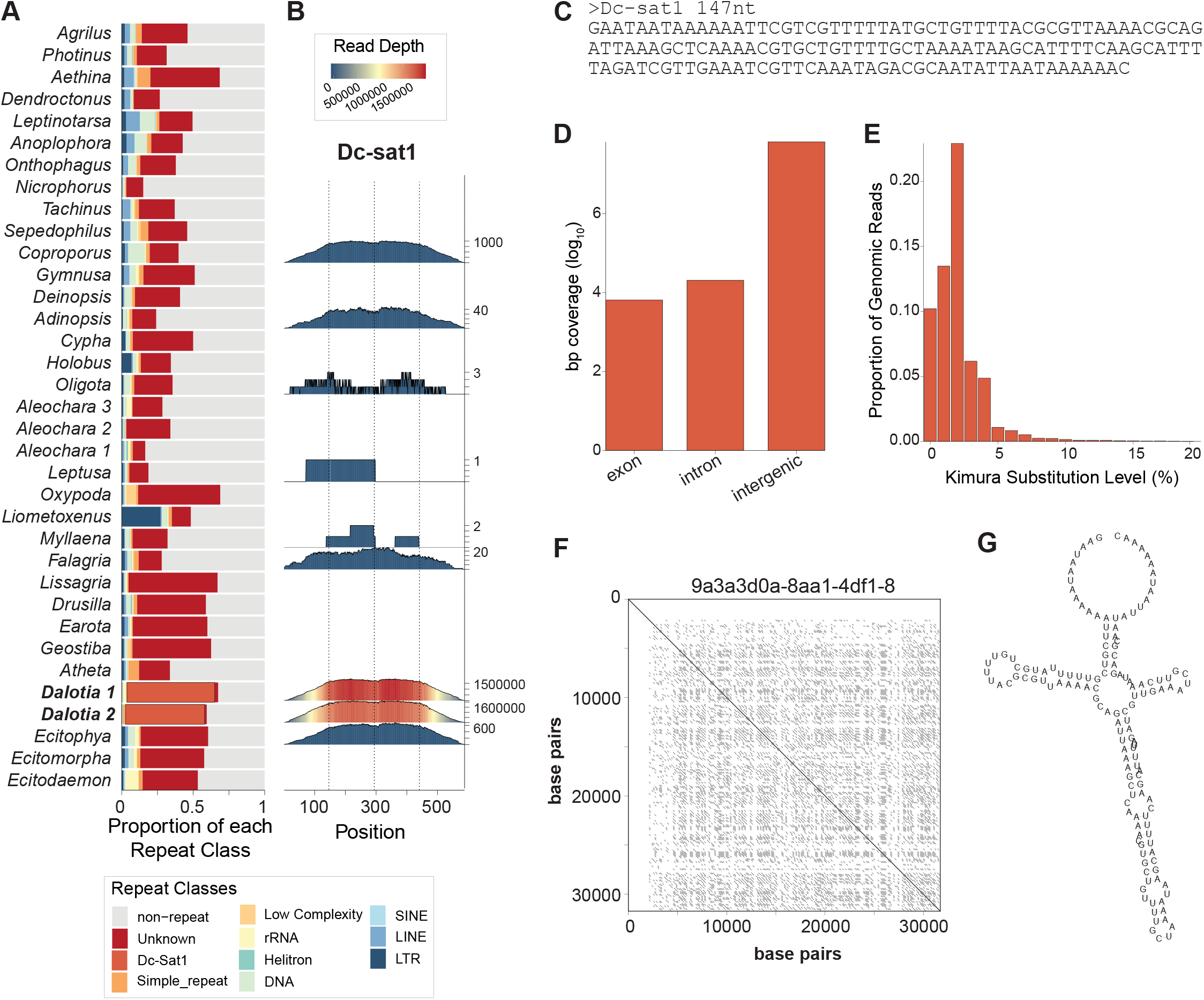
Repetitive content of the aleocharine genomes. **(A)** Predicted proportion of the genome composed of different repeat classes (LTR, LINE, SINE, DNA, Helitron, rRNA, low complexity, simple repeats, satellite Dc-Sat1, and unknown) and non-repetitive sequence from 0.25x subsampled short-reads for each respective species using dnaPipeTE v1.3.1. **(B)** Read depth across four concatenated copies of the 147 bp satellite Dc-Sat1 from subsampled short-reads from each respective species using RepeatProfiler v1.1. The y-axis was adjusted for each species based on maximum read depth. **(C)** The consensus sequence of Dc-Sat1 from RepeatExplorer2. **(D)** Genome coverage of Dc-Sat1 in the exons, introns and intergenic regions of *Dcor* v3 assembly. **(E)** Estimated proportion of *Dalotia* short-reads with different levels of Kimura substitution, a measure of sequence divergence over time, for Dc-Sat1. **(F)** Self dot-plot of one example minION read (9a3a3d0a-8aa1-4df1-8) with 35 tandem copies of Dc-Sat1. **(G)** Predicted secondary structure of Dc-Sat1.

**Supplemental Figure 3.**
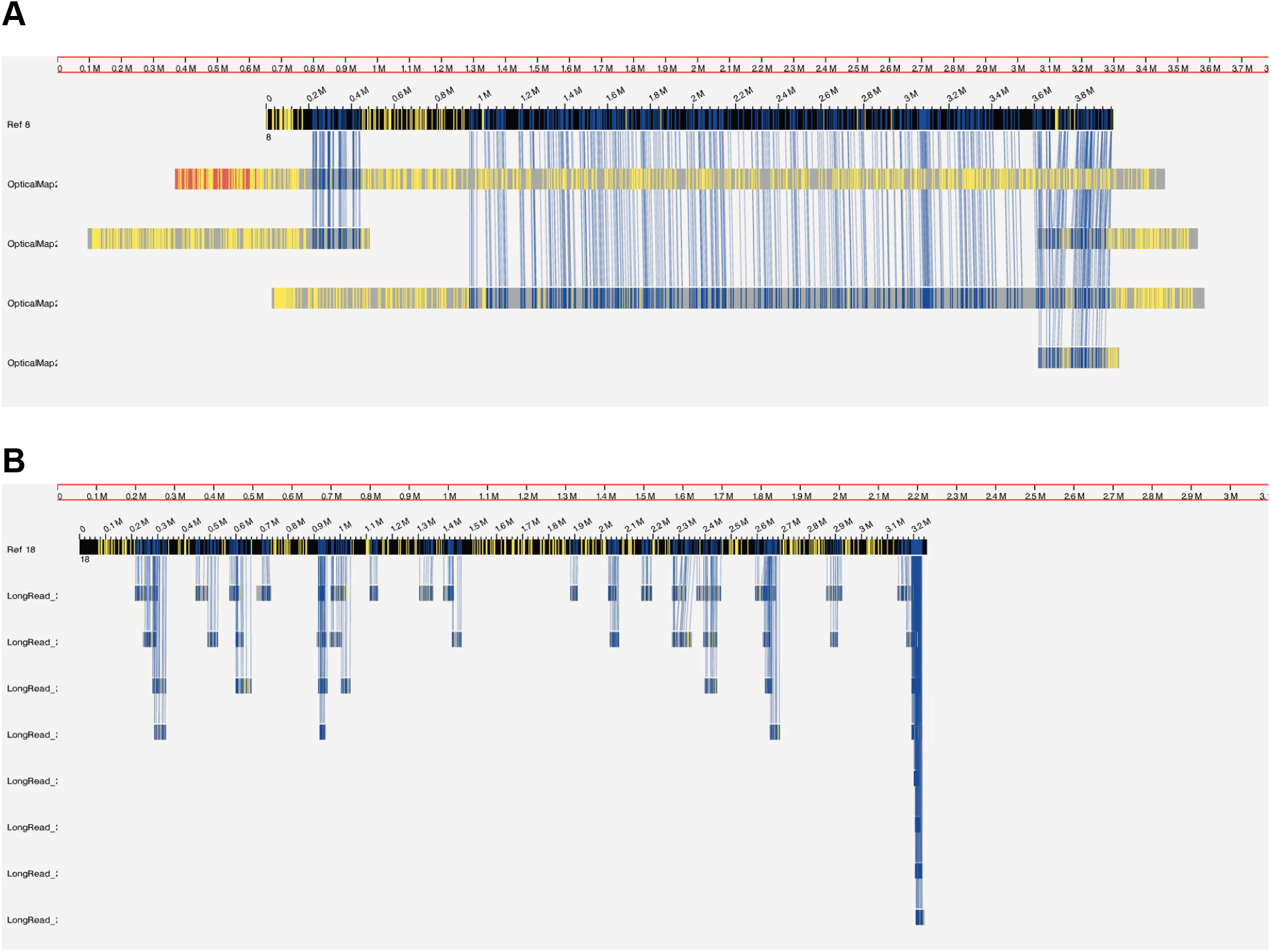
Visualization of Bionano optical map alignments against the *Dcor* v1 assembly and long-reads. **(A)** Five optical maps (13, 37, 95, 187, 243) aligned to scaffold ctg4 (Ref 8) from the *Dcor* v1 assembly. **(B)** Multiple minION long-reads mapped to optical map 18 (Ref 18) that was not captured by the hybrid assembly process combining optical maps with *Dcor* v1. Aligned labels are dark blue and unaligned labels are yellow along the reference sequence on top (background black) and corresponding query sequences below (background grey) in the genome browser of Bionano Access.

**Supplemental Figure 4.**
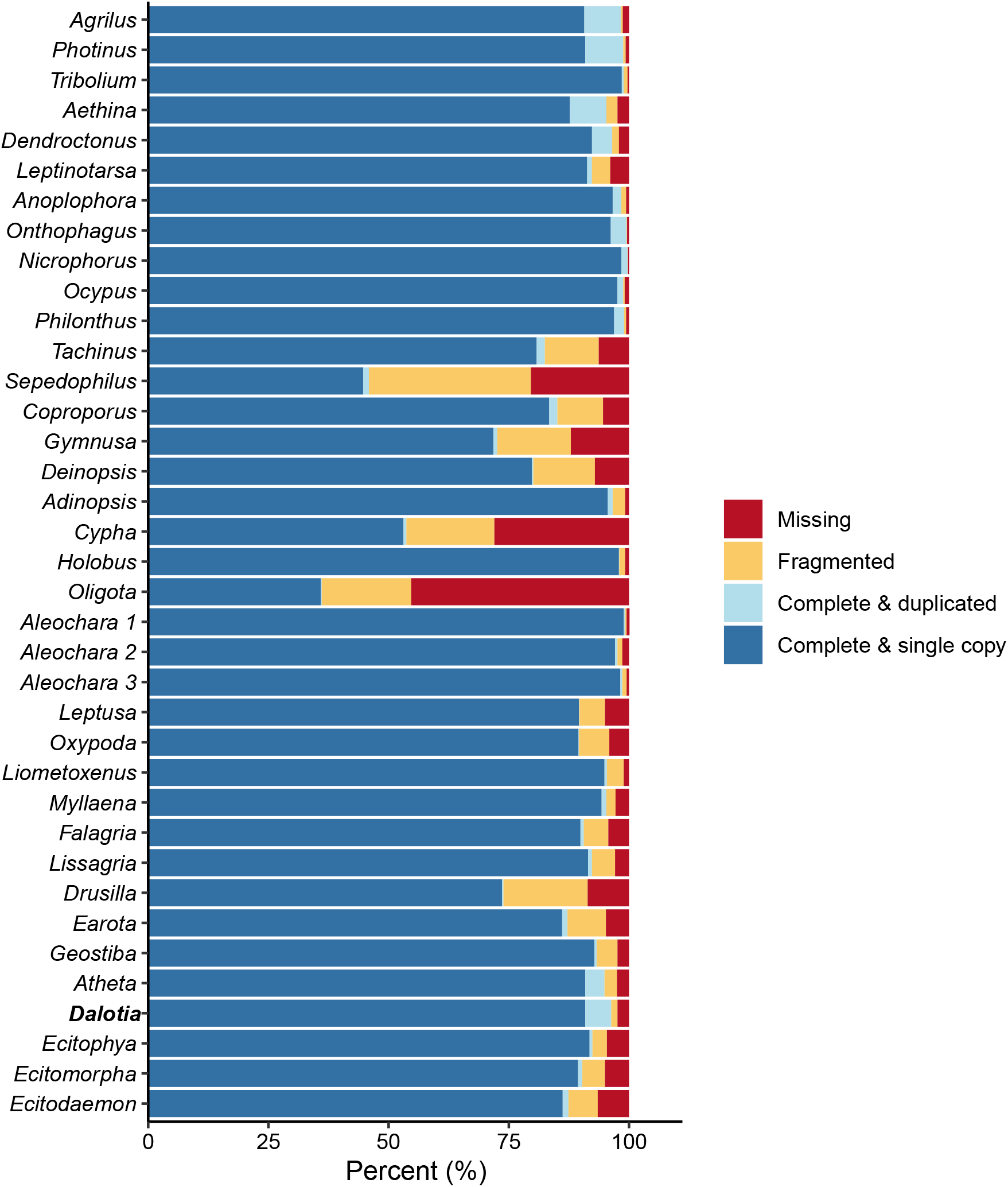
BUSCO genome completeness assessment for new and previously published beetle genome assemblies. Percentage of single-copy genes present in the genome assembly of each species using the arthropoda odb10 gene set (n=1013). Dark blue= complete and single copy, light blue= complete and duplicated, orange= fragmented or partial copy, red= missing orthologues.

**Supplemental Figure 5.**
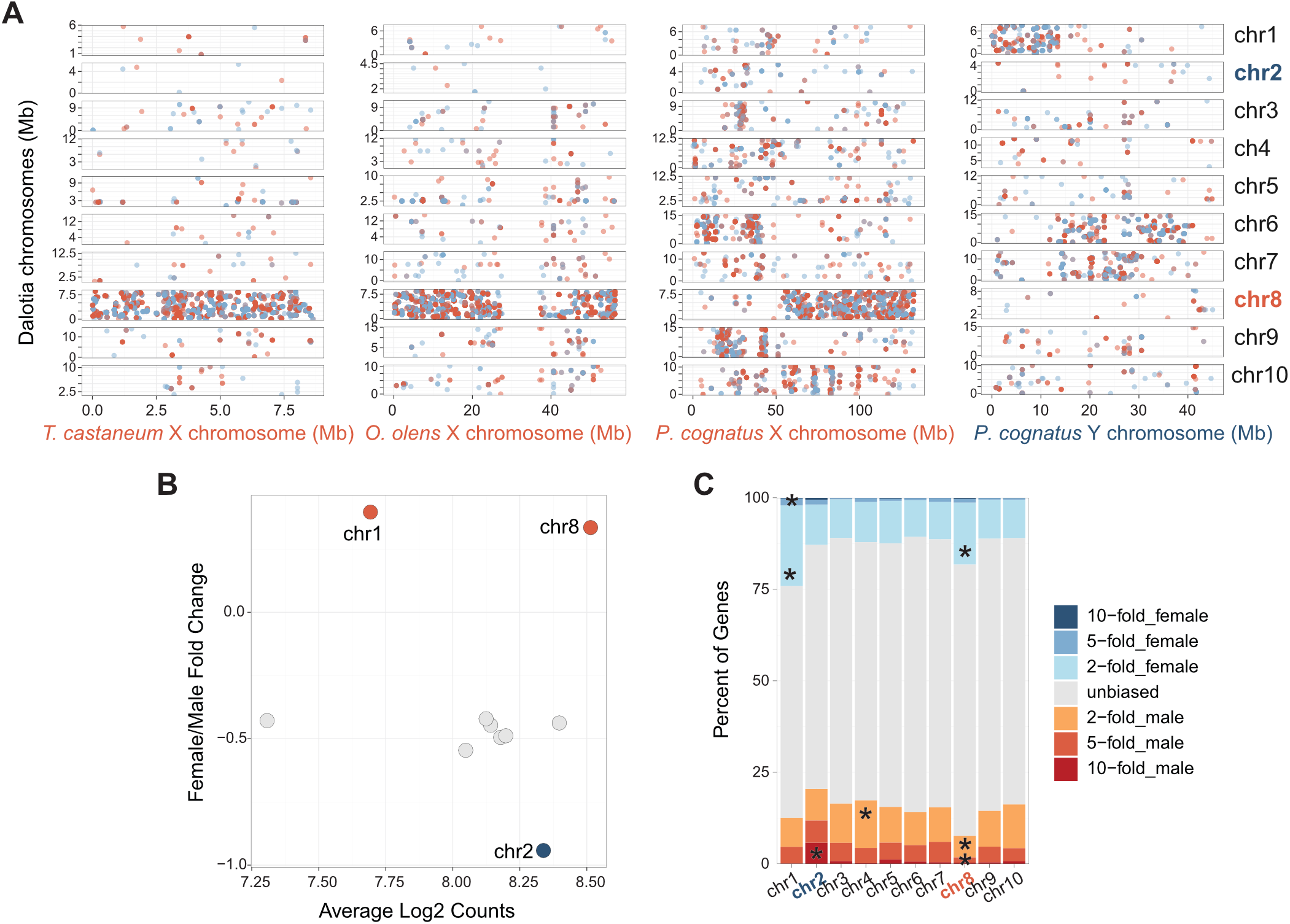
Sex chromosomes of *Dalotia* genome assembly. **(A)** PROmer amino acid sequence alignment of *Dalotia*’s ten chromosomes against sex chromosomes of *T. cadtaenum*, *O. olens*, and *P. cognatus*. Alignments were filtered to a minimum length of 200 aa. Each point represents an alignment with percent identity of 50% or higher and colored based on the strand, red is for the negative strand and blue is for the positive strand. **(B)** Summarized average log_2_ counts for all genes on a given chromosome for both sexes correlated to the fold-change in female to male expression for all genes for a given chromosome. Female-biased expression would have values greater than 0 whereas male-biased expression would be less than 0. **(C)** Genes with 2-, 5- and 10-fold difference in normalized log_2_ counts between the sexes were tabulated for each chromosome. Categories with significantly over or under representations of genes from Pearson’s Chi-square tests adjusted for multiple testing are indicated by asterisk.

**Supplemental Figure 6.**
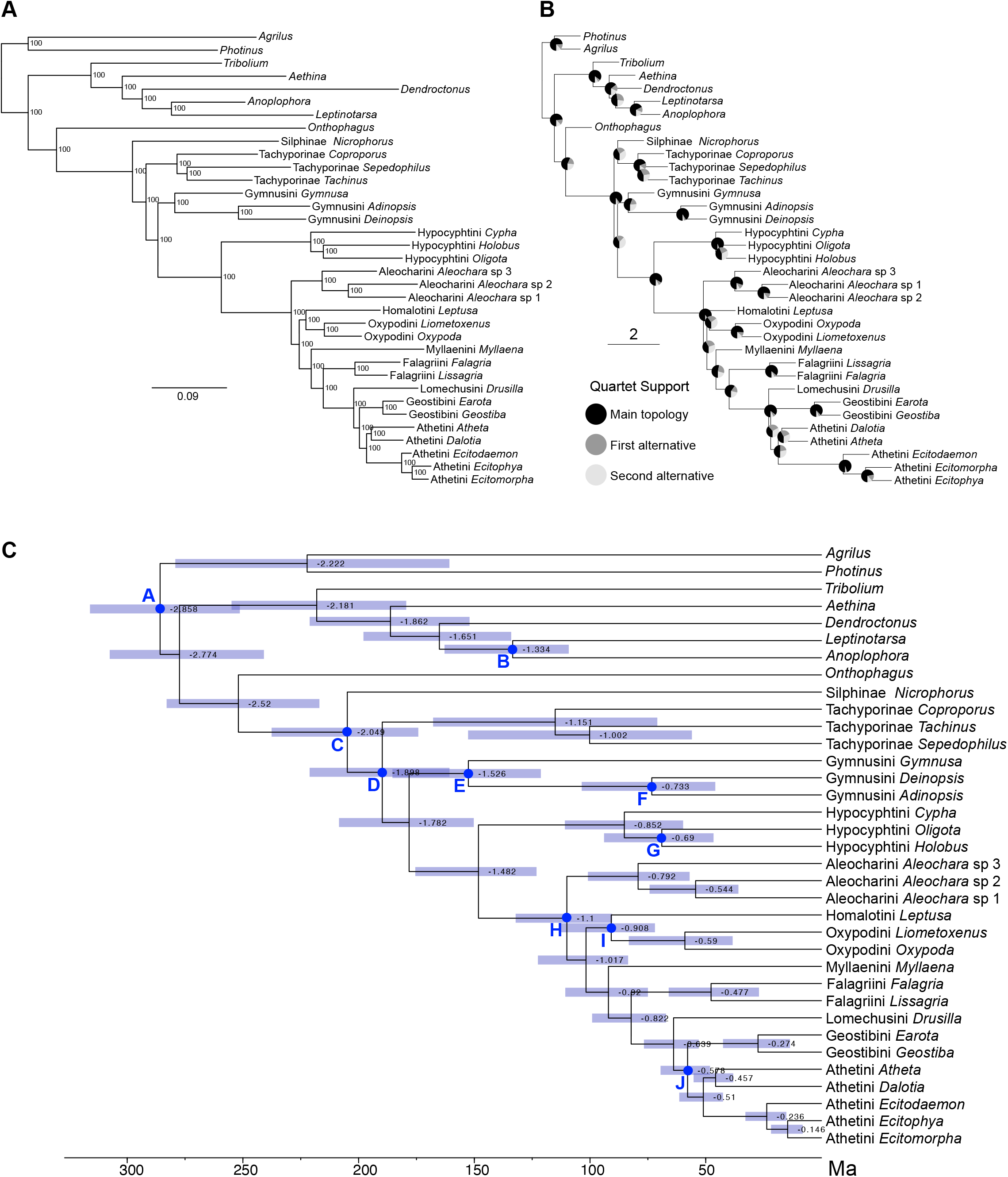
Phylogenomic relationships of Aleocharinae. **(A)** Maximum likelihood tree of 1,520 gene partitions (577,200 aligned amino acid sites) made with IQ-TREE v2.2.0-beta and 1,000 ultrafast bootstrap replicates. All nodes receive maximal bootstrap support. **(B)** Coalescent species tree made with ASTRAL v5.6.3. Quartet support is shown in pie charts at each node. **(C)** Dated phylogenomic tree made with MCMCtree and CODEML implemented in the PAML v4.9 package. Nodes used for fossil calibrations (A-J) are indicated (See **Table S4** for list of fossils and age bounds used to calibrate these nodes).

**Supplemental Figure 7.**
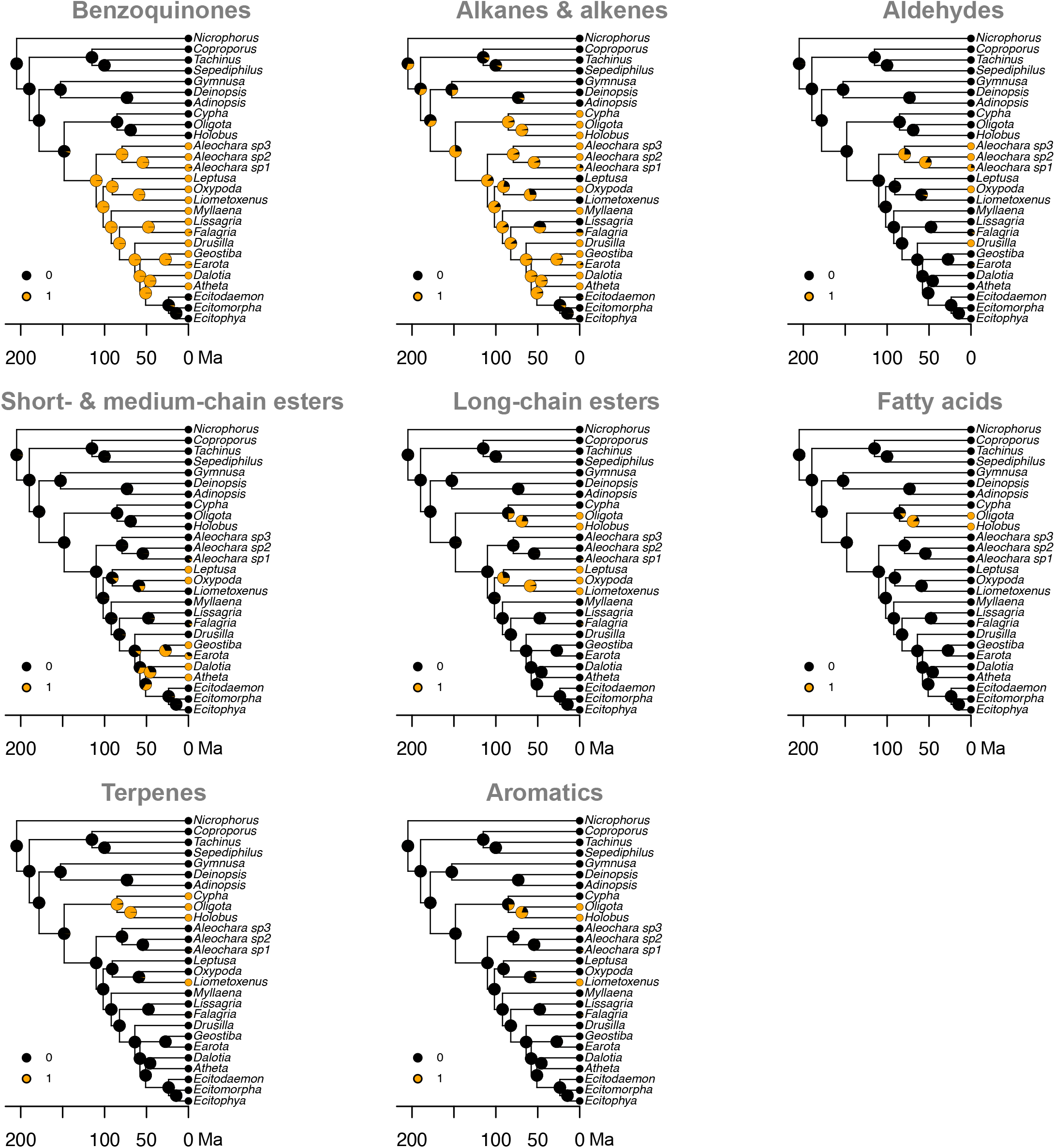
Ancestral state reconstruction of chemical classes found in the tergal gland. Pie charts at the nodes represent the maximum likelihood estimates of chemical class evolution along the dated species tree, starting at *Nicrophorus vespilloides*. Each chemical class was marked as present (1 = orange) or absent (0 = black) for extant species from the GC/MS data presented in Figure 3B. If no chemical data was available, we provided a probability of the chemical being absent as 0.5 in *Aleochara* sp1, *Falagria* and *Earota* and 0.9 in the ecitocharine clade based on morphology and chemical data from their closest sister taxon.

**Supplemental Figure 8.**
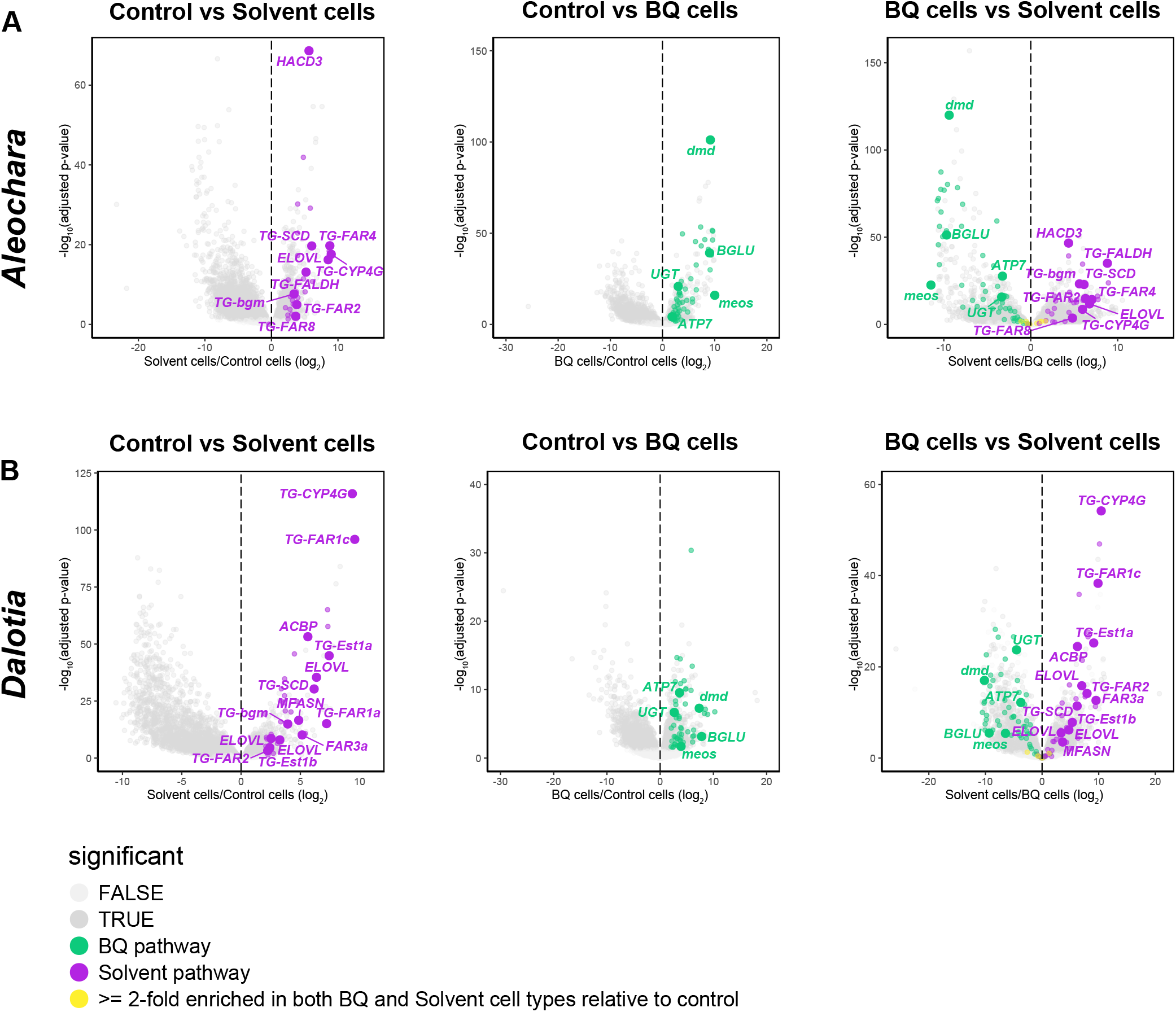
Pairwise comparisons of solvent cells, BQ cells and the control segment for *Aleochara* sp. 3 (A) and *Dalotia* (B). In each volcano plot the log_2_ fold-change between the pairwise comparison is plotted against the -log_10_ adjusted p-value. Differentially expressed genes (DEGs) that are characterized in either the solvent or BQ biosynthesis pathways are labelled and colored for each species. These DEGs may not be shared between the two species. Additional differentially expressed orthologs (DEOs) with 2-fold higher expression in each respective cell type relative to the control samples are colored but not labeled (magenta = solvent pathway, green = BQ pathway, yellow = both cell types, gland enriched). Other DEGs that are specific to each species are dark grey while those that are not significant are light grey for a given pairwise comparison.

**Supplemental Figure 9.**
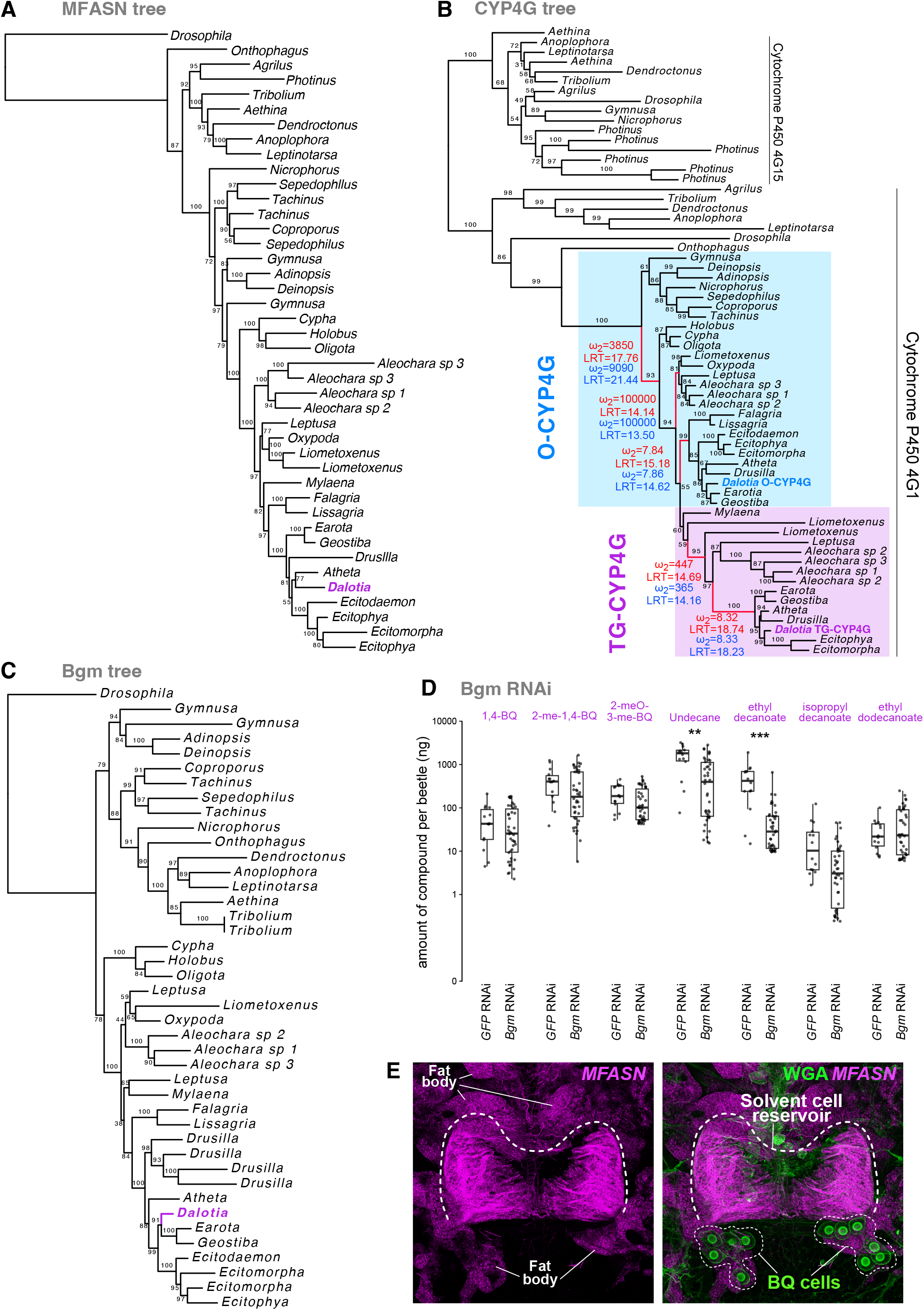
Evolution and function of solvent pathway enzymes. **(A-C)** Maximum likelihood trees of the enzymes Master Fatty Acid Synthase/MFASN (**A**; Q.insect+R5 model), Cytochrome P450 4G/CYP4G (**B**; Q.insect+R5) and Bubblegum/Bgm (**C**; LG+I+G4 model). Bootstrap support values are shown for each branch. *Dalotia* solvent pathway enzymes are highlighted in magenta. In **B**, colored branches show periods of episodic selection. aBSREL results from the all branches test are shown in red and on select branch test in blue. Associated omega (dN/dS) estimates with significant likelihood ratio test estimate (LRT) are presented for colored branches. The branch labeled for the CodeML results is indicated by a star. **(D)** RNAi silencing of the very long-chain-fatty-acid-CoA ligase *bgm* in *Dalotia* selectively diminishes the levels of undecane and ethyl decanoate. **(E)** HCR labeling of *MFASN* (magenta) in *Dalotia* reveals expression in solvent cells as well as fat body tissue distributed throughout the abdomen. Green: wheat germ agglutinin (WGA), which label the BQ cells.

**Supplemental Figure 10.**
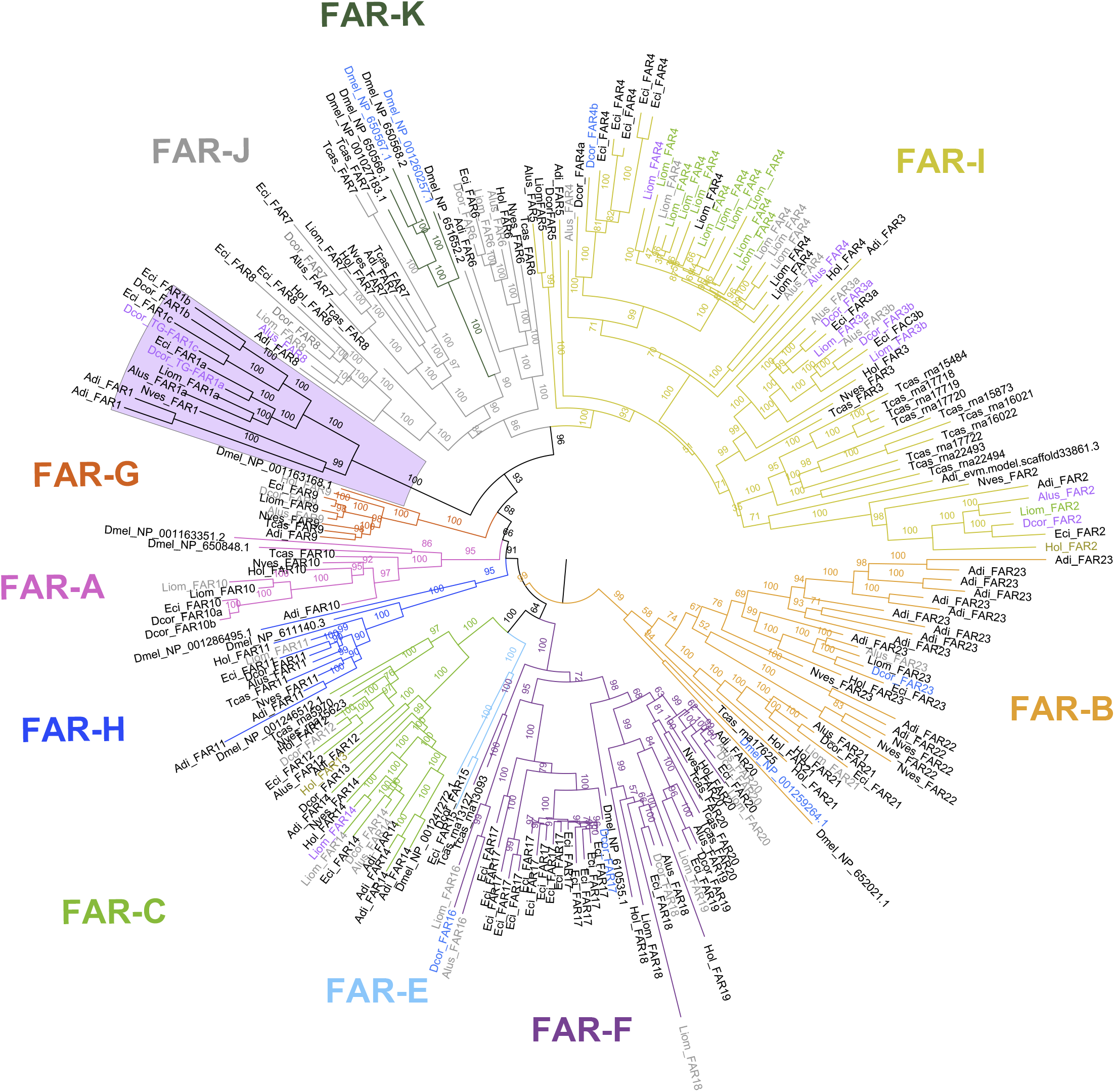
Fatty acyl-CoA reductase (FAR) tree. Maximum likelihood tree made with amino acid sequences using the LG+F+R8 model. Bootstrap support values are shown for each branch. FAR sequences were from selected Aleocharinae and outgroups: Dcor: *Dalotia coriaria*, Eci: *Ecitophya simulans*, Alus: *Aleochara* sp.3, Hol: *Holobus* sp., Liom: *Liometoxenus newtonarum*, Adi: *Adinopsis* sp., Tcas: *Tribolium castaneum*; Nves: *Nicrophorus vespilloides*; Dmel: *Drosophila melanogaster*. The magenta-boxed clade includes TG-FAR1c that functions in *Dalotia*’s solvent pathway. Classification of FARs followed the assigned groups designated by Tupec et al. (*185*).

**Supplemental Figure 11.**
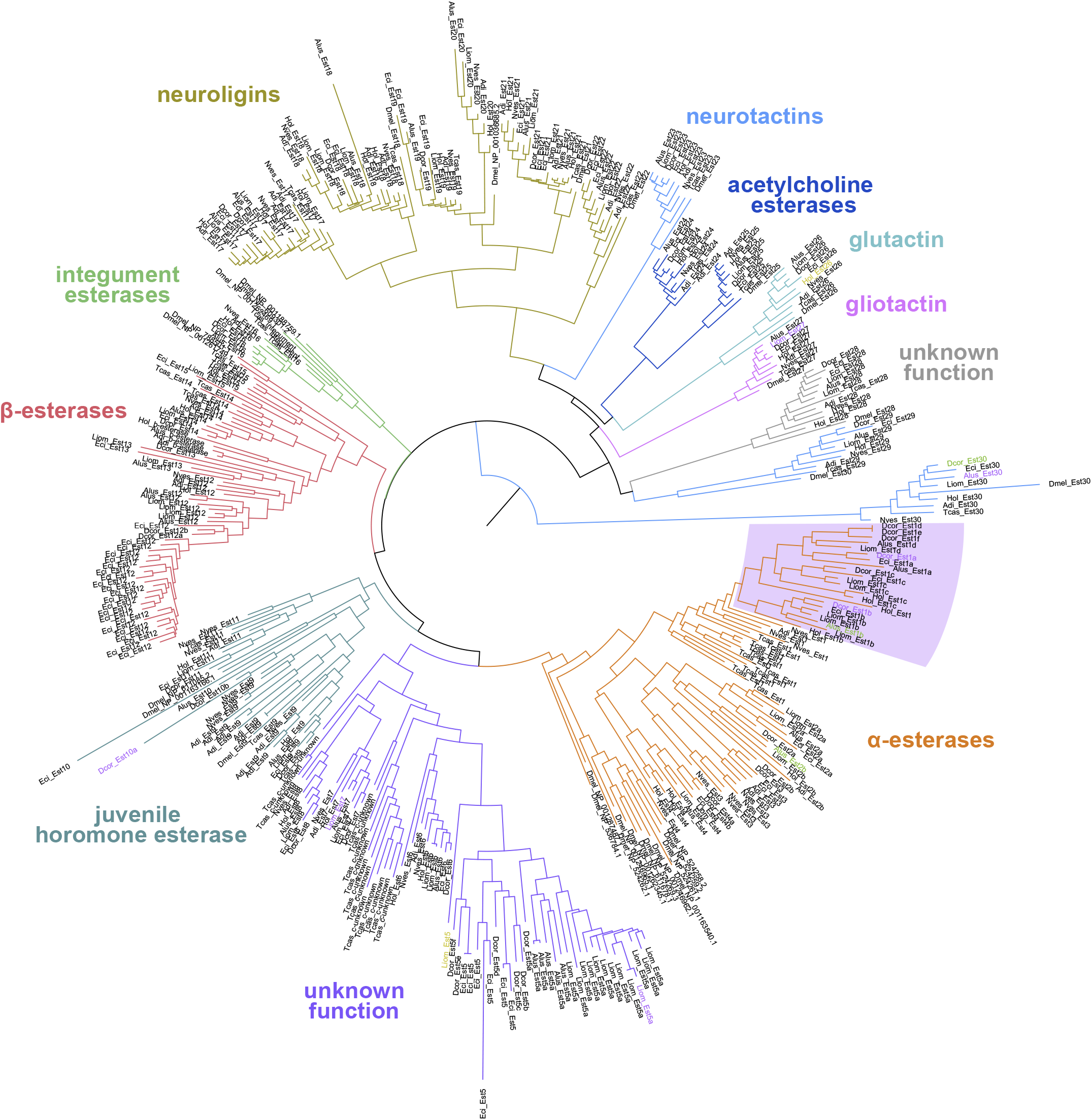
Esterase tree. Maximum likelihood tree of carboxylesterase enzyme family made with amino acid sequences using the LG+R10 model. Bootstrap support values are shown for each branch. Esterase sequences were from selected Aleocharinae and outgroups: Dcor: *Dalotia coriaria*, Eci: *Ecitophya simulans*, Alus: *Aleochara* sp.3, Hol: *Holobus* sp., Liom: *Liometoxenus newtonarum*, Adi: *Adinopsis* sp., Tcas: *Tribolium castaneum*; Nves: *Nicrophorus vespilloides*; Dmel: *Drosophila melanogaster*. The magenta-boxed clade includes TG-Est1a that functions in *Dalotia*’s solvent pathway, and TG-Est1b that is co-expressed in solvent cells. Classification of esterases followed the assigned groups designated by Oakeshott et al. (*186*).

**Supplemental Figure 12.**
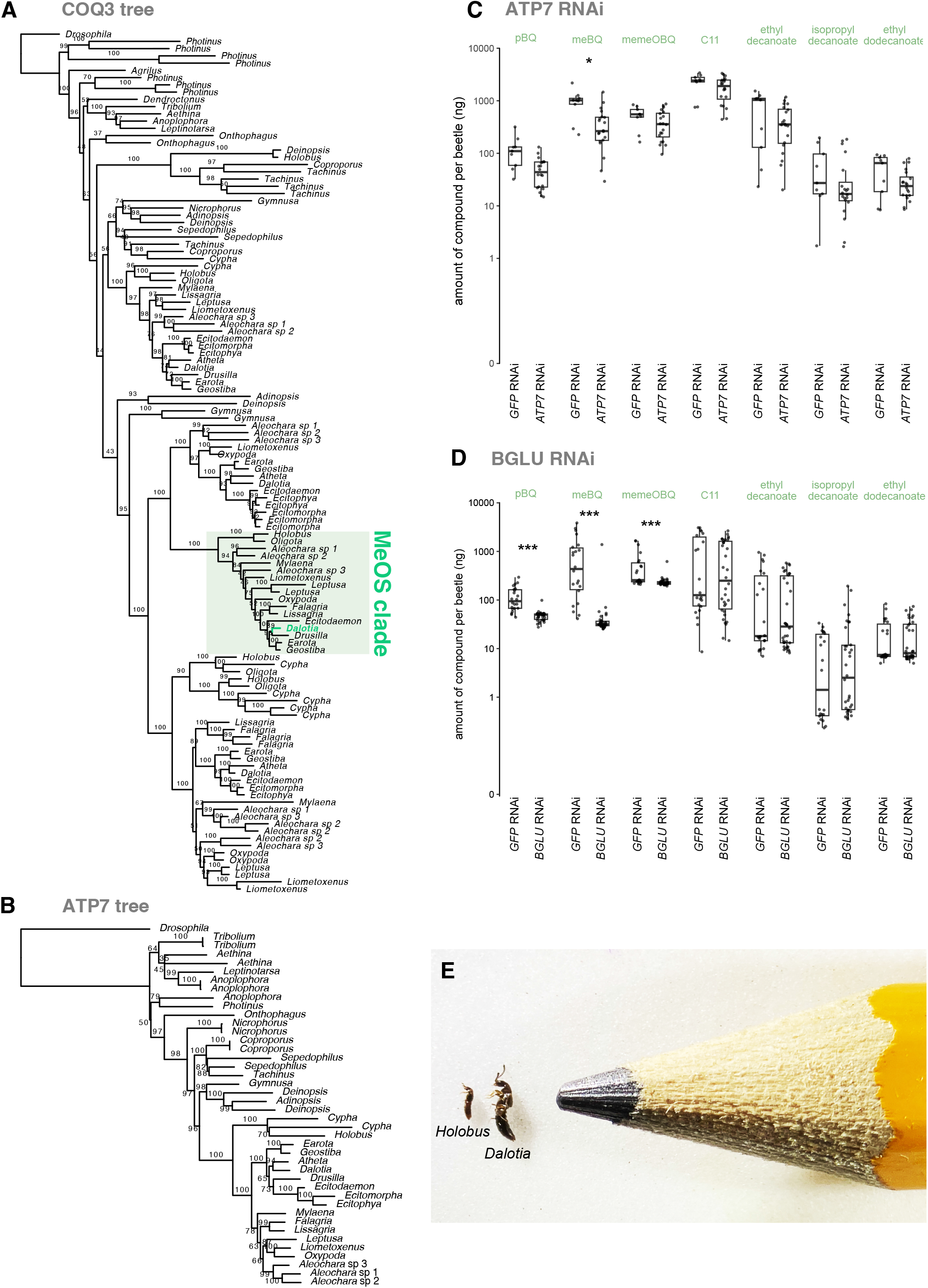
Evolution and function of BQ pathway enzymes. **(A)** Maximum likelihood tree of methyoxyless/MeOs using the LG+R6 model. The MeOs clade is highlighted in the green box. **(B)** Maximum likelihood tree of copper-transporting ATPase 1/ ATP7 using Q.insect+R5 model. For each tree, bootstrap support values are shown for each branch. *Dalotia* BQ pathway enzymes are highlighted in green. **(C-D)** RNAi silencing of the *ATP7* **(C)** and *β-glucosidase* **(D)** in *Dalotia* selectively diminishes the levels of benzoquinones. **(E)** Photograph of *Holobus*, on the left, next to *Dalotia*, in the center, and a standard size pencil on the right.

**Supplemental Figure 13.**
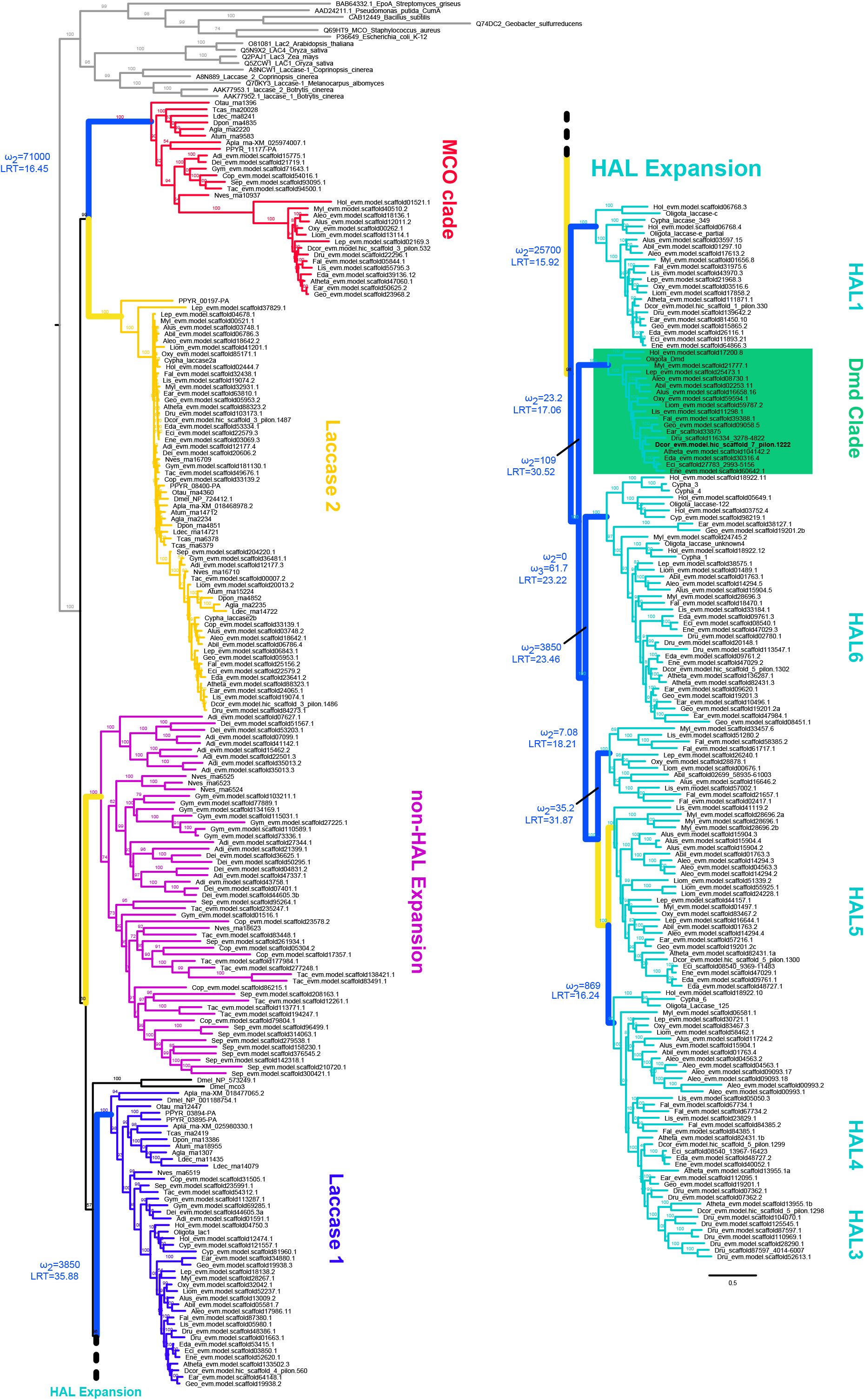
Laccase tree. Complete maximum likelihood tree of the laccase gene family made with amino acid substitution model LG+R10. Bootstrap support values are shown for each branch. The d*md* clade is highlighted in green. aBSREL significant results for select branches are shown in blue and non-significant, selected branches in yellow. Associated omega (dN/dS) estimates with significant likelihood ratio test estimate (LRT) are presented for blue branches. All terminal branches leading to *Liometoxenus* and ecitocharine clade were also tested but their significance is not labeled. Abbreviations of species are as follows: Abil= *Aleochara bilineata* (sp1), Adi = *Adinopsis* sp., Agla = *Anoplophora glabripennis*, Aleo= *Aleochara nigra* (sp2), Alus= *Aleochara* sp.3, Apla= *Agrilus planipennis*, Atheta = *Atheta pasadenae*, Atum= *Aethina tumida*, Cop= *Coproporus ventriculus*, Cypha= *Cypha longicornis*, Dcor= *Dalotia coriaria*, Dei= *Deinopsis earosa*, Dmel= *Drosophila melanogaster*, Dpon= *Dendroctonus ponderosae*, Dru= *Drusilla canaliculata*, Ear= *Earota dentata*, Eci= *Ecitophya simulans*, Eda= *Ecitodaemon* sp., Ene= *Ecitomorpha nevermanni*, Fal= *Falagria* sp., Geo= *Geostiba* sp., Gym= *Gymnusa* sp., Hol= *Holobus* sp., Lep= *Leptusa* sp., Liom= *Liometoxenus newtonarum*, Lis= *Lissagria laeviuscula* Ldec = *Leptinotarsa decemlineata*, Myl= *Myllaena* sp., Nves= *Nicrophorus vespilloides*, Oxy= *Oxypoda opaca*, Oli = *Oligota* sp., Otau= *Onthophagus taurus*, PPYR= *Photinus pyralis*, Sep= *Sepedophilus* sp., Tac= *Tachinus* sp., Tcas= *Tribolium castaneum*,.

**Supplemental Figure 14.**
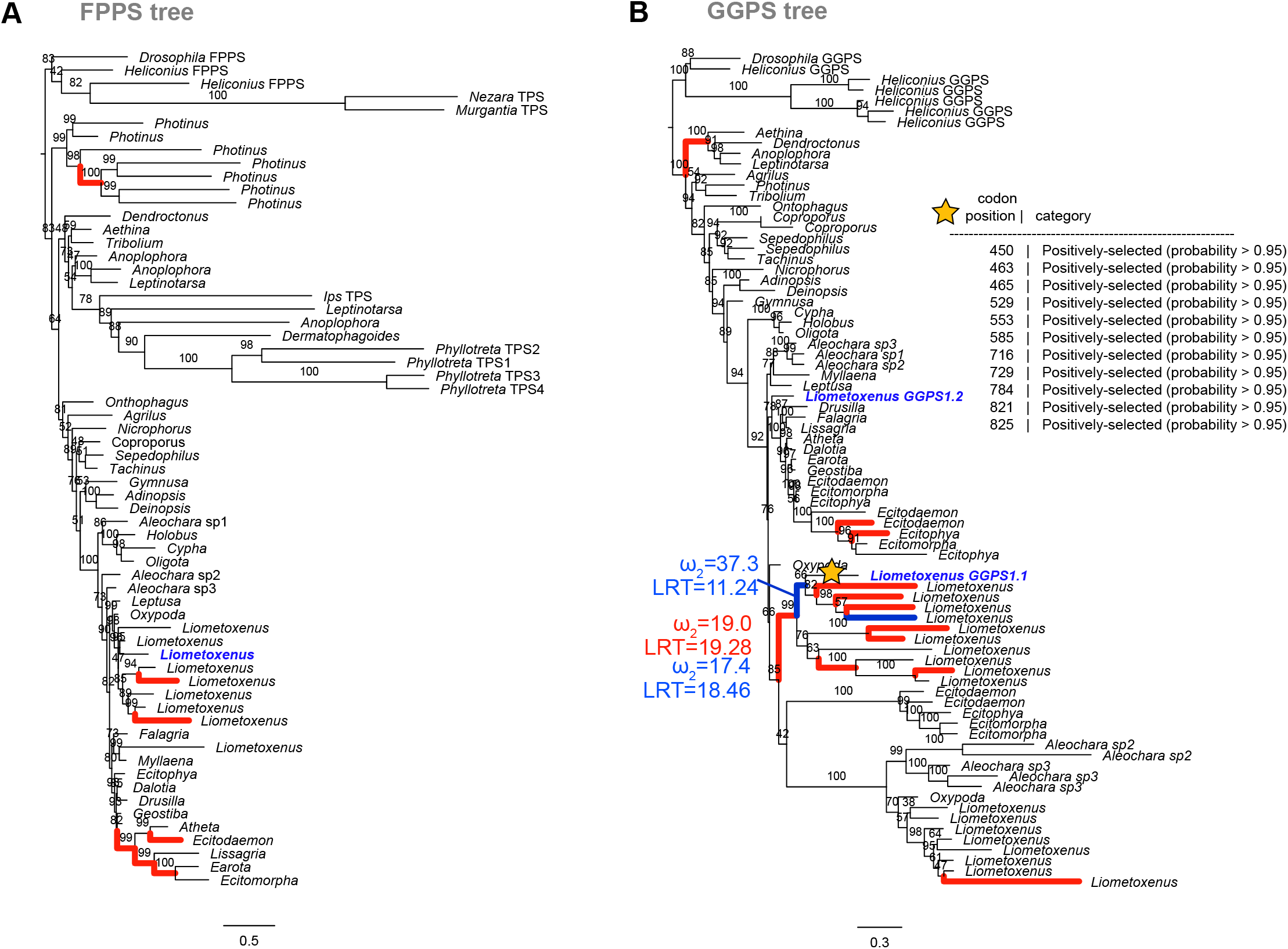
Putative terpene synthase enzymes in *Liometoxenus*. **(A)** Maximum likelihood tree of farnesyl pyrophosphate synthase/FPPS using the Q.insect+R5 model. **(B)** Maximum likelihood tree of geranylgeranyl pyrophosphate synthase/GGPS using the JTT+F+I+G4 model. Bootstrap support values are shown for each branch for each tree. In both trees, colored branches show periods of episodic selection. Results of the all branches test are shown in red and on select branch test in blue with the associated omega (dN/dS) estimates and likelihood ratio test estimate (LRT) for branches leading to *Liometoxenus* genes upregulated in BQ cells (blue). The branch labeled with a star was tested with CodeMLbranch-site model. Significant amino acid positions under selection based on Bayes Empirical Bayes analysis are presented in panel B inset table.

**Supplemental Figure 15.**
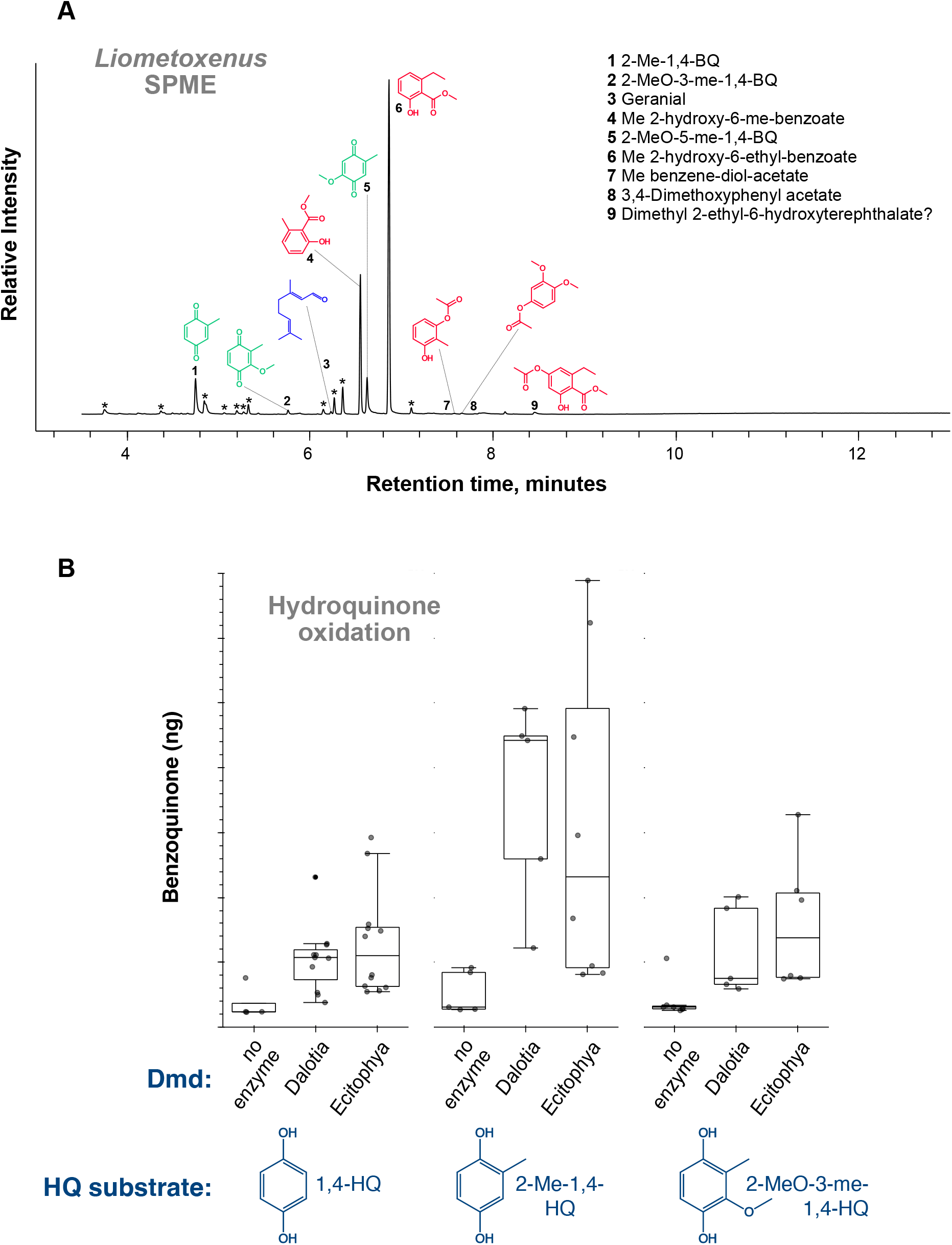
Chemical secretion and enzymatic activity of myrmecophilous aleocharines. **(A)** Volatilized chemicals from *Liometoxenus* glandular excretion. Headspace volatiles from a single *Liometoxenus* beetle detected via single-phase microextraction (SPME). **(B)** Enzyme activity of Dmd from *Ecitophya*. Synthesized Dmd of *Ecitophya* can convert hydroquinone substrates (HQs) to the corresponding benzoquinones at comparable efficiency to *Dalotia* Dmd.

**Supplemental Video 1. *Liometoxenus*-host ant interaction.** Two *Liometoxenus newtonarum* beetles flexing their abdomens to deploy tergal gland volatiles. On encountering the beetles, a *Liometopum occidentale* worker ant does not attack them but instead begins to self-groom before experiencing impaired locomotion and appearing to "tremble".

